# Detection of potential new SARS-CoV-2 Gamma-related lineage in Tocantins shows the spread and ongoing evolution of P.1 in Brazil

**DOI:** 10.1101/2021.06.30.450617

**Authors:** U.J.B. Souza, R.N. Santos, A. Belmok, F.L. Melo, J.D. Galvão, S.B. Damasceno, T.C.V. Rezende, M.S. Andrade, B.M. Ribeiro, J.C. Ribeiro Junior, R.F. Carvalho, I.G.C. Santos, M.S. Oliveira, F.R. Spilki, F.S. Campos

## Abstract

After more than a year of the pandemic situation of COVID-19, the United Kingdom (UK), South Africa, and Brazil became the epicenter of new lineages of severe acute respiratory syndrome coronavirus 2 (SARS-CoV-2). Variants of Concern (VOCs) were identified through a continuous genomic surveillance global effort, the B.1.1.7 (Alpha), B.1.351 (Beta), B.1.617.2 (Delta), and P.1 (Gamma) harboring a constellation set of mutations. This research aims to: (i) report the predominance of the Gamma (P.1) lineage presenting the epidemiological situation of the SARS-CoV-2 genomic surveillance at the state of Tocantins, and (ii) describe the emergence of possible new mutations and viral variants with the potential new lineage (P1-related) represented by 8 genomes from the Tocantins harboring the mutation L106F in ORF3a. At the moment, 6,687 SARS-CoV-2 genomes from GISAID carry this mutation. The whole-genome sequencing has an important role in understanding the evolution and genomic diversity of SARS-CoV-2, thus, the continuous monitoring will help in the control measures and restrictions imposed by the secretary of health of the state to prevent the spread of variants.

## Introduction

Since 2019, the host spillover of the severe acute respiratory syndrome coronavirus 2 (SARS-CoV-2) [1], caused the biggest pandemic of the 21st century with severe impacts on health, economy and social life. To understand the SARS-CoV-2 variants the genomic surveillance tool was applied to track cases providing important clues to the complex virus-host relationship [2,3]. In Brazil specially, the Corona-ômica-BR is conducting efforts and human resources to track the viral spread and contribute to public health authorities. The emergence of viruses may be associated with a lack of public interventions, non-scientific communications, social inequality and for now, there is a slow vaccination [4,5].

The international public efforts have been concerned about the emergence of virus variants leading to changes in viral fitness. These mutations may increase transmissibility, enhance escape from the human immune response, or otherwise alter biologically important phenotypes [6]. The global emergence of several variants of concern (VOCs), recently re-named by WHO, including the UK variant (Alpha or B.1.1.7), the South Africa variant (Beta or B.1.351), Brazil variant (Gamma or P.1 and P.2), and Delta or B.1.617.2 variant in India lead the genomic surveillance essential to monitoring mutations [7]. We hereby report the complete genome sequences and phylogenetic analysis of 24 SARS-CoV-2 genomes from Tocantins, North region of Brazil including our P.1-related sequences.

## Results and discussion

Tocantins is a state in Brazil located in the north of the country. It is a region with intense traffic of people, connecting the north, northeast and central-west regions of Brazil between frontiers with more six states: Maranhão, Piauí, Bahia, Goiás, Mato Grosso and Pará. Thus, the daily/seasonal travelling dynamics favors the introduction and spread of new SARS-CoV-2 lineages in the state. We analyzed in both spatio-temporal and phylogenetic contexts 24 SARS-CoV-2 samples collected between May 26, 2021 and June 01, 2021 from 12 municipalities of the state of Tocantins. Previous SARS-CoV-2 genomes available in GISAID [8–10] indicated that the P.1 variant that emerged during November 2020 has rapidly and dominantly increased its prevalence in Brazil. In Tocantins state, P.1 was first detected in February, 2021. Since then, there was a fast replacement by P.1 lineage, the latter being present in 75% of the sampling sequenced for this report.

Overall, 58 genomes (24 from this study and 34 deposited in GISAID) from 21 municipalities of the state of Tocantins state have been sequenced (**Figure 1**). The Genomic analysis reveals six SARS-CoV-2 lineages circulating with a high prevalence of P.1 (Gamma) lineage (63.8% or 37 genomes), followed by B.1.1.28 (19.0% or 11 genomes), P.2 (Zeta) (10.3% or six genomes), B.1.1.33 (3.4% or two genomes), B.1.1 (1.7% or one genome) and B.1.1.401 (1.7% or 1 genome).

**Figure 1.**
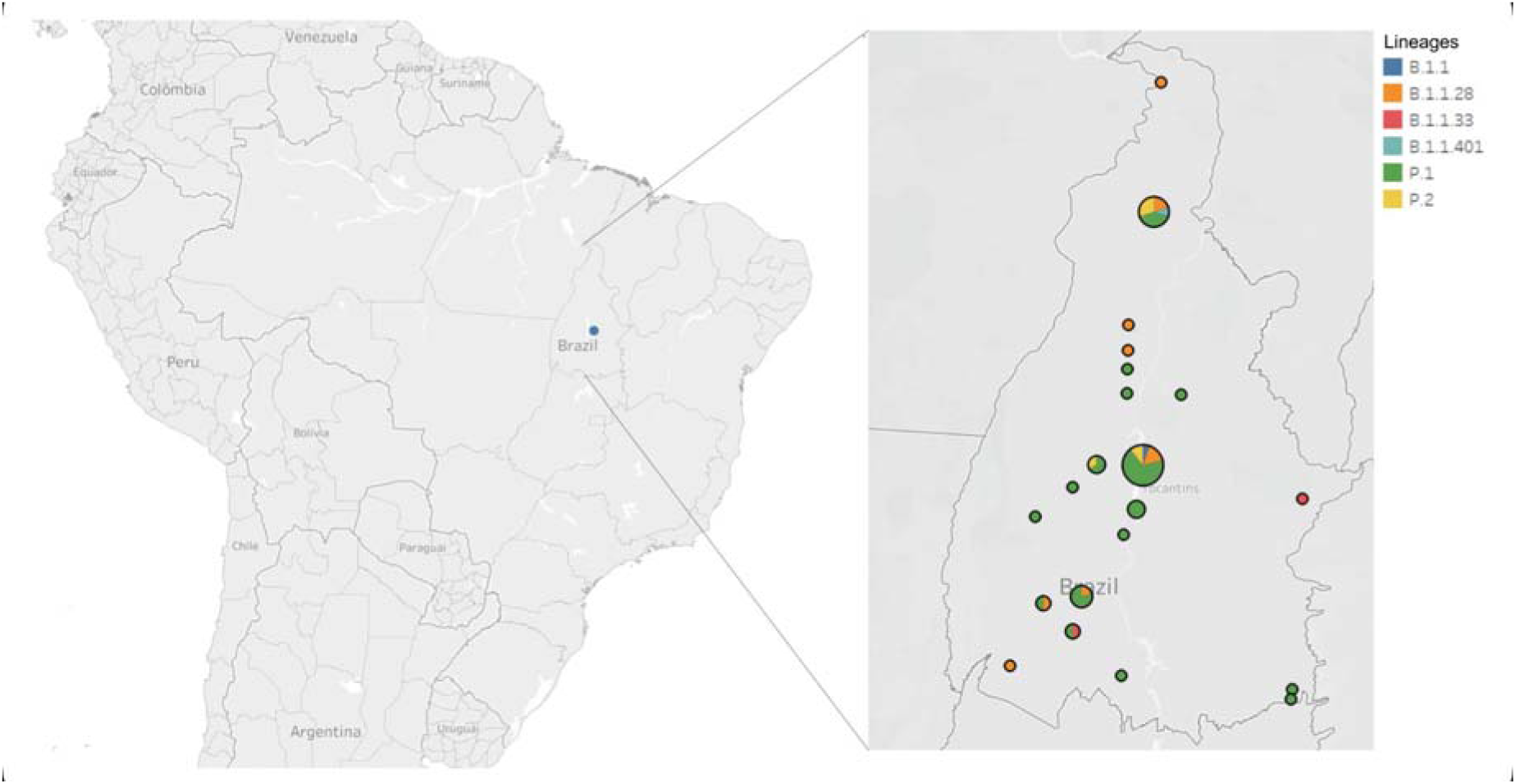
A) State map of Brazil with emphasis on the state of Tocantins. The color shows details about the lineages circulating in Tocantins. Size shows the sum of number of whole genomes sequenced. Araguaína (B.1.1.28, B.1.1.401, P.1 and P.2), Araguatins (B.1.1.28), Brejinho de Nazaré (P.1), Combinado (P.1), Figueirópolis (P.1), Formoso do Araguaia (B.1.1.28 and P.1), Fortaleza do Tabocão (P.1), Guaraí (B.1.1.28), Gurupi (B.1.1.28 and P.1), Jaú do Tocantins (P.1), Lagoa da Confusão (P.1), Mateiros (B.1.1.33), Novo Alegre (P.1), Palmas (capital of the state - B.1.1, B.1.1.28, P.1 and P.2), Paraíso do Tocantins (P.1 and P.2), Pium (P.1), Porto Nacional (P.1), Presidente Kennedy (B.1.1.28), Rio dos Bois (P.1), Rio Sono (P.1) and Sandolândia (B.1.1.28).

The variant P.1 of the SARS-CoV-2 virus has 17 mutations, including three (K417T, E484K, and N501Y) in the spike protein which may exhibit increased transmissibility and/or immune evasion [5,11]. Currently, P.1 (Gamma) is the dominant lineage widespread across all Brazilian regions corresponding for 62.6% (10,429 genomes) of the complete and high coverage genomes on GISAID (A total of 16,663 complete and high coverage genomes. June 28, 2021). Figure 2 shows the six prevalent lineages in Brazil from April 2020 to June, 2021. Six lineages account for 94.3% (15,721 genomes sequenced). The P.2 (Zeta) lineage (1,981 genomes) is the second most prevalent, followed by B. 1.1.28 (1,322 genomes), B.1.1.33 (1,266 genomes), B.1.1.7 (410 genomes) and P.1.2 (313 genomes).

**Figure 2.**
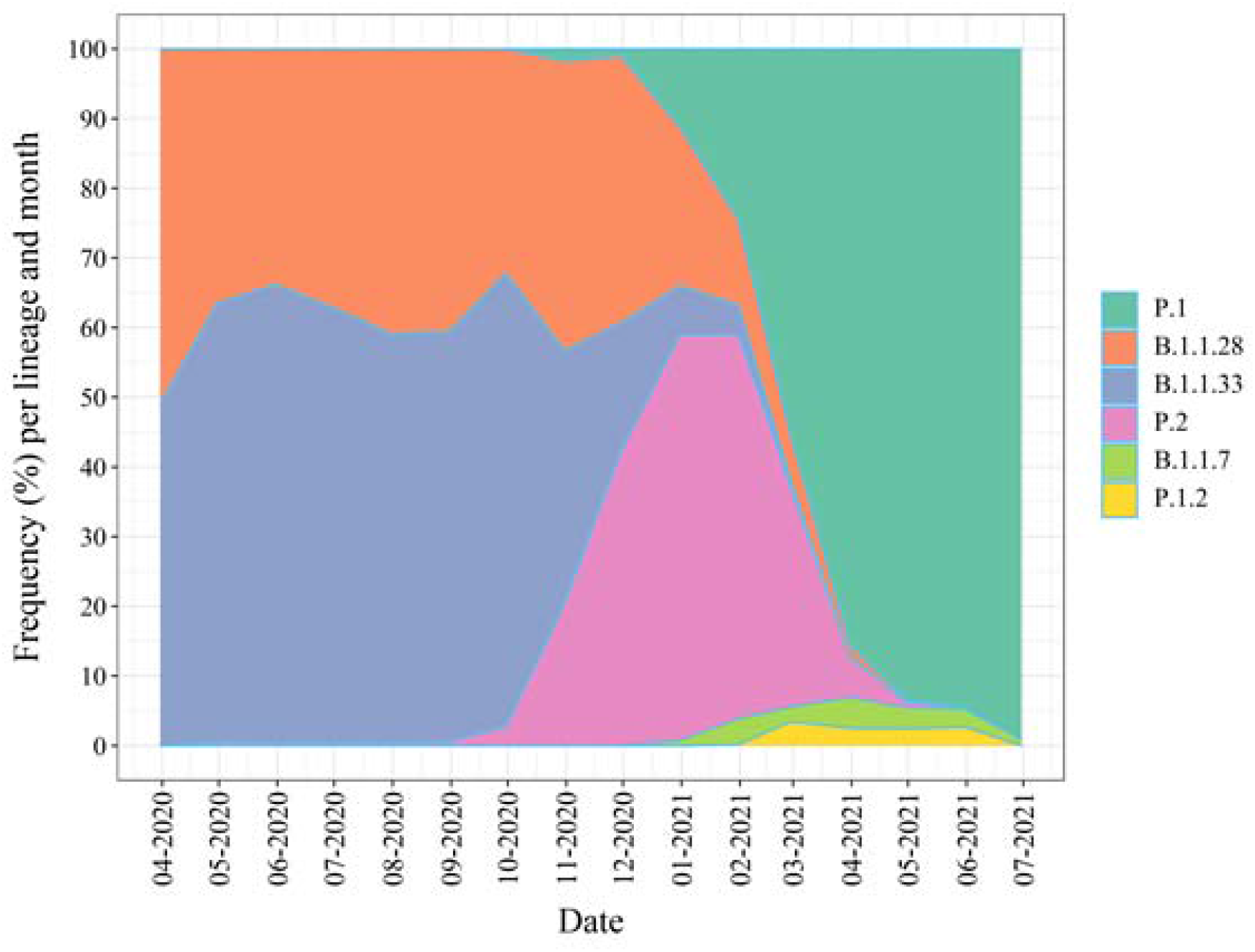
Frequency of the most abundant lineages across Brazil within the last 6 months.

The cluster previously identified in Tocantins (northern region of Brazil) brings together 91 Gamma genomes, the majority from São Paulo (86.8% or 79 genomes) and Goiás (6.6% or 6 genomes) and all harbor the L106F mutation in ORF3a (**Figure 3**). To date there are 6,687 SARS-CoV-2 genomes worldwide sharing this mutation in the GISAID database. The first detections were carried out in March 2020 in North America and Europe. Until the end of December, 1,298 (19.4%) genomes had this mutation, with the remainder, 5,389 (80.6%), detected in 2021 (with a predominance in April of 1,625 genomes). Europe corresponds to 72.2% (4,829) of the detected genomes and North America 17% (1,140). South America represents 3.3% (219) in which Brazil corresponds to 2% (131) genomes containing this mutation. These findings reinforce the importance of genomic surveillance and accurate attention in deposited data.

**Figure 3.**
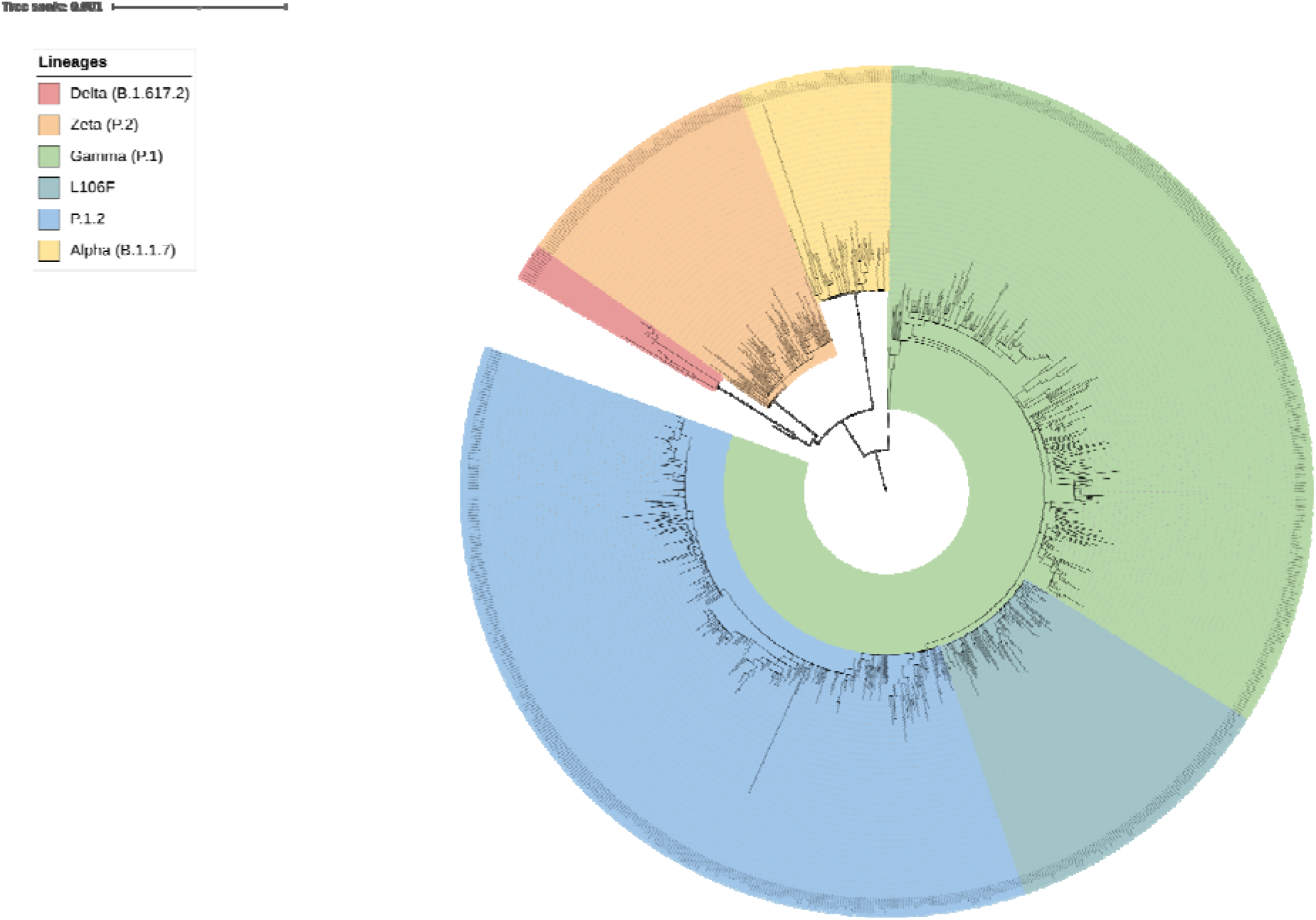
Phylogenetic tree of 851 genomes retrieved from GISAID and the 24 genomes newly sequenced on Tocantins state. Colored clade indicates lineage Delta (B.1.617.2) in red, P.2 (Zeta) in orange, B.1.1.7 (Alpha) in yellow, P.1 (Gamma) in green, P.1.2 in blue and the newly lineage P.1 (Gamma)-related, named here L106F, in dark blue. The maximum likelihood tree was built with IQ-TREE2.

Recently, new mutations were observed in the P.1 variant, giving rise to a new P.1.2 sub lineage initially identified in Rio de Janeiro [12] (**Figura 3**). Together, Tocantins genomes harbor the main mutations that characterize the P1 variant (including 11 mutations in the spike protein, such as E484K, N501Y, D614G) [11]. Furthermore, we detect a non-synonymous mutation in ORF3a (L106F) in 8 genomes (which appears to be increasing in frequency) and a non-synonymous mutation in ORF1b (M2260I) that is unprecedented. Importantly these both mutations appear in the same seven genomes, reinforcing our finding. Moreover, L106F appears in 6,687 GISAID genomes, sustaining the convergent character of this change.

Another feature of these sequences is a deletion of nine nucleotides in ORF1a (del 11288 to 11296) in all genomes excluding three amino acids (S-G-F) from these proteins. Below we show a phylogenetic reconstruction of 851 genomes retrieved from GISAID plus our 24 sequenced genomes from the Tocantins (**Figure 4**). The maximum likelihood tree shows the genomes with L106F mutation mentioned. Besides the alignment and phylogenetic analysis, we observed that the genomes described here formed a monophyletic branch with samples from Sao Paulo (the majority) and Goiás (Data not shown). This finding indicates that this potential P1-related lineage from the Tocantins emerged in São Paulo and spread to other Brazilian regions, including the states of Goiás and Tocantins.

**Figure 4.**
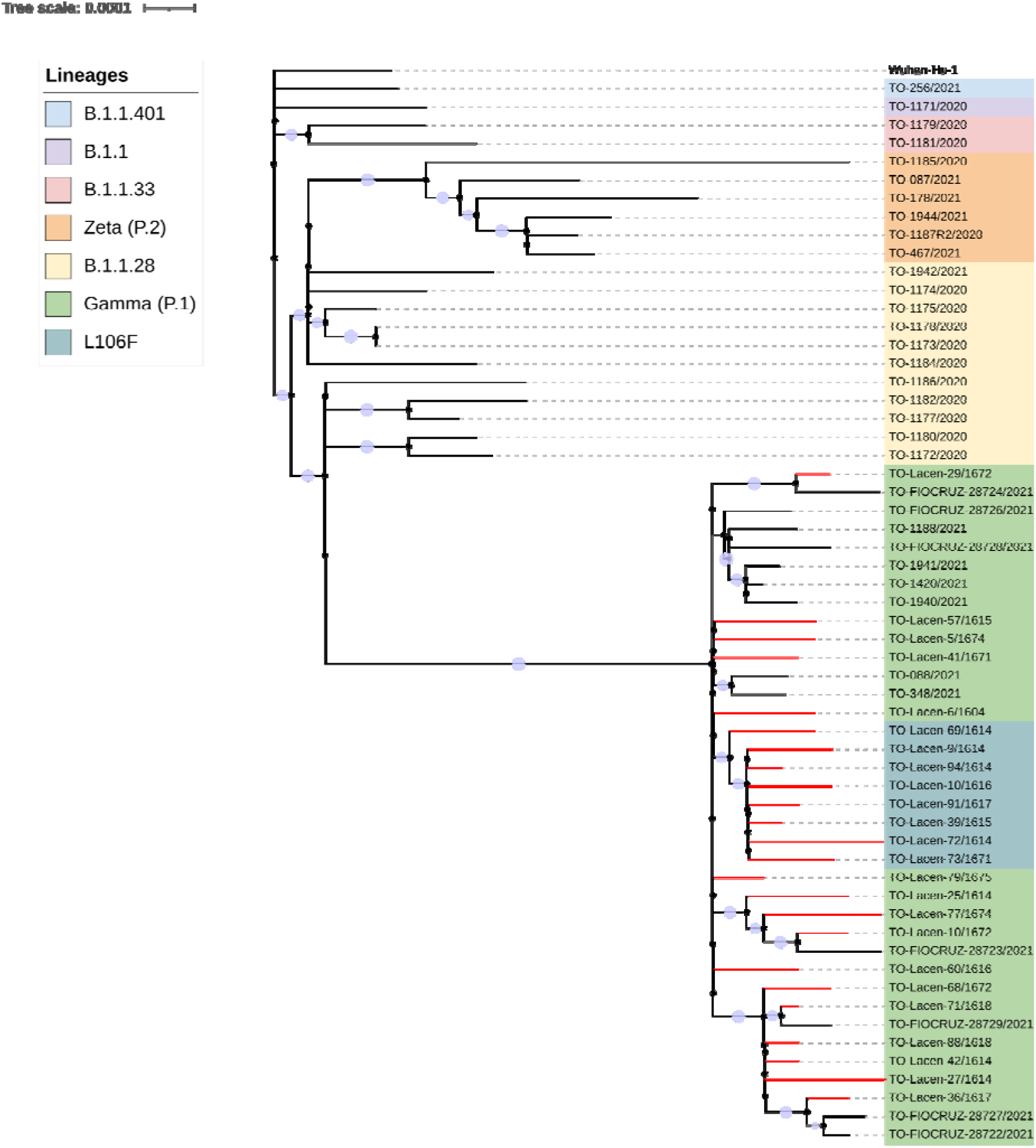
Phylogenetic tree of 59 SARS-CoV-2 genomes from Tocantins State. 34 genomes were retrieved from GISAID. Colored branches in red indicate the 24 genomes recently sequenced. Colored clade indicates lineage B. 1.1.401 in blue, B.1.1 in purple, B.1.1.33 in pink, Zeta (P.2) in orange, B.1.1.28 in yellow, Gamma (P.1) in green and the newly lineage Gamma (P.1) related in dark blue. The maximum likelihood tree was inferred with IQ-TREE2.

In addition, we identified 42 mutations of the SNPs type (Single Nucleotide Polymorphisms) and two mutations of the InDel type (Insertion-Deletion) in these genomes. Among the 42 SNPs identified, 41 are in coding regions of the viral genome and one in the 5’UTR portion (C241T). Eleven mutations in the coding region were characterized as synonymous and 30 of these were predicted with amino acid substitutions. Among these 30 non-synonymous substitutions, 43.3% are located in the S gene (with 13 mutations: L18F, T20N, P26S, D138Y, R190S, K417T, E484K, N501Y, D614G, H655Y, P681H, T1027I, V1176F). The ORF1ab, which comprises approximately 67% of the genome and encodes 16 non-structural proteins, had a total of 8 non-synonymous mutations (S1188L, K1230N, K1795Q, S1924I, T2846I, P4715L, E5665D, M6661I). Six substitutions are located in the N gene (P80R, T135I, S202C, S202T, R203K, G204R). One was found in ORF3a (L106F), one in ORF8 (E92K), and ORF 1b (M2260). Below we emphasize the evolutionary steps associated with the emergence of the P.1 and P.1.2 lineages adding our P.1-related (**Figure 5**).

**Figure 5.**
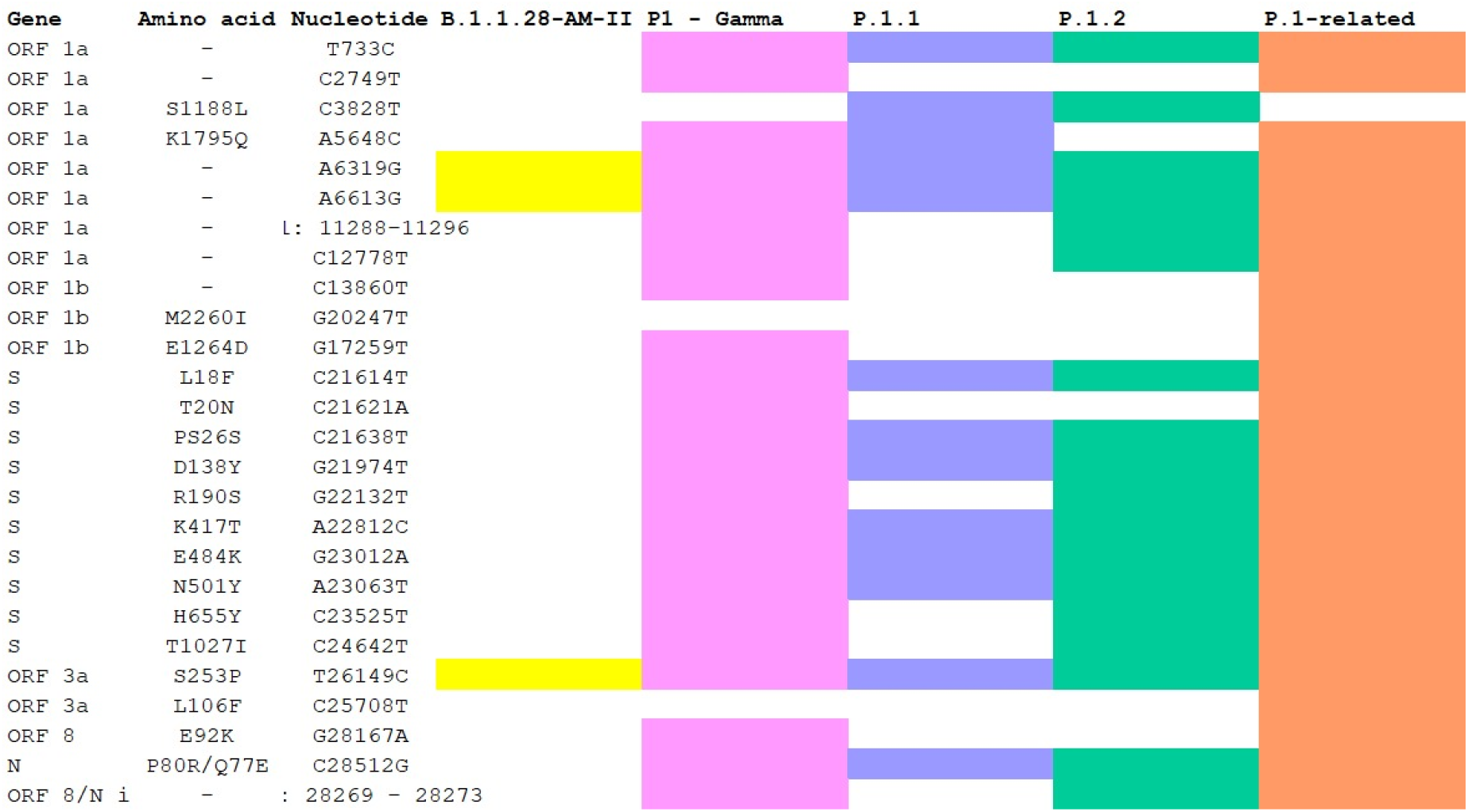
Mutations present in P.1 and P.1.2 lineages. Each line represents a mutation that emerged during the diversification of the B.1.1.28 lineage in Brazil originating the P.1, P.1.2 and now the P1-related identified here.

## Conclusion

In this report, we showed the predominance of the P.1 variant in the Tocantins. Moreover, we detect for the first time a potential new SARS-CoV-2 Gamma-related lineage through the sequencing of samples. We also show the spread and evolution of P.1 in Brazil. These findings reinforce the importance of continuous genomic surveillance in the state of Tocantins aiming to monitor and prevent the dispersion of variants. We reinforce the importance of genomic surveillance to support public health decisions and strategies that are advocated by the state health department, in order to prevent the spread of SARS-CoV-2 variants.

## Methods

### Sequencing

The 24 analyzed genomes were collected between May 26, 2021 and June 01, 2021 from 12 municipalities in the state of Tocantins/Brazil. Samples from patients with SARS-CoV-2 positive nasopharyngeal RT-PCR were collected at the Central Public Health Laboratory of the State of Tocantins. The study was approved by the Ethics Committee (33202820.7.1001.5348). Patients were aged between 01 and 80 years old, being 54.2% men and 45.8% women. Extraction of the genetic material was performed at the Central Public Health Laboratory of the State of Tocantins, with Extracta kit Viral RNA MVXA-P096 FAST (Loccus) in an automated extractor (Extracta 96, Loccus) following the manufacturer’s guidelines. Annealing of cDNA was conducted with 8.5 μl of viral RNA extracted from each sample. Libraries for whole virus genome sequencing were prepared according to version 3 of the ARTIC nCoV-2019 sequencing protocol (https://artic.network/ncov-2019). Long reads were generated with the LSK-109 sequencing kit, 24 native barcodes (NBD104 and NBD114 kits), and a MinION instrument (Oxford Nanopore, Oxford, UK). High accuracy base calling was carried out after sequencing from the fast5 files using the Oxford Nanopore Guppy tool. Assembly of the high accuracy base called fastq files was performed using the nCoV-2019 novel coronavirus bioinformatics protocol (https://artic.network/ncov-2019/ncov2019-bioinformatics-sop.html) with minimap2 [13] and medaka (https://github.com/nanoporetech/medaka) for consensus sequence generation.

Pango lineages were attributed to the newly assembled genomes using the Pangolin v3.1.5 software tool (https://pangolin.cog-uk.io/) [14]. In order to construct a phylogenetic tree we first aligned the 24 genomes recently sequenced and the 34 genomes deposited in GISAID with MAFFT v.7.480 [15]. The resulting alignment were subject to Maximum Likelihood phylogenetic analysis with IQ-TREE v.2.1.2 [16] under the Generalized Time Reversible GRT model of nucleotide substitution with empirical base frequencies (+F) and invariable sites (+I), as selected by the ModelFinder software. Bootstrap support was calculated with 10,000 tree replicates. The tree was visualized and edited using iTOL [17].

### Characterization of the new variant

First the mutational profile was investigated using the Nextclade tool (https://clades.nextstrain.org). After checking the occurrence of G20247T and C25708F, the sequences were obtained from the EpiCoV database in the GISAID (https://www.gisaid.org). In total, we downloaded 826 [51 Alpha (B.1.1.7), 12 Delta (B. 1.617.2), 84 Zeta (P.2), 313 (P.1.2), 366 (P.1)] SARS-CoV-2 brazilian genomes from GISAID and the reference genome from the NCBI database (NC_045512.2). We also included 24 genomes recently sequenced. The final dataset consisting of 851 was aligned using MAFFT v.7.480 [15]. Model testing was carried out using the ModelFinder software and the Generalized Time Reversible (GTR) model with empirical base frequencies (+F) plus FreeRate model (+R4). A Maximum Likelihood tree was generated using IQ-TREE v.2.1.2 [16] with 10,000 bootstraps as branch support. The tree was visualized and edited using iTOL [17].

All Bioinformatic and phylogenetic analyses were performed using the computational infrastructure of the Bioinformatics and Biotechnology Laboratory (LABINFTEC).

## Acknowledgments

We would like to thank all the authors and administrators of the GISAID database, which allowed this study of genomic epidemiology to be conducted properly. A full list acknowledging the authors publishing data used in this study can be found in the following file: Supplementary Table 2 (75.5 Kb). U.J.B.S., R.N.S. and A.B. are granted a post-doctoral scholarship (DTI-A) from CNPq. We acknowledge the support from the Rede Corona-ômica BR MCTI/FINEP affiliated to RedeVírus/MCTI (FINEP 01.20.0029.000462/20, CNPq 404096/2020-4) and Project CAPES # 88887.504570/2020-00 “EDITAL N° 9/2020 PREVENÇÃO E COMBATE A SURTOS, ENDEMIAS, EPIDEMIAS E PANDEMIAS”. We would like to thank Anderson Brito for his help with the Nextstrain tool. The authors would like to thank the whole team at Central Public Health Lab (LACEN) from Palmas, Tocantins, for their work and effort on COVID-19 diagnosis and surveillance. We would like to give a special thanks to Rafael Brustulin, who developed the COVID-19 sample management at LACEN, concerning the releasing of RT-PCR trails in a faster manner, directly wired to the Environment Management System Lab (GAL).

---

**Supplementary material – Table 1.**
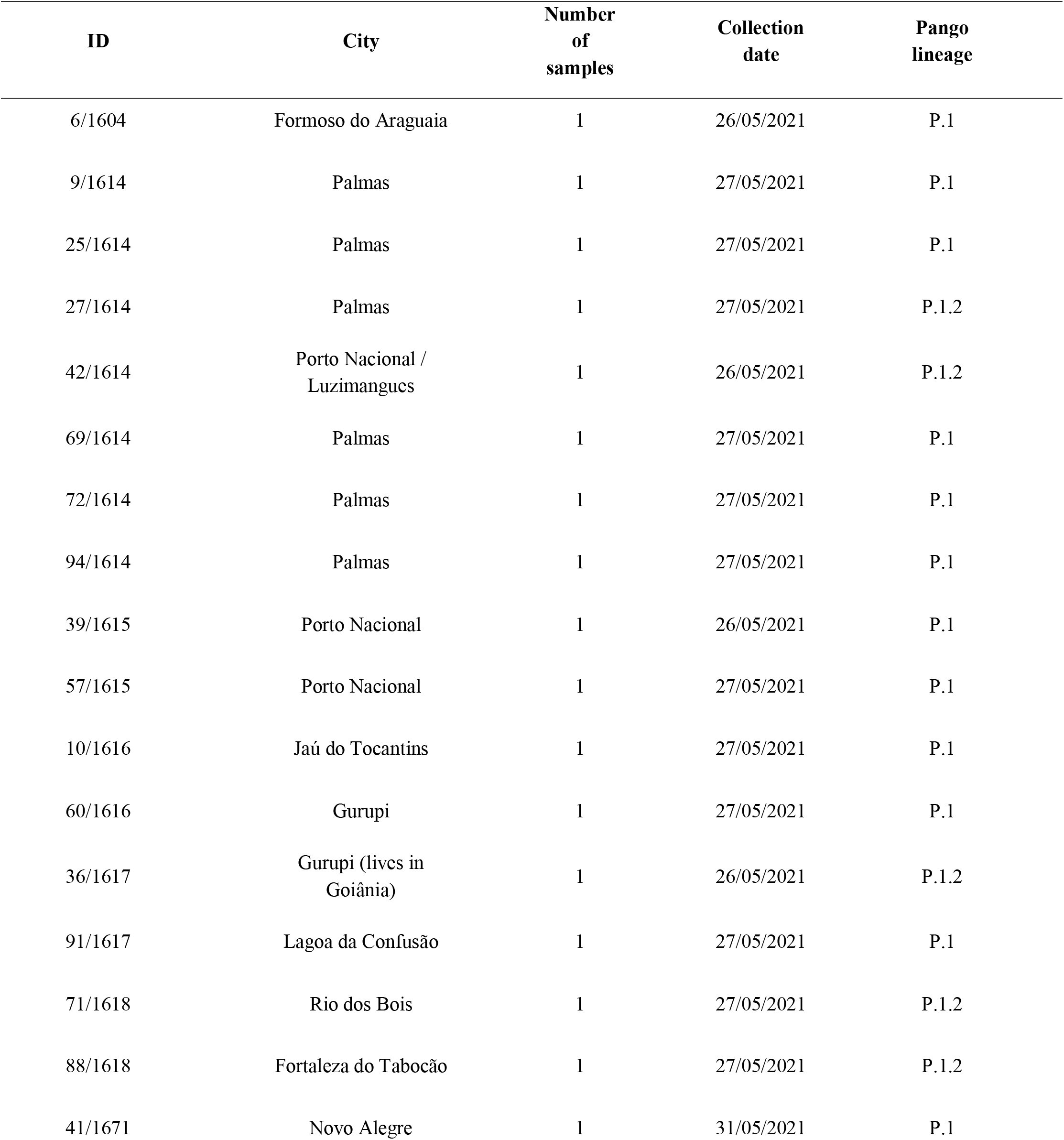
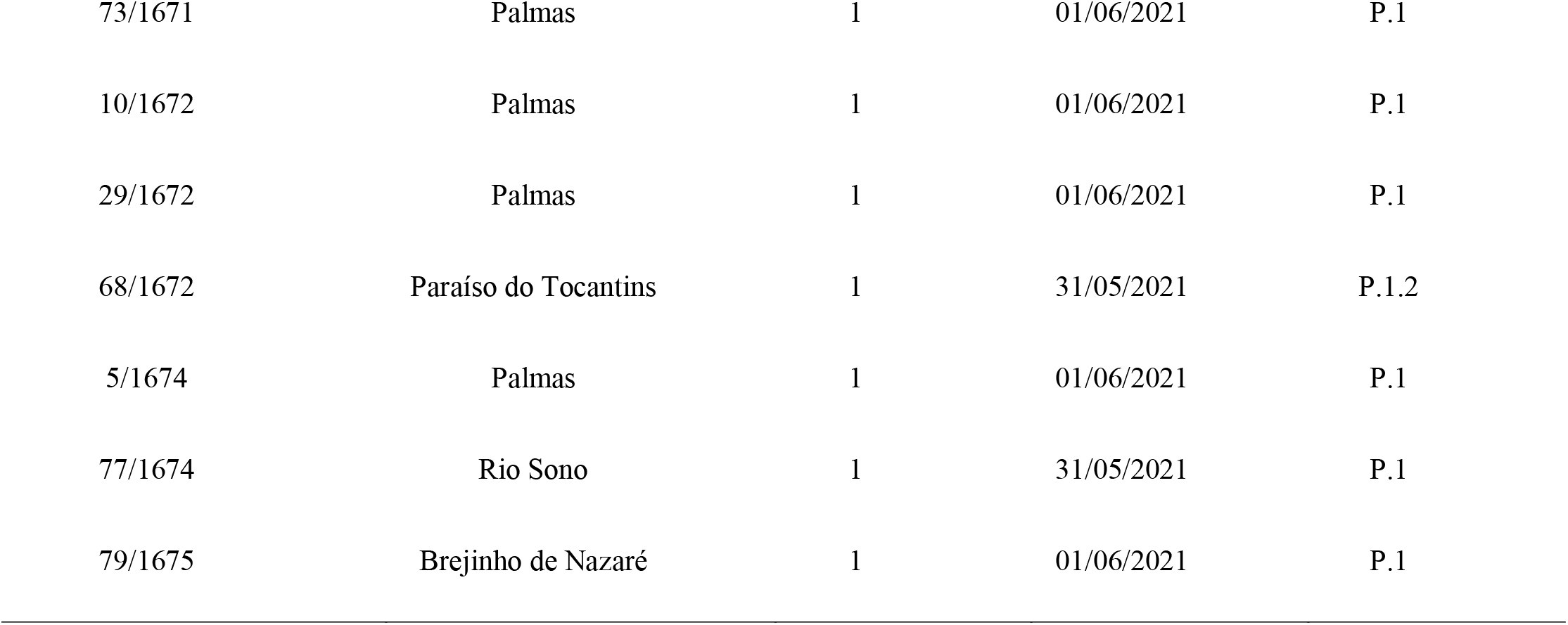
Location, date of collection and variant identified in the 24 samples from 12 cities in the state of Tocantins.

