## Supplementary Table 2 for "Detection of potential new SARS-CoV-2 Gamma-related lineage in Tocantins shows the spread and ongoing evolution of P.1 in Brazil"

We gratefully acknowledge the following Authors from the Originating laboratories responsible for obtaining the specimens, as well as the Submitting laboratories where the genome data were generated and shared via GISAID, on which this research is based.

All Submitters of data may be contacted directly via www.gisaid.org

Authors are sorted alphabetically.

| Accession ID | Originating Laboratory | Submitting Laboratory | Authors |
|---|---|---|---|
| EPI_ISL_2534881, EPI_ISL_2534943, EPI_ISL_2535018, EPI_ISL_2535101 | Laboratorio Central Noel Nutels | Bioinformatics Laboratory / LNCC | Luiz G P de Almeida, Alessandra P Lamarca, Ronaldo da Silva F Jr, Liliane Cavalcante, Alexandra L Gerber, Ana Paula de C Guimaraes, Douglas Terra Machado, Cassia Alves, Diana Mariani, Cintia Policarpo, Gleidson da Silva de Oliveira, Mario Sergio Ribeiro, Silvia Carvalho, Flavio Dias da Silva, Marcio Henrique de Oliveira Garcia, Leandro Magalhaes de Souza, Cristiane Gomes da Silva, Caio Luiz Pereira Ribeiro, Andrea Cony Cavalcanti, Claudia Maria Braga de Mello, Amilcar Tanuri, Ana Tereza R Vasconcelos |
| EPI_ISL_2535175, EPI_ISL_2535189, EPI_ISL_2535220 | Unidade de apoio ao diagnostico da COVID - UNADIG | Bioinformatics Laboratory / LNCC | Luiz G P de Almeida, Alessandra P Lamarca, Ronaldo da Silva F Jr, Liliane Cavalcante, Alexandra L Gerber, Ana Paula de C Guimaraes, Douglas Terra Machado, Cassia Alves, Diana Mariani, Cintia Policarpo, Gleidson da Silva de Oliveira, Mario Sergio Ribeiro, Silvia Carvalho, Flavio Dias da Silva, Marcio Henrique de Oliveira Garcia, Leandro Magalhaes de Souza, Cristiane Gomes da Silva, Caio Luiz Pereira Ribeiro, Andrea Cony Cavalcanti, Claudia Maria Braga de Mello, Amilcar Tanuri, Ana Tereza R Vasconcelos |
| EPI_ISL_2536269, EPI_ISL_2536270 | Laboratory of Respiratory Viruses and Measles, Oswaldo Cruz Institute, FIOCRUZ | Laboratory of Respiratory Viruses and Measles, Oswaldo Cruz Institute, FIOCRUZ | Paola Resende, Patricia Brasil, Luciana Appolinario, Fernando Motta, Anna Carolina Paixao, Ana Carolina Mendonca, Alice Sampaio Rocha, Taina Venas, Elisa Cavalcante Pereira, Renata Serrano Lopes, Marilda Siqueira on behalf of the Fiocruz COVID-19 Genomic Surveillance Network |
| EPI_ISL_2536327 | Laboratorio Central de Saude Publica do Estado da Paraiba (LACEN-PB) | Laboratory of Respiratory Viruses and Measles, Oswaldo Cruz Institute, FIOCRUZ | Paola Resende, Luciana Appolinario, Fernando Motta, Anna Carolina Paixao, Ana Carolina Mendonca, Alice Sampaio Rocha, Taina Venas, Elisa Cavalcante Pereira, Renata Serrano Lopes, Joao Felipe Bezerra, Dalane Loudal Florentino Teixeira, Marilda Siqueira on behalf of the Fiocruz COVID-19 Genomic Surveillance Network |
| EPI_ISL_2551529 | Belo Horizonte center-south emergency care unit - UPA-BH | Laboratório de Virologia Clínica e Molecular | Erick Gustavo Dorlass, Karine Lima Lourenço, Rubens Daniel Miserani Magalhães, Hugo Sato, Alex Fiorini, Renata Peixoto, Helena Perez Coelho, Ana Paula Salles Fernandes, Bruna Larotonda Telezynski, Guilherme Pereira Scagion, Tatiana Ometto, Luciano Matsumiya Thomazelli, Danielle Bruna Leal Oliveira, Edison Luiz Durigon, Flavio Fonseca e Santuza Teixeira |
| EPI_ISL_2551535 | Hospital Infantnil Darcy Vargas | Laboratório de Virologia Clínica e Molecular | Erick Gustavo Dorlass, Karine Lima Lourenço, Rubens Daniel Miserani Magalhães, Hugo Sato, Alex Fiorini, Renata Peixoto, Helena Perez Coelho, Ana Paula Salles Fernandes, Bruna Larotonda Telezynski, Guilherme Pereira Scagion, Tatiana Ometto, Luciano Matsumiya Thomazelli, Danielle Bruna Leal Oliveira, Edison Luiz Durigon, Flavio Fonseca e Santuza Teixeira |
| EPI_ISL_2551536 | Laboratório de Virologia Clínica e Molecular | Laboratório de Virologia Clínica e Molecular | Erick Gustavo Dorlass, Karine Lima Lourenço, Rubens Daniel Miserani Magalhães, Hugo Sato, Alex Fiorini, Renata Peixoto, Helena Perez Coelho, Ana Paula Salles Fernandes, Bruna Larotonda Telezynski, Guilherme Pereira Scagion, Tatiana Ometto, Luciano Matsumiya Thomazelli, Danielle Bruna Leal Oliveira, Edison Luiz Durigon, Flavio Fonseca e Santuza Teixeira |
| EPI_ISL_2614183, EPI_ISL_2614238, EPI_ISL_2614285, EPI_ISL_2614287, EPI_ISL_2614310 | Laboratory of Respiratory Viruses and Measles, Oswaldo Cruz Institute, FIOCRUZ | Laboratory of Respiratory Viruses and Measles, Oswaldo Cruz Institute, FIOCRUZ | Paola Resende, Luciana Appolinario, Fernando Motta, Anna Carolina Paixao, Ana Carolina Mendonca, Alice Sampaio Rocha, Taina Venas, Elisa Cavalcante Pereira, Renata Serrano Lopes, Marilda Siqueira on behalf of the Fiocruz COVID-19 Genomic Surveillance Network |
| EPI_ISL_2614357 | Labortorio Central de Saude Publica do Estado do Rio de Janeiro (LACEN/RJ) | Laboratory of Respiratory Viruses and Measles, Oswaldo Cruz Institute, FIOCRUZ | Paola Resende, Luciana Appolinario, Fernando Motta, Anna Carolina Paixao, Ana Carolina Mendonca, Alice Sampaio Rocha, Taina Venas, Elisa Cavalcante Pereira, Renata Serrano Lopes, Andrea Cony Cavalcanti, Marilda Siqueira on behalf of the Fiocruz COVID-19 Genomic Surveillance Network |
| EPI_ISL_2614527, EPI_ISL_2614528 | Instituto Adolfo Lutz - Regional de Campinas | Instituto Adolfo Lutz, Interdiciplinary Procedures Center, Strategic Laboratory | Claudio Tavares Sacchi, Claudia Regina Gonçalves, Erica Valessa Ramos Gomes, Karoline Rodrigues Campos, Caio Vinicius Dias Lopes, Leonardo Jose Tadeu de Araujo |
| EPI_ISL_2614539 | Centro de Diagnostico SECEDI | Instituto Adolfo Lutz, Interdiciplinary Procedures Center, Strategic Laboratory | Claudio Tavares Sacchi, Claudia Regina Gonçalves, Erica Valessa Ramos Gomes, Karoline Rodrigues Campos, Caio Vinicius Dias Lopes, Leonardo Jose Tadeu de Araujo |
| EPI_ISL_2614554, EPI_ISL_2614556, EPI_ISL_2614564 | IAL Presidente Prudente | Instituto Adolfo Lutz, Interdiciplinary Procedures Center, Strategic Laboratory | Claudio Tavares Sacchi, Claudia Regina Gonçalves, Erica Valessa Ramos Gomes, Karoline Rodrigues Campos, Caio Vinicius Dias Lopes, Leonardo Jose Tadeu de Araujo |
| EPI_ISL_2614591 | Hospital Municipal Enf Antonio Policarpo de Oliveira | Instituto Adolfo Lutz, Interdiciplinary Procedures Center, Strategic Laboratory | Claudio Tavares Sacchi, Claudia Regina Gonçalves, Erica Valessa Ramos Gomes, Karoline Rodrigues Campos, Caio Vinicius Dias Lopes, Leonardo Jose Tadeu de Araujo |
| EPI_ISL_2615400 | IAL Presidente Prudente | Instituto Adolfo Lutz, Interdiciplinary Procedures Center, Strategic Laboratory | Claudio Tavares Sacchi, Claudia Regina Gonçalves, Erica Valessa Ramos Gomes, Karoline Rodrigues Campos, Caio Vinicius Dias Lopes, Leonardo Jose Tadeu de Araujo |
| EPI_ISL_2617591 | HLAGYN - Laboratorio de Imunologia de Transplantes de Goias | HLAGYN - Laboratorio de Imunologia de Transplantes de Goias | Fernando Antonio Vinhal dos Santos, Erika Lopes Rocha Batista, Alessandro Leonardo Alvares Magalhaes, Frederico Rodrigues Vinhal, Sabrina Sara Moreira Duarte, Lucas Carlos Gomes Pereira, Daniel Ferreira de Sousa |
| EPI_ISL_2627737 | Belo Horizonte center-south emergency care unit - UPA-BH | Laboratório de Virologia Clínica e Molecular | Erick Gustavo Dorlass, Karine Lima Lourenço, Rubens Daniel Miserani Magalhães, Hugo Sato, Alex Fiorini, Renata Peixoto, Helena Perez Coelho, Ana Paula Salles Fernandes, Bruna Larotonda Telezynski, Guilherme Pereira Scagion, Tatiana Ometto, Luciano Matsumiya Thomazelli, Danielle Bruna Leal Oliveira, Edison Luiz Durigon, Flavio Fonseca e Santuza Teixeira |
| EPI_ISL_2629751 | Laboratório de Virologia Molecular - Universidade Federal do Rio de Janeiro | Laboratório de Virologia Molecular - Universidade Federal do Rio de Janeiro | Filipe Romero Rebello Moreira, Mirela D'arc, Diana Mariani, Alice Laschuk Herlinger, Francine Bittencourt Schiffler, Átila Duque Rossi, Isabela de Carvalho Leitão, Thamiris dos Santos Miranda, Matheus Augusto Calvano Cosentino, Marcelo Calado de Paula Tôrres, Raíssa Mirella dos Santos Cunha da Costa, Cássia Cristina Alves Gonçalves, Débora Souza Faffe, Rafael Mello Galliez, Orlando da Costa Ferreira Junior, Renato Santana de Aguiar,, André Felipe Andrade dos Santos, Carolina Moreira Voloch, Terezinha Marta Pereira Pinto Castineiras, Amilcar Tanuri |
| EPI_ISL_2645519, EPI_ISL_2645522, EPI_ISL_2645523, EPI_ISL_2645524, EPI_ISL_2645525, EPI_ISL_2645526, EPI_ISL_2645527, EPI_ISL_2645528 | Laboratorio Central de Saude Publica do Estado do Espirito Santo (LACEN/ES) | Laboratory of Respiratory Viruses and Measles, Oswaldo Cruz Institute, FIOCRUZ | Paola Resende, Luciana Appolinario, Fernando Motta, Anna Carolina Paixao, Ana Carolina Mendonca, Alice Sampaio Rocha, Taina Venas, Eilisa Cavalcante Pereira, Renata Serrano Lopes, Rodrigo Ribeiro Rodrigues, Marilda Siqueira on behalf of the Fiocruz COVID-19 Genomic Surveillance Network |
| EPI_ISL_2660484, EPI_ISL_2660525 | Laboratorio Central de Saude Publica do Estado de Minas Gerais (LACEN/MG) | Laboratory of Respiratory Viruses and Measles, Oswaldo Cruz Institute, FIOCRUZ | Paola Resende, Luciana Appolinario, Fernando Motta, Anna Carolina Paixao, Ana Carolina Mendonca, Alice Sampaio Rocha, Taina Venas, Elisa Cavalcante Pereira, Renata Serrano Lopes, Andre Felipe Leal Bernardes, Marilda Siqueira on behalf of the Fiocruz COVID-19 Genomic Surveillance Network |
| EPI_ISL_2661750 | Laboratorio Central de Saude Publica do Estado do Rio Grande do Sul (LACEN-RS) | Laboratory of Respiratory Viruses and Measles, Oswaldo Cruz Institute, FIOCRUZ | Paola Resende, Luciana Appolinario, Fernando Motta, Anna Carolina Paixao, Ana Carolina Mendonca, Alice Sampaio Rocha, Taina Venas, Elisa Cavalcante Pereira, Renata Serrano Lopes, Tatiana Schaffer Gregianini, Richard Salvato, Marilda Siqueira on behalf of the Fiocruz COVID-19 Genomic Surveillance Network |
| EPI_ISL_2691107, EPI_ISL_2691108, EPI_ISL_2691112, EPI_ISL_2691114, EPI_ISL_2691117, EPI_ISL_2691120 | Instituto Adolfo Lutz - Regional de Bauru | Instituto Adolfo Lutz, Interdiciplinary Procedures Center, Strategic Laboratory | Claudio Tavares Sacchi, Claudia Regina Gonçalves, Erica Valessa Ramos Gomes, Karoline Rodrigues Campos, Caio Vinicius Dias Lopes, Leonardo Jose Tadeu de Araujo |
| EPI_ISL_2691179 | Instituto Adolfo Lutz - Reginal de Sao José do Rio Preto | Instituto Adolfo Lutz, Interdiciplinary Procedures Center, Strategic Laboratory | Claudio Tavares Sacchi, Claudia Regina Gonçalves, Erica Valessa Ramos Gomes, Karoline Rodrigues Campos, Caio Vinicius Dias Lopes, Leonardo Jose Tadeu de Araujo |
| EPI_ISL_2691329, EPI_ISL_2691379, EPI_ISL_2691559 | Laboratorio Central Noel Nutels | Bioinformatics Laboratory / LNCC | Luiz G P de Almeida, Alessandra P Lamarca, Ronaldo da Silva F Jr, Liliane Cavalcante, Alexandra L Gerber, Ana Paula de C Guimaraes, Douglas Terra Machado, Cassia Alves, Diana Mariani, Cintia Policarpo, Gleidson da Silva de Oliveira, Mario Sergio Ribeiro, Silvia Carvalho, Flavio Dias da Silva, Marcio Henrique de Oliveira Garcia, Leandro Magalhaes de Souza, Cristiane Gomes da Silva, Caio Luiz Pereira Ribeiro, Andrea Cony Cavalcanti, Claudia Maria |

We gratefully acknowledge the following Authors from the Originating laboratories responsible for obtaining the specimens, as well as the Submitting laboratories where the genome data were generated and shared via GISAID, on which this research is based.

All Submitters of data may be contacted directly via www.gisaid.org

Authors are sorted alphabetically.

| Accession ID | Originating Laboratory | Submitting Laboratory | Authors |
|---|---|---|---|
| EPI_ISL_2274979 | Aeroporto Internacional de Guarulhos | Instituto Adolfo Lutz, Interdiciplinary Procedures Center, Strategic Laboratory | Claudio Tavares Sacchi, Claudia Regina Gonçalves, Erica Valessa Ramos Gomes, Karoline Rodrigues Campos, Caio Vinicius Dias Lopes, Leonardo Jose Tadeu de Araujo |
| EPI_ISL_2311861 | Fundação Ezequiel Dias | Coordenação Geral de Laboratórios de Saúde Pública (CGLAB/DAEVS/SVS/MS) | Vagner Fonseca, et al. |
| EPI_ISL_2443545 | Lboratorio Central de Saude Publica do Estado do Parana (LACEN/PR) | Laboratory of Respiratory Viruses and Measles, Oswaldo Cruz Institute, FIOCRUZ | Paola Resende, Luciana Appolinario, Fernando Motta, Anna Carolina Paixao, Ana Carolina Mendonca, Alice Sampaio Rocha, Taina Venas, Elisa Cavalcante Pereira, Renata Serrano Lopes, Irina Riediger, Marilda Siqueira on behalf of the Fiocruz COVID-19 Genomic Surveillance Network |
| EPI_ISL_2466268 | Laboratorio Central de Saude Publica do Estado do Rio de Janeiro (LACEN-RJ) | Laboratory of Respiratory Viruses and Measles, Oswaldo Cruz Institute, FIOCRUZ | Paola Resende, Luciana Appolinario, Fernando Motta, Anna Carolina Paixao, Ana Carolina Mendonca, Alice Sampaio Rocha, Taina Venas, Elisa Cavalcante Pereira, Renata Serrano Lopes, Andrea Cony Cavalcanti, Marilda Siqueira on behalf of the Fiocruz COVID-19 Genomic Surveillance Network |
| EPI_ISL_2645412, EPI_ISL_2645413, EPI_ISL_2645414, EPI_ISL_2645415 | Laboratorio Central de Saude Publica do Estado Maranhao (LACEN-MA) | Laboratory of Respiratory Viruses and Measles, Oswaldo Cruz Institute, FIOCRUZ | Paola Resende, Luciana Appolinario, Fernando Motta, Anna Carolina Paixao, Ana Carolina Mendonca, Alice Sampaio Rocha, Taina Venas, Elisa Cavalcante Pereira, Renata Serrano Lopes, Lidio Gonçalves Lima Neto, Marilda Siqueira on behalf of the Fiocruz COVID-19 Genomic Surveillance Network |
| EPI_ISL_2645416 | Laboratorio Central de Saude Publica do Estado Maranhao (LACEN-MA) | Laboratory of Respiratory Viruses and Measles, Oswaldo Cruz Institute, FIOCRUZ | Paola Resende, Alex Pauvolid-Corrêa, Mia Ferreira de Araujo, Ana Beatriz Machado Lima, Luciana Appolinario, Fernando Motta, Anna Carolina Paixao, Ana Carolina Mendonca, Alice Sampaio Rocha, Taina Venas, Elisa Cavalcante Pereira, Renata Serrano Lopes, Lidio Gonçalves Lima Neto, Marilda Siqueira on behalf of the Fiocruz COVID-19 Genomic Surveillance Network |
| EPI_ISL_2645417 | Laboratorio Central de Saude Publica do Estado do Maranhao (LACEN-MA) | Laboratory of Respiratory Viruses and Measles, Oswaldo Cruz Institute, FIOCRUZ | Paola Resende, Luciana Appolinario, Fernando Motta, Anna Carolina Paixao, Ana Carolina Mendonca, Alice Sampaio Rocha, Taina Venas, Elisa Cavalcante Pereira, Renata Serrano Lopes, Lidio Gonçalves Lima Neto, Marilda Siqueira on behalf of the Fiocruz COVID-19 Genomic Surveillance Network |
| EPI_ISL_2645418 | Laboratorio Central de Saude Publica do Estado do Maranhao (LACEN-MA) | Laboratory of Respiratory Viruses and Measles, Oswaldo Cruz Institute, FIOCRUZ | Paola Resende, Alex Pauvolid-Corrêa, Mia Ferreira de Araujo, Ana Beatriz Machado Lima, Luciana Appolinario, Fernando Motta, Anna Carolina Paixao, Ana Carolina Mendonca, Alice Sampaio Rocha, Taina Venas, Elisa Cavalcante Pereira, Renata Serrano Lopes, Lidio Gonçalves Lima Neto, Marilda Siqueira on behalf of the Fiocruz COVID-19 Genomic Surveillance Network |
| EPI_ISL_2677318 | Labortorio Central de Saude Publica do Estado do Rio de Janeiro (LACEN/RJ) | Laboratory of Respiratory Viruses and Measles, Oswaldo Cruz Institute, FIOCRUZ | Paola Resende, Alex Pauvolid-Corrêa, Mia Ferreira de Araujo, Ana Beatriz Machado Lima,Luciana Appolinario, Fernando Motta, Anna Carolina Paixao, Ana Carolina Mendonca, Alice Sampaio Rocha, Taina Venas, Elisa Cavalcante Pereira, Renata Serrano Lopes, Andrea Cony Cavalcanti, Marilda Siqueira on behalf of the Fiocruz COVID-19 Genomic Surveillance Network |

We gratefully acknowledge the following Authors from the Originating laboratories responsible for obtaining the specimens, as well as the Submitting laboratories where the genome data were generated and shared via GISAID, on which this research is based.

All Submitters of data may be contacted directly via www.gisaid.org

Authors are sorted alphabetically.

| Accession ID | Originating Laboratory | Submitting Laboratory | Authors |
|---|---|---|---|
| EPI_ISL_1213270 | Laboratório HLA/UERJ | Bioinformatics Laboratory / LNCC | Alessandra P Lamarca, Luiz G P de Almeida, Ronaldo da Silva Francisco Jr, Lucymara Fassarella Agnez Lima, Kátia Castanho Scortecci, Vinícius Pietta Perez, Otavio J. Brustolini, Eduardo Sérgio Soares Sousa, Danielle Angst Secco, Angela Maria Guimarães Santos, George Rego Albuquerque, Ana Paula Melo Mariano, Bianca Mendes Maciel, Alexandra L Gerber, Ana Paula de C Guimarães, Paulo Ricardo Nascimento, Francisco Paulo Freire Neto, Sandra Rocha Gadelha, Luís Cristóvão Porto, Eloiza Helena Campana, Selma Maria Bezerra Jeronimo, Ana Tereza R Vasconcelos |
| EPI_ISL_1213392 | LBM/UFPB | Bioinformatics Laboratory / LNCC | Alessandra P Lamarca, Luiz G P de Almeida, Ronaldo da Silva Francisco Jr, Lucymara Fassarella Agnez Lima, Kátia Castanho Scortecci, Vinícius Pietta Perez, Otavio J. Brustolini, Eduardo Sérgio Soares Sousa, Danielle Angst Secco, Angela Maria Guimarães Santos, George Rego Albuquerque, Ana Paula Melo Mariano, Bianca Mendes Maciel, Alexandra L Gerber, Ana Paula de C Guimarães, Paulo Ricardo Nascimento, Francisco Paulo Freire Neto, Sandra Rocha Gadelha, Luís Cristóvão Porto, Eloiza Helena Campana, Selma Maria Bezerra Jeronimo, Ana Tereza R Vasconcelos |
| EPI_ISL_1381052, EPI_ISL_1381056 | IAL Regional de Santo Andre | Instituto Adolfo Lutz, Interdiciplinary Procedures Center, Strategic Laboratory | Claudio Tavares Sacchi, Claudia Regina Gonçalves, Erica Valessa Ramos Gomes, Karoline Rodrigues Campos, Caio Vinicius Dias Lopes |
| EPI_ISL_1445090 | POLICLINICA COVID 19 ITAPETININGA | Instituto Butantan / Mendelics | Dimas Tadeu Covas, Sandra Coccuzzo Sampaio, Maria Carolina Elias, José Salvatore Leister Patané, Vincent Louis Viala, Antonio Jorge Martins, Ricardo Haddad, Claudia Renata dos Santos Barros, Elaine Cristina Marqueze, Raul Machado Neto, Debora Botequio Moretti, Bibiana Santos, João Paulo Kitajima, Erika Freitas, David Schlesinger, Simone Kashima, Evandra Strazza Rodrigues, Svetoslav Nanev Slavov, Elaine Vieira dos Santos, Rafael dos Santos Bezerra, Luiz Carlos Junior de Alcantara, Marta Giovanetti, Vagner Fonseca, Flavia Aburjaile, Rodrigo Tocantins Calado. |
| EPI_ISL_1445126 | UNIDADE DE PRONTO ATENDIMENTO UPA DRA ANA OLIVIA BENTIVOGLIO | Instituto Butantan / Mendelics | Dimas Tadeu Covas, Sandra Coccuzzo Sampaio, Maria Carolina Elias, José Salvatore Leister Patané, Vincent Louis Viala, Antonio Jorge Martins, Ricardo Haddad, Claudia Renata dos Santos Barros, Elaine Cristina Marqueze, Raul Machado Neto, Debora Botequio Moretti, Bibiana Santos, João Paulo Kitajima, Erika Freitas, David Schlesinger, Simone Kashima, Evandra Strazza Rodrigues, Svetoslav Nanev Slavov, Elaine Vieira dos Santos, Rafael dos Santos Bezerra, Luiz Carlos Junior de Alcantara, Marta Giovanetti, Vagner Fonseca, Flavia Aburjaile, Rodrigo Tocantins Calado. |
| EPI_ISL_1445146 | CS II EGIDIO BRUNHARA MORRO AGUDO | Instituto Butantan / Mendelics | Dimas Tadeu Covas, Sandra Coccuzzo Sampaio, Maria Carolina Elias, José Salvatore Leister Patané, Vincent Louis Viala, Antonio Jorge Martins, Ricardo Haddad, Claudia Renata dos Santos Barros, Elaine Cristina Marqueze, Raul Machado Neto, Debora Botequio Moretti, Bibiana Santos, João Paulo Kitajima, Erika Freitas, David Schlesinger, Simone Kashima, Evandra Strazza Rodrigues, Svetoslav Nanev Slavov, Elaine Vieira dos Santos, Rafael dos Santos Bezerra, Luiz Carlos Junior de Alcantara, Marta Giovanetti, Vagner Fonseca, Flavia Aburjaile, Rodrigo Tocantins Calado. |
| EPI_ISL_1445150 | CENTRO DE REFERENCIA DO IDOSO DR HUMBERTO MENDES DE CARVALHO | Instituto Butantan / Mendelics | Dimas Tadeu Covas, Sandra Coccuzzo Sampaio, Maria Carolina Elias, José Salvatore Leister Patané, Vincent Louis Viala, Antonio Jorge Martins, Ricardo Haddad, Claudia Renata dos Santos Barros, Elaine Cristina Marqueze, Raul Machado Neto, Debora Botequio Moretti, Bibiana Santos, João Paulo Kitajima, Erika Freitas, David Schlesinger, Simone Kashima, Evandra Strazza Rodrigues, Svetoslav Nanev Slavov, Elaine Vieira dos Santos, Rafael dos Santos Bezerra, Luiz Carlos Junior de Alcantara, Marta Giovanetti, Vagner Fonseca, Flavia Aburjaile, Rodrigo Tocantins Calado. |
| EPI_ISL_1468468 | Secretaria municipal de saude de Itapolis | Instituto Adolfo Lutz, Interdiciplinary Procedures Center, Strategic Laboratory | Claudio Tavares Sacchi, Claudia Regina Gonçalves, Erica Valessa Ramos Gomes, Karoline Rodrigues Campos, Caio Vinicius Dias Lopes |
| EPI_ISL_1469623 | Diretoria de Vigilância em Saúde | Epiclin | Fernando Hayashi Sant'Anna, Ana Paula Muterle, Janira Prichula, Juliana Comerlato, Carolina Comerlato, Eliana Márcia Da Ros Wendland |
| EPI_ISL_1493585 | Unidade de Saude Dr Phebo de Oliveira Roge Ferreira | Instituto Adolfo Lutz, Interdiciplinary Procedures Center, Strategic Laboratory | Claudio Tavares Sacchi, Claudia Regina Gonçalves, Erica Valessa Ramos Gomes, Karoline Rodrigues Campos, Caio Vinicius Dias Lopes |
| EPI_ISL_1493589 | CS II Dr Jose Ferreira Telles | Instituto Adolfo Lutz, Interdiciplinary Procedures Center, Strategic Laboratory | Claudio Tavares Sacchi, Claudia Regina Gonçalves, Erica Valessa Ramos Gomes, Karoline Rodrigues Campos, Caio Vinicius Dias Lopes |
| EPI_ISL_1580269 | Laboratório de Biologia Molecular do Hospital das Clínicas da Faculdade de Medicina de Botucatu/SP | Laboratórios de Genômica Funcional (FCA/UNESP) e Biologia Molecular (FMB-HC/UNESP) - Rede de Vigilância Genômica (Vigenômica)/UNESP | Patrícia Akemi Assato; Felipe Allan da Silva da Costa; Bianca Cechetto Carlos; Flavia Hebner Barbosa Trovão; Guilherme Targino Valente; Rejane Maria Tommasini Grotto; Jayme A. Souza-Neto. |
| EPI_ISL_1716484 | LACEN (Laboratorio de Saude Publica Dr. Giovanni Cysneiros) | LGBio (Laboratorio de Genetica & Biodiversidade) | Mariana Pires de Campos Telles, Daniela de Melo e Silva, Elisangela de Paula Silveira Lacerda, Renata de Oliveira Dias, Rhewter Nunes, Cintia Pelegrineti Targueta de Azevedo Brito, Ramilla dos Santos Braga, Thais Guimarães Castro, Thays Millena Alves Pedroso, Amanda Alves de Melo, Aparecido Divino da Cruz, Luiz Augusto Pereira, Thais Cidália Vieira Gigonzac, Marc Alexandre Duarte Gigonzac, Alex Honda Bernardes, Francyelli Mello Andrade |
| EPI_ISL_1731589 | Instituto Adolfo Lutz Central | Instituto Adolfo Lutz, Interdiciplinary Procedures Center, Strategic Laboratory | Claudio Tavares Sacchi, Claudia Regina Gonçalves, Erica Valessa Ramos Gomes, Karoline Rodrigues Campos, Caio Vinicius Dias Lopes, Leonardo Jose Tadeu de Araujo, Katia Correa de Oliveira Santos |
| EPI_ISL_1734881 | Instituto de Biotecnologia - UNESP-Botucatu-SP | Instituto de Biotecnologia - UNESP-Botucatu-SP | Fábio Sossai Possebon; Leila Sabrina Ullmann; Cecília Artico Banho; Cíntia Bittar; Guilherme Campos; Helena Lage Ferreira; Jorge A. Petroli Marchesi; Lívia Sacchetto; Maisa C. Pereira Parra; Marília Moraes; Maurício L. Nogueira; Paula Rahal; Paulo Inacio da Costa; João Pessoa Araújo Jr. |
| EPI_ISL_1752645 | Instituto Adolfo Lutz - Regional de Ribeirao Preto | Instituto Adolfo Lutz, Interdiciplinary Procedures Center, Strategic Laboratory | Claudio Tavares Sacchi, Claudia Regina Gonçalves, Erica Valessa Ramos Gomes, Karoline Rodrigues Campos, Caio Vinicius Dias Lopes, Leonardo Jose Tadeu de Araujo, Katia Correa de Oliveira Santos |
| EPI_ISL_1795193 | USF SALERNO | Instituto Butantan / ESALQ-Piracicaba | Instituto Butantan: Alexander Roberto Precioso, Dimas Tadeu Covas, Sandra Coccuzzo Sampaio, Maria Carolina Elias, José Salvatore Leister Patané, Vincent Louis Viala, Antonio Jorge Martins, Ricardo Haddad, Claudia Renata dos Santos Barros, Elaine Cristina Marqueze, Raul Machado Neto, Debora Botequio Moretti. Centro de Genômica Funcional da ESALQ: Luiz Lehmann Coutinho, Ricardo Augusto Brassaloti, Raquel de Lello Rocha Campos Cassano. NGS Soluções Genômicas: Pilar Drummond Sampaio Corrêa Mariani. FZEA-USP Pirassununga: Mirele Daiana Poleti, Jessika Cristina Chagas Lesbon, Elisangela Chicaroni Mattos, Heidge Fukumasu. USP-Botucatu: Rejane Maria Tommasini Grotto, Jayme A. Souza-Neto, Guilherme Targino Valente, Patricia Akemi Assato, Felipe Allan da Silva da Costa, Bianca Cechetto Carlos. Mendelics: Bibiana Santos, João Paulo Kitajima, Erika Freitas, David Schlesinger. Hemocentro Ribeirão Preto: Simone Kashima, Evandra Strazza Rodrigues, Svetoslav Nanev Slavov, Elaine Vieira dos Santos, Rafael dos Santos Bezerra, Luiz Carlos Junior de Alcantara, Marta Giovanetti, Vagner Fonseca, Flavia Aburjaile, Rodrigo Tocantins Calado. |
| EPI_ISL_1795219 | PRONTO ATENDIMENTO UNIDADE SAUDE ADALBERTO ROCHA GUAREI | Instituto Butantan / ESALQ-Piracicaba | Instituto Butantan: Alexander Roberto Precioso, Dimas Tadeu Covas, Sandra Coccuzzo Sampaio, Maria Carolina Elias, José Salvatore Leister Patané, Vincent Louis Viala, Antonio Jorge Martins, Ricardo Haddad, Claudia Renata dos Santos Barros, Elaine Cristina Marqueze, Raul Machado Neto, Debora Botequio Moretti. Centro de Genômica Funcional da ESALQ: Luiz Lehmann Coutinho, Ricardo Augusto Brassaloti, Raquel de Lello Rocha Campos Cassano. NGS Soluções Genômicas: Pilar Drummond Sampaio Corrêa Mariani. FZEA-USP Pirassununga: Mirele Daiana Poleti, Jessika Cristina Chagas Lesbon, Elisangela Chicaroni Mattos, Heidge Fukumasu. USP-Botucatu: Rejane Maria Tommasini Grotto, Jayme A. Souza-Neto, Guilherme Targino Valente, Patricia Akemi Assato, Felipe Allan da Silva da Costa, Bianca Cechetto Carlos. Mendelics: Bibiana Santos, João Paulo Kitajima, Erika Freitas, David Schlesinger. Hemocentro Ribeirão Preto: Simone Kashima, Evandra Strazza Rodrigues, Svetoslav Nanev Slavov, Elaine Vieira dos Santos, Rafael dos Santos Bezerra, Luiz Carlos Junior de Alcantara, Marta Giovanetti, Vagner Fonseca, Flavia Aburjaile, Rodrigo Tocantins Calado. |
| EPI_ISL_1795221 | HOSPITAL REGIONAL DE ITAPETININGA | Instituto Butantan / ESALQ-Piracicaba | Instituto Butantan: Alexander Roberto Precioso, Dimas Tadeu Covas, Sandra Coccuzzo Sampaio, Maria Carolina Elias, José Salvatore Leister Patané, Vincent Louis Viala, Antonio Jorge Martins, Ricardo Haddad, Claudia Renata dos Santos Barros, Elaine Cristina Marqueze, Raul Machado Neto, Debora Botequio Moretti. Centro de Genômica Funcional da ESALQ: Luiz Lehmann Coutinho, Ricardo Augusto Brassaloti, Raquel de Lello Rocha Campos Cassano. NGS Soluções Genômicas: Pilar Drummond Sampaio Corrêa Mariani. FZEA-USP Pirassununga: Mirele Daiana Poleti, Jessika Cristina Chagas |

| | | | |
|---|---|---|---|
| | | | Lesbon, Elisangela Chicaroni Mattos, Heidge Fukumasu. USP-Botucatu: Rejane Maria Tommasini Grotto, Jayme A. Souza-Neto, Guilherme Targino Valente, Patricia Akemi Assato, Felipe Allan da Silva da Costa, Bianca Cechetto Carlos. Mendelics: Bibiana Santos, João Paulo Kitajima, Erika Freitas, David Schlesinger. Hemocentro Ribeirão Preto: Simone Kashima, Evandra Strazza Rodrigues, Svetoslav Nanev Slavov, Elaine Vieira dos Santos, Rafael dos Santos Bezerra, Luiz Carlos Junior de Alcantara, Marta Giovanetti, Vagner Fonseca, Flavia Aburjaile, Rodrigo Tocantins Calado. |
| EPI_ISL_1795231 | UNIDADE DE SAUDE DA FAMILIA JOSE ADALBERTO LELLIS GARCIA | Instituto Butantan / ESALQ-Piracicaba | Instituto Butantan: Alexander Roberto Precioso, Dimas Tadeu Covas, Sandra Coccuzzo Sampaio, Maria Carolina Elias, José Salvatore Leister Patané, Vincent Louis Viala, Antonio Jorge Martins, Ricardo Haddad, Claudia Renata dos Santos Barros, Elaine Cristina Marqueze, Raul Machado Neto, Debora Botequio Moretti. Centro de Genômica Funcional da ESALQ: Luiz Lehmann Coutinho, Ricardo Augusto Brassaloti, Raquel de Lello Rocha Campos Cassano. NGS Soluções Genômicas: Pilar Drummond Sampaio Corrêa Mariani. FZEA-USP Pirassununga: Mirele Daiana Poleti, Jessika Cristina Chagas Lesbon, Elisangela Chicaroni Mattos, Heidge Fukumasu. USP-Botucatu: Rejane Maria Tommasini Grotto, Jayme A. Souza-Neto, Guilherme Targino Valente, Patricia Akemi Assato, Felipe Allan da Silva da Costa, Bianca Cechetto Carlos. Mendelics: Bibiana Santos, João Paulo Kitajima, Erika Freitas, David Schlesinger. Hemocentro Ribeirão Preto: Simone Kashima, Evandra Strazza Rodrigues, Svetoslav Nanev Slavov, Elaine Vieira dos Santos, Rafael dos Santos Bezerra, Luiz Carlos Junior de Alcantara, Marta Giovanetti, Vagner Fonseca, Flavia Aburjaile, Rodrigo Tocantins Calado. |
| EPI_ISL_1795249, EPI_ISL_1795250 | ESF NOVA TANABI II | Instituto Butantan / ESALQ-Piracicaba | Instituto Butantan: Alexander Roberto Precioso, Dimas Tadeu Covas, Sandra Coccuzzo Sampaio, Maria Carolina Elias, José Salvatore Leister Patané, Vincent Louis Viala, Antonio Jorge Martins, Ricardo Haddad, Claudia Renata dos Santos Barros, Elaine Cristina Marqueze, Raul Machado Neto, Debora Botequio Moretti. Centro de Genômica Funcional da ESALQ: Luiz Lehmann Coutinho, Ricardo Augusto Brassaloti, Raquel de Lello Rocha Campos Cassano. NGS Soluções Genômicas: Pilar Drummond Sampaio Corrêa Mariani. FZEA-USP Pirassununga: Mirele Daiana Poleti, Jessika Cristina Chagas Lesbon, Elisangela Chicaroni Mattos, Heidge Fukumasu. USP-Botucatu: Rejane Maria Tommasini Grotto, Jayme A. Souza-Neto, Guilherme Targino Valente, Patricia Akemi Assato, Felipe Allan da Silva da Costa, Bianca Cechetto Carlos. Mendelics: Bibiana Santos, João Paulo Kitajima, Erika Freitas, David Schlesinger. Hemocentro Ribeirão Preto: Simone Kashima, Evandra Strazza Rodrigues, Svetoslav Nanev Slavov, Elaine Vieira dos Santos, Rafael dos Santos Bezerra, Luiz Carlos Junior de Alcantara, Marta Giovanetti, Vagner Fonseca, Flavia Aburjaile, Rodrigo Tocantins Calado. |
| EPI_ISL_1795257 | UBS II DE TANABI MILTON MARTINS PERCHES | Instituto Butantan / ESALQ-Piracicaba | Instituto Butantan: Alexander Roberto Precioso, Dimas Tadeu Covas, Sandra Coccuzzo Sampaio, Maria Carolina Elias, José Salvatore Leister Patané, Vincent Louis Viala, Antonio Jorge Martins, Ricardo Haddad, Claudia Renata dos Santos Barros, Elaine Cristina Marqueze, Raul Machado Neto, Debora Botequio Moretti. Centro de Genômica Funcional da ESALQ: Luiz Lehmann Coutinho, Ricardo Augusto Brassaloti, Raquel de Lello Rocha Campos Cassano. NGS Soluções Genômicas: Pilar Drummond Sampaio Corrêa Mariani. FZEA-USP Pirassununga: Mirele Daiana Poleti, Jessika Cristina Chagas Lesbon, Elisangela Chicaroni Mattos, Heidge Fukumasu. USP-Botucatu: Rejane Maria Tommasini Grotto, Jayme A. Souza-Neto, Guilherme Targino Valente, Patricia Akemi Assato, Felipe Allan da Silva da Costa, Bianca Cechetto Carlos. Mendelics: Bibiana Santos, João Paulo Kitajima, Erika Freitas, David Schlesinger. Hemocentro Ribeirão Preto: Simone Kashima, Evandra Strazza Rodrigues, Svetoslav Nanev Slavov, Elaine Vieira dos Santos, Rafael dos Santos Bezerra, Luiz Carlos Junior de Alcantara, Marta Giovanetti, Vagner Fonseca, Flavia Aburjaile, Rodrigo Tocantins Calado. |
| EPI_ISL_1795338 | HOSPITAL MUNICIPAL REYNALDO GUERRA CAJATI | Instituto Butantan / ESALQ-Piracicaba | Instituto Butantan: Alexander Roberto Precioso, Dimas Tadeu Covas, Sandra Coccuzzo Sampaio, Maria Carolina Elias, José Salvatore Leister Patané, Vincent Louis Viala, Antonio Jorge Martins, Ricardo Haddad, Claudia Renata dos Santos Barros, Elaine Cristina Marqueze, Raul Machado Neto, Debora Botequio Moretti. Centro de Genômica Funcional da ESALQ: Luiz Lehmann Coutinho, Ricardo Augusto Brassaloti, Raquel de Lello Rocha Campos Cassano. NGS Soluções Genômicas: Pilar Drummond Sampaio Corrêa Mariani. FZEA-USP Pirassununga: Mirele Daiana Poleti, Jessika Cristina Chagas Lesbon, Elisangela Chicaroni Mattos, Heidge Fukumasu. USP-Botucatu: Rejane Maria Tommasini Grotto, Jayme A. Souza-Neto, Guilherme Targino Valente, Patricia Akemi Assato, Felipe Allan da Silva da Costa, Bianca Cechetto Carlos. Mendelics: Bibiana Santos, João Paulo Kitajima, Erika Freitas, David Schlesinger. Hemocentro Ribeirão Preto: Simone Kashima, Evandra Strazza Rodrigues, Svetoslav Nanev Slavov, Elaine Vieira dos Santos, Rafael dos Santos Bezerra, Luiz Carlos Junior de Alcantara, Marta Giovanetti, Vagner Fonseca, Flavia Aburjaile, Rodrigo Tocantins Calado. |
| EPI_ISL_1795429 | CENTRO INTEGRADO DE SAUDE | Instituto Butantan / ESALQ-Piracicaba | Instituto Butantan: Alexander Roberto Precioso, Dimas Tadeu Covas, Sandra Coccuzzo Sampaio, Maria Carolina Elias, José Salvatore Leister Patané, Vincent Louis Viala, Antonio Jorge Martins, Ricardo Haddad, Claudia Renata dos Santos Barros, Elaine Cristina Marqueze, Raul Machado Neto, Debora Botequio Moretti. Centro de Genômica Funcional da ESALQ: Luiz Lehmann Coutinho, Ricardo Augusto Brassaloti, Raquel de Lello Rocha Campos Cassano. NGS Soluções Genômicas: Pilar Drummond Sampaio Corrêa Mariani. FZEA-USP Pirassununga: Mirele Daiana Poleti, Jessika Cristina Chagas Lesbon, Elisangela Chicaroni Mattos, Heidge Fukumasu. USP-Botucatu: Rejane Maria Tommasini Grotto, Jayme A. Souza-Neto, Guilherme Targino Valente, Patricia Akemi Assato, Felipe Allan da Silva da Costa, Bianca Cechetto Carlos. Mendelics: Bibiana Santos, João Paulo Kitajima, Erika Freitas, David Schlesinger. Hemocentro Ribeirão Preto: Simone Kashima, Evandra Strazza Rodrigues, Svetoslav Nanev Slavov, Elaine Vieira dos Santos, Rafael dos Santos Bezerra, Luiz Carlos Junior de Alcantara, Marta Giovanetti, Vagner Fonseca, Flavia Aburjaile, Rodrigo Tocantins Calado. |
| EPI_ISL_1821204 | Santa Casa de Cravinhos | Instituto Adolfo Lutz, Interdiciplinary Procedures Center, Strategic Laboratory | Claudio Tavares Sacchi, Claudia Regina Gonçalves, Erica Valessa Ramos Gomes, Karoline Rodrigues Campos, Caio Vinicius Dias Lopes, Leonardo Jose Tadeu de Araujo |
| EPI_ISL_1821205 | UPA Dr Luis Atilio Losi Viana Ribeirao Preto | Instituto Adolfo Lutz, Interdiciplinary Procedures Center, Strategic Laboratory | Claudio Tavares Sacchi, Claudia Regina Gonçalves, Erica Valessa Ramos Gomes, Karoline Rodrigues Campos, Caio Vinicius Dias Lopes, Leonardo Jose Tadeu de Araujo |
| EPI_ISL_1821234 | Instituto Adolfo Lutz - Regional de Marilia | Instituto Adolfo Lutz, Interdiciplinary Procedures Center, Strategic Laboratory | Claudio Tavares Sacchi, Claudia Regina Gonçalves, Erica Valessa Ramos Gomes, Karoline Rodrigues Campos, Caio Vinicius Dias Lopes, Leonardo Jose Tadeu de Araujo |
| EPI_ISL_1966104 | POLICLINICA COVID 19 ITAPETININGA | Instituto Butantan / Mendelics | Instituto Butantan: Dimas Tadeu Covas, Sandra Coccuzzo Sampaio, Maria Carolina Elias, José Salvatore Leister Patané, Vincent Louis Viala, Antonio Jorge Martins, Ricardo Haddad, Claudia Renata dos Santos Barros, Elaine Cristina Marqueze, Raul Machado Neto, Debora Botequio Moretti, Jardelina de Souza Todao Bernardino, Loyze Paola Oliveira de Lima, Luiz Aurelio de Campos Crispin. Centro de Genômica Funcional da ESALQ: Luiz Lehmann Coutinho, Ricardo Augusto Brassaloti, Raquel de Lello Rocha Campos Cassano. NGS Soluções Genômicas: Pilar Drummond Sampaio Corrêa Mariani. FZEA-USP Pirassununga: Mirele Daiana Poleti, Jessika Cristina Chagas Lesbon, Elisangela Chicaroni Mattos, Heidge Fukumasu. USP-Botucatu: Rejane Maria Tommasini Grotto, Jayme A. Souza-Neto, Guilherme Targino Valente, Patricia Akemi Assato, Felipe Allan da Silva da Costa, Bianca Cechetto Carlos. Mendelics: Bibiana Santos, João Paulo Kitajima, Erika Freitas, David Schlesinger. Hemocentro Ribeirão Preto: Simone Kashima, Evandra Strazza Rodrigues, Svetoslav Nanev Slavov, Elaine Vieira dos Santos, Rafael dos Santos Bezerra, Luiz Carlos Junior de Alcantara, Marta Giovanetti, Vagner Fonseca, Flavia Aburjaile, Rodrigo Tocantins Calado. FAMERP-SJRP: Cecília Artico Banho, Lívia Sacchetto, Fábio Sossai Possebon, Leila Sabrina Ullmann, Cíntia Bittar, Guilherme Campos, Helena Lage Ferreira, Jorge A. Petroli Marchesi, Maisa C. Pereira Parra, Marília Moraes, Paula Rahal, Paulo Inacio da Costa, João Pessoa Araújo Jr., Maurício Lacerda Nogueira. Prefeitura de Sao Paulo: Melissa Palmieri. |
| EPI_ISL_1966117 | CS II EGIDIO BRUNHARA MORRO AGUDO | Instituto Butantan / Mendelics | Instituto Butantan: Dimas Tadeu Covas, Sandra Coccuzzo Sampaio, Maria Carolina Elias, José Salvatore Leister Patané, Vincent Louis Viala, Antonio Jorge Martins, Ricardo Haddad, Claudia Renata dos Santos Barros, Elaine Cristina Marqueze, Raul Machado Neto, Debora Botequio Moretti, Jardelina de Souza Todao Bernardino, Loyze Paola Oliveira de Lima, Luiz Aurelio de Campos Crispin. Centro de Genômica Funcional da ESALQ: Luiz Lehmann Coutinho, Ricardo Augusto Brassaloti, Raquel de Lello Rocha Campos Cassano. NGS Soluções Genômicas: Pilar Drummond Sampaio Corrêa Mariani. FZEA-USP Pirassununga: Mirele Daiana Poleti, Jessika Cristina Chagas Lesbon, Elisangela Chicaroni Mattos, Heidge Fukumasu. USP-Botucatu: Rejane Maria Tommasini Grotto, Jayme A. Souza-Neto, Guilherme Targino Valente, Patricia Akemi Assato, Felipe Allan da Silva da Costa, Bianca Cechetto Carlos. Mendelics: Bibiana Santos, João Paulo Kitajima, Erika Freitas, David Schlesinger. Hemocentro Ribeirão Preto: Simone Kashima, Evandra Strazza Rodrigues, Svetoslav Nanev Slavov, Elaine Vieira dos Santos, Rafael dos Santos Bezerra, Luiz Carlos Junior de Alcantara, Marta Giovanetti, Vagner Fonseca, Flavia Aburjaile, Rodrigo Tocantins Calado. FAMERP-SJRP: Cecília Artico Banho, Lívia Sacchetto, Fábio Sossai Possebon, Leila Sabrina Ullmann, Cíntia Bittar, Guilherme Campos, Helena Lage Ferreira, Jorge A. Petroli Marchesi, Maisa C. Pereira Parra, Marília Moraes, Paula Rahal, Paulo Inacio da Costa, João Pessoa Araújo Jr., Maurício Lacerda Nogueira. Prefeitura de Sao Paulo: Melissa Palmieri. |
| EPI_ISL_1966121 | CENTRO DE REFERENCIA DO IDOSO DR HUMBERTO MENDES DE CARVALHO | Instituto Butantan / Mendelics | Instituto Butantan: Dimas Tadeu Covas, Sandra Coccuzzo Sampaio, Maria Carolina Elias, José Salvatore Leister Patané, Vincent Louis Viala, Antonio Jorge Martins, Ricardo Haddad, Claudia Renata dos Santos Barros, Elaine Cristina Marqueze, Raul Machado Neto, Debora Botequio Moretti, Jardelina de Souza Todao Bernardino, Loyze Paola Oliveira de Lima, Luiz Aurelio de Campos Crispin. Centro de Genômica Funcional da ESALQ: Luiz Lehmann Coutinho, Ricardo Augusto Brassaloti, Raquel de Lello Rocha Campos Cassano. NGS Soluções Genômicas: Pilar Drummond Sampaio Corrêa Mariani. FZEA-USP Pirassununga: Mirele Daiana Poleti, Jessika Cristina Chagas Lesbon, Elisangela Chicaroni Mattos, Heidge Fukumasu. USP-Botucatu: Rejane Maria Tommasini Grotto, Jayme A. Souza-Neto, Guilherme Targino Valente, Patricia Akemi Assato, Felipe Allan da Silva da Costa, Bianca Cechetto Carlos. Mendelics: Bibiana Santos, João Paulo Kitajima, Erika Freitas, David Schlesinger. Hemocentro Ribeirão Preto: Simone Kashima, Evandra Strazza Rodrigues, Svetoslav Nanev Slavov, Elaine Vieira dos Santos, Rafael dos Santos Bezerra, Luiz Carlos Junior de Alcantara, Marta Giovanetti, Vagner |

| | | | |
|---|---|---|---|
| | | | Fonseca, Flavia Aburjaile, Rodrigo Tocantins Calado. FAMERP-SJRP: Cecília Artico Banho, Lívia Sacchetto, Fábio Sossai Possebon, Leila Sabrina Ullmann, Cíntia Bittar, Guilherme Campos, Helena Lage Ferreira, Jorge A. Petroli Marchesi, Maisa C. Pereira Parra, Marília Moraes, Paula Rahal, Paulo Inacio da Costa, João Pessoa Araújo Jr., Maurício Lacerda Nogueira. Prefeitura de Sao Paulo: Melissa Palmieri. |
| EPI_ISL_1966173, EPI_ISL_1966388 | UNIDADE DE PRONTO ATENDIMENTO UPA DRA ANA OLIVIA BENTIVOGLIO | Instituto Butantan / Mendelics | Instituto Butantan: Dimas Tadeu Covas, Sandra Coccuzzo Sampaio, Maria Carolina Elias, José Salvatore Leister Patané, Vincent Louis Viala, Antonio Jorge Martins, Ricardo Haddad, Claudia Renata dos Santos Barros, Elaine Cristina Marqueze, Raul Machado Neto, Debora Botequio Moretti, Jardelina de Souza Todao Bernardino, Loyze Paola Oliveira de Lima, Luiz Aurelio de Campos Crispin. Centro de Genômica Funcional da ESALQ: Luiz Lehmann Coutinho, Ricardo Augusto Brassaloti, Raquel de Lello Rocha Campos Cassano. NGS Soluções Genômicas: Pilar Drummond Sampaio Corrêa Mariani. FZEA-USP Pirassununga: Mirele Daiana Poleti, Jessika Cristina Chagas Lesbon, Elisangela Chicaroni Mattos, Heidge Fukumasu. USP-Botucatu: Rejane Maria Tommasini Grotto, Jayme A. Souza-Neto, Guilherme Targino Valente, Patricia Akemi Assato, Felipe Allan da Silva da Costa, Bianca Cechetto Carlos. Mendelics: Bibiana Santos, João Paulo Kitajima, Erika Freitas, David Schlesinger. Hemocentro Ribeirão Preto: Simone Kashima, Evandra Strazza Rodrigues, Svetoslav Nanev Slavov, Elaine Vieira dos Santos, Rafael dos Santos Bezerra, Luiz Carlos Junior de Alcantara, Marta Giovanetti, Vagner Fonseca, Flavia Aburjaile, Rodrigo Tocantins Calado. FAMERP-SJRP: Cecília Artico Banho, Lívia Sacchetto, Fábio Sossai Possebon, Leila Sabrina Ullmann, Cíntia Bittar, Guilherme Campos, Helena Lage Ferreira, Jorge A. Petroli Marchesi, Maisa C. Pereira Parra, Marília Moraes, Paula Rahal, Paulo Inacio da Costa, João Pessoa Araújo Jr., Maurício Lacerda Nogueira. Prefeitura de Sao Paulo: Melissa Palmieri. |
| EPI_ISL_1966410 | HOSPITAL GERAL DE VILA NOVA CACHOEIRINHA SAO PAULO | Instituto Butantan / Mendelics | Instituto Butantan: Dimas Tadeu Covas, Sandra Coccuzzo Sampaio, Maria Carolina Elias, José Salvatore Leister Patané, Vincent Louis Viala, Antonio Jorge Martins, Ricardo Haddad, Claudia Renata dos Santos Barros, Elaine Cristina Marqueze, Raul Machado Neto, Debora Botequio Moretti, Jardelina de Souza Todao Bernardino, Loyze Paola Oliveira de Lima, Luiz Aurelio de Campos Crispin. Centro de Genômica Funcional da ESALQ: Luiz Lehmann Coutinho, Ricardo Augusto Brassaloti, Raquel de Lello Rocha Campos Cassano. NGS Soluções Genômicas: Pilar Drummond Sampaio Corrêa Mariani. FZEA-USP Pirassununga: Mirele Daiana Poleti, Jessika Cristina Chagas Lesbon, Elisangela Chicaroni Mattos, Heidge Fukumasu. USP-Botucatu: Rejane Maria Tommasini Grotto, Jayme A. Souza-Neto, Guilherme Targino Valente, Patricia Akemi Assato, Felipe Allan da Silva da Costa, Bianca Cechetto Carlos. Mendelics: Bibiana Santos, João Paulo Kitajima, Erika Freitas, David Schlesinger. Hemocentro Ribeirão Preto: Simone Kashima, Evandra Strazza Rodrigues, Svetoslav Nanev Slavov, Elaine Vieira dos Santos, Rafael dos Santos Bezerra, Luiz Carlos Junior de Alcantara, Marta Giovanetti, Vagner Fonseca, Flavia Aburjaile, Rodrigo Tocantins Calado. FAMERP-SJRP: Cecília Artico Banho, Lívia Sacchetto, Fábio Sossai Possebon, Leila Sabrina Ullmann, Cíntia Bittar, Guilherme Campos, Helena Lage Ferreira, Jorge A. Petroli Marchesi, Maisa C. Pereira Parra, Marília Moraes, Paula Rahal, Paulo Inacio da Costa, João Pessoa Araújo Jr., Maurício Lacerda Nogueira. Prefeitura de Sao Paulo: Melissa Palmieri. |
| EPI_ISL_1966468 | SECRETARIA MUNICIPAL DE SAUDE | Instituto Butantan / Mendelics | Instituto Butantan: Dimas Tadeu Covas, Sandra Coccuzzo Sampaio, Maria Carolina Elias, José Salvatore Leister Patané, Vincent Louis Viala, Antonio Jorge Martins, Ricardo Haddad, Claudia Renata dos Santos Barros, Elaine Cristina Marqueze, Raul Machado Neto, Debora Botequio Moretti, Jardelina de Souza Todao Bernardino, Loyze Paola Oliveira de Lima, Luiz Aurelio de Campos Crispin. Centro de Genômica Funcional da ESALQ: Luiz Lehmann Coutinho, Ricardo Augusto Brassaloti, Raquel de Lello Rocha Campos Cassano. NGS Soluções Genômicas: Pilar Drummond Sampaio Corrêa Mariani. FZEA-USP Pirassununga: Mirele Daiana Poleti, Jessika Cristina Chagas Lesbon, Elisangela Chicaroni Mattos, Heidge Fukumasu. USP-Botucatu: Rejane Maria Tommasini Grotto, Jayme A. Souza-Neto, Guilherme Targino Valente, Patricia Akemi Assato, Felipe Allan da Silva da Costa, Bianca Cechetto Carlos. Mendelics: Bibiana Santos, João Paulo Kitajima, Erika Freitas, David Schlesinger. Hemocentro Ribeirão Preto: Simone Kashima, Evandra Strazza Rodrigues, Svetoslav Nanev Slavov, Elaine Vieira dos Santos, Rafael dos Santos Bezerra, Luiz Carlos Junior de Alcantara, Marta Giovanetti, Vagner Fonseca, Flavia Aburjaile, Rodrigo Tocantins Calado. FAMERP-SJRP: Cecília Artico Banho, Lívia Sacchetto, Fábio Sossai Possebon, Leila Sabrina Ullmann, Cíntia Bittar, Guilherme Campos, Helena Lage Ferreira, Jorge A. Petroli Marchesi, Maisa C. Pereira Parra, Marília Moraes, Paula Rahal, Paulo Inacio da Costa, João Pessoa Araújo Jr., Maurício Lacerda Nogueira. Prefeitura de Sao Paulo: Melissa Palmieri. |
| EPI_ISL_1966658 | UPA UNIDADE DE PRONTO ATENDIMENTO 24 HORAS BOM JESUS | Instituto Butantan / Mendelics | Instituto Butantan: Dimas Tadeu Covas, Sandra Coccuzzo Sampaio, Maria Carolina Elias, José Salvatore Leister Patané, Vincent Louis Viala, Antonio Jorge Martins, Ricardo Haddad, Claudia Renata dos Santos Barros, Elaine Cristina Marqueze, Raul Machado Neto, Debora Botequio Moretti, Jardelina de Souza Todao Bernardino, Loyze Paola Oliveira de Lima, Luiz Aurelio de Campos Crispin. Centro de Genômica Funcional da ESALQ: Luiz Lehmann Coutinho, Ricardo Augusto Brassaloti, Raquel de Lello Rocha Campos Cassano. NGS Soluções Genômicas: Pilar Drummond Sampaio Corrêa Mariani. FZEA-USP Pirassununga: Mirele Daiana Poleti, Jessika Cristina Chagas Lesbon, Elisangela Chicaroni Mattos, Heidge Fukumasu. USP-Botucatu: Rejane Maria Tommasini Grotto, Jayme A. Souza-Neto, Guilherme Targino Valente, Patricia Akemi Assato, Felipe Allan da Silva da Costa, Bianca Cechetto Carlos. Mendelics: Bibiana Santos, João Paulo Kitajima, Erika Freitas, David Schlesinger. Hemocentro Ribeirão Preto: Simone Kashima, Evandra Strazza Rodrigues, Svetoslav Nanev Slavov, Elaine Vieira dos Santos, Rafael dos Santos Bezerra, Luiz Carlos Junior de Alcantara, Marta Giovanetti, Vagner Fonseca, Flavia Aburjaile, Rodrigo Tocantins Calado. FAMERP-SJRP: Cecília Artico Banho, Lívia Sacchetto, Fábio Sossai Possebon, Leila Sabrina Ullmann, Cíntia Bittar, Guilherme Campos, Helena Lage Ferreira, Jorge A. Petroli Marchesi, Maisa C. Pereira Parra, Marília Moraes, Paula Rahal, Paulo Inacio da Costa, João Pessoa Araújo Jr., Maurício Lacerda Nogueira. Prefeitura de Sao Paulo: Melissa Palmieri. |
| EPI_ISL_1966670 | HOSPITAL MUNICIPAL REYNALDO GUERRA CAJATI | Instituto Butantan / Mendelics | Instituto Butantan: Dimas Tadeu Covas, Sandra Coccuzzo Sampaio, Maria Carolina Elias, José Salvatore Leister Patané, Vincent Louis Viala, Antonio Jorge Martins, Ricardo Haddad, Claudia Renata dos Santos Barros, Elaine Cristina Marqueze, Raul Machado Neto, Debora Botequio Moretti, Jardelina de Souza Todao Bernardino, Loyze Paola Oliveira de Lima, Luiz Aurelio de Campos Crispin. Centro de Genômica Funcional da ESALQ: Luiz Lehmann Coutinho, Ricardo Augusto Brassaloti, Raquel de Lello Rocha Campos Cassano. NGS Soluções Genômicas: Pilar Drummond Sampaio Corrêa Mariani. FZEA-USP Pirassununga: Mirele Daiana Poleti, Jessika Cristina Chagas Lesbon, Elisangela Chicaroni Mattos, Heidge Fukumasu. USP-Botucatu: Rejane Maria Tommasini Grotto, Jayme A. Souza-Neto, Guilherme Targino Valente, Patricia Akemi Assato, Felipe Allan da Silva da Costa, Bianca Cechetto Carlos. Mendelics: Bibiana Santos, João Paulo Kitajima, Erika Freitas, David Schlesinger. Hemocentro Ribeirão Preto: Simone Kashima, Evandra Strazza Rodrigues, Svetoslav Nanev Slavov, Elaine Vieira dos Santos, Rafael dos Santos Bezerra, Luiz Carlos Junior de Alcantara, Marta Giovanetti, Vagner Fonseca, Flavia Aburjaile, Rodrigo Tocantins Calado. FAMERP-SJRP: Cecília Artico Banho, Lívia Sacchetto, Fábio Sossai Possebon, Leila Sabrina Ullmann, Cíntia Bittar, Guilherme Campos, Helena Lage Ferreira, Jorge A. Petroli Marchesi, Maisa C. Pereira Parra, Marília Moraes, Paula Rahal, Paulo Inacio da Costa, João Pessoa Araújo Jr., Maurício Lacerda Nogueira. Prefeitura de Sao Paulo: Melissa Palmieri. |
| EPI_ISL_1966770 | POLICLINICA COVID 19 ITAPETININGA | Instituto Butantan / Mendelics | Instituto Butantan: Dimas Tadeu Covas, Sandra Coccuzzo Sampaio, Maria Carolina Elias, José Salvatore Leister Patané, Vincent Louis Viala, Antonio Jorge Martins, Ricardo Haddad, Claudia Renata dos Santos Barros, Elaine Cristina Marqueze, Raul Machado Neto, Debora Botequio Moretti, Jardelina de Souza Todao Bernardino, Loyze Paola Oliveira de Lima, Luiz Aurelio de Campos Crispin. Centro de Genômica Funcional da ESALQ: Luiz Lehmann Coutinho, Ricardo Augusto Brassaloti, Raquel de Lello Rocha Campos Cassano. NGS Soluções Genômicas: Pilar Drummond Sampaio Corrêa Mariani. FZEA-USP Pirassununga: Mirele Daiana Poleti, Jessika Cristina Chagas Lesbon, Elisangela Chicaroni Mattos, Heidge Fukumasu. USP-Botucatu: Rejane Maria Tommasini Grotto, Jayme A. Souza-Neto, Guilherme Targino Valente, Patricia Akemi Assato, Felipe Allan da Silva da Costa, Bianca Cechetto Carlos. Mendelics: Bibiana Santos, João Paulo Kitajima, Erika Freitas, David Schlesinger. Hemocentro Ribeirão Preto: Simone Kashima, Evandra Strazza Rodrigues, Svetoslav Nanev Slavov, Elaine Vieira dos Santos, Rafael dos Santos Bezerra, Luiz Carlos Junior de Alcantara, Marta Giovanetti, Vagner Fonseca, Flavia Aburjaile, Rodrigo Tocantins Calado. FAMERP-SJRP: Cecília Artico Banho, Lívia Sacchetto, Fábio Sossai Possebon, Leila Sabrina Ullmann, Cíntia Bittar, Guilherme Campos, Helena Lage Ferreira, Jorge A. Petroli Marchesi, Maisa C. Pereira Parra, Marília Moraes, Paula Rahal, Paulo Inacio da Costa, João Pessoa Araújo Jr., Maurício Lacerda Nogueira. Prefeitura de Sao Paulo: Melissa Palmieri. |
| EPI_ISL_1966773, EPI_ISL_1966774 | VIGILANCIA EPIDEMIOLOGICA JARDINOPOLIS SP | Instituto Butantan / Mendelics | Instituto Butantan: Dimas Tadeu Covas, Sandra Coccuzzo Sampaio, Maria Carolina Elias, José Salvatore Leister Patané, Vincent Louis Viala, Antonio Jorge Martins, Ricardo Haddad, Claudia Renata dos Santos Barros, Elaine Cristina Marqueze, Raul Machado Neto, Debora Botequio Moretti, Jardelina de Souza Todao Bernardino, Loyze Paola Oliveira de Lima, Luiz Aurelio de Campos Crispin. Centro de Genômica Funcional da ESALQ: Luiz Lehmann Coutinho, Ricardo Augusto Brassaloti, Raquel de Lello Rocha Campos Cassano. NGS Soluções Genômicas: Pilar Drummond Sampaio Corrêa Mariani. FZEA-USP Pirassununga: Mirele Daiana Poleti, Jessika Cristina Chagas Lesbon, Elisangela Chicaroni Mattos, Heidge Fukumasu. USP-Botucatu: Rejane Maria Tommasini Grotto, Jayme A. Souza-Neto, Guilherme Targino Valente, Patricia Akemi Assato, Felipe Allan da Silva da Costa, Bianca Cechetto Carlos. Mendelics: Bibiana Santos, João Paulo Kitajima, Erika Freitas, David Schlesinger. Hemocentro Ribeirão Preto: Simone Kashima, Evandra Strazza Rodrigues, Svetoslav Nanev Slavov, Elaine Vieira dos Santos, Rafael dos Santos Bezerra, Luiz Carlos Junior de Alcantara, Marta Giovanetti, Vagner Fonseca, Flavia Aburjaile, Rodrigo Tocantins Calado. FAMERP-SJRP: Cecília Artico Banho, Lívia Sacchetto, Fábio Sossai Possebon, Leila Sabrina Ullmann, Cíntia Bittar, Guilherme Campos, Helena Lage Ferreira, Jorge A. Petroli Marchesi, Maisa C. Pereira Parra, Marília Moraes, Paula Rahal, Paulo Inacio da Costa, João Pessoa Araújo Jr., Maurício Lacerda Nogueira. Prefeitura de Sao Paulo: Melissa Palmieri. |
| EPI_ISL_1966814 | CENTRO DE SAUDE II SAO MIGUEL ARCANJO | Instituto Butantan / Mendelics | Instituto Butantan: Dimas Tadeu Covas, Sandra Coccuzzo Sampaio, Maria Carolina Elias, José Salvatore Leister Patané, Vincent Louis Viala, Antonio Jorge Martins, Ricardo Haddad, Claudia Renata dos Santos Barros, Elaine Cristina Marqueze, Raul Machado Neto, Debora Botequio Moretti, Jardelina de Souza Todao Bernardino, Loyze Paola Oliveira de Lima, Luiz Aurelio de Campos Crispin. Centro de Genômica Funcional da ESALQ: Luiz Lehmann |

| | | | |
|---|---|---|---|
| | | | Coutinho, Ricardo Augusto Brassaloti, Raquel de Lello Rocha Campos Cassano. NGS Soluções Genômicas: Pilar Drummond Sampaio Corrêa Mariani. FZEA-USP Pirassununga: Mirele Daiana Poleti, Jessika Cristina Chagas Lesbon, Elisangela Chicaroni Mattos, Heidge Fukumasu. USP-Botucatu: Rejane Maria Tommasini Grotto, Jayme A. Souza-Neto, Guilherme Targino Valente, Patricia Akemi Assato, Felipe Allan da Silva da Costa, Bianca Cechetto Carlos. Mendelics: Bibiana Santos, João Paulo Kitajima, Erika Freitas, David Schlesinger. Hemocentro Ribeirão Preto: Simone Kashima, Evandra Strazza Rodrigues, Svetoslav Nanev Slavov, Elaine Vieira dos Santos, Rafael dos Santos Bezerra, Luiz Carlos Junior de Alcantara, Marta Giovanetti, Vagner Fonseca, Flavia Aburjaile, Rodrigo Tocantins Calado. FAMERP-SJRP: Cecília Artico Banho, Lívia Sacchetto, Fábio Sossai Possebon, Leila Sabrina Ullmann, Cíntia Bittar, Guilherme Campos, Helena Lage Ferreira, Jorge A. Petroli Marchesi, Maisa C. Pereira Parra, Marília Moraes, Paula Rahal, Paulo Inacio da Costa, João Pessoa Araújo Jr., Maurício Lacerda Nogueira. Prefeitura de Sao Paulo: Melissa Palmieri. |
| EPI_ISL_1966816 | AMBULATORIO COVID 19 | Instituto Butantan / Mendelics | Instituto Butantan: Dimas Tadeu Covas, Sandra Coccuzzo Sampaio, Maria Carolina Elias, José Salvatore Leister Patané, Vincent Louis Viala, Antonio Jorge Martins, Ricardo Haddad, Claudia Renata dos Santos Barros, Elaine Cristina Marqueze, Raul Machado Neto, Debora Botequio Moretti, Jardelina de Souza Todao Bernardino, Loyze Paola Oliveira de Lima, Luiz Aurelio de Campos Crispin. Centro de Genômica Funcional da ESALQ: Luiz Lehmann Coutinho, Ricardo Augusto Brassaloti, Raquel de Lello Rocha Campos Cassano. NGS Soluções Genômicas: Pilar Drummond Sampaio Corrêa Mariani. FZEA-USP Pirassununga: Mirele Daiana Poleti, Jessika Cristina Chagas Lesbon, Elisangela Chicaroni Mattos, Heidge Fukumasu. USP-Botucatu: Rejane Maria Tommasini Grotto, Jayme A. Souza-Neto, Guilherme Targino Valente, Patricia Akemi Assato, Felipe Allan da Silva da Costa, Bianca Cechetto Carlos. Mendelics: Bibiana Santos, João Paulo Kitajima, Erika Freitas, David Schlesinger. Hemocentro Ribeirão Preto: Simone Kashima, Evandra Strazza Rodrigues, Svetoslav Nanev Slavov, Elaine Vieira dos Santos, Rafael dos Santos Bezerra, Luiz Carlos Junior de Alcantara, Marta Giovanetti, Vagner Fonseca, Flavia Aburjaile, Rodrigo Tocantins Calado. FAMERP-SJRP: Cecília Artico Banho, Lívia Sacchetto, Fábio Sossai Possebon, Leila Sabrina Ullmann, Cíntia Bittar, Guilherme Campos, Helena Lage Ferreira, Jorge A. Petroli Marchesi, Maisa C. Pereira Parra, Marília Moraes, Paula Rahal, Paulo Inacio da Costa, João Pessoa Araújo Jr., Maurício Lacerda Nogueira. Prefeitura de Sao Paulo: Melissa Palmieri. |
| EPI_ISL_1966846 | HOSPITAL MUNICIPAL REYNALDO GUERRA CAJATI | Instituto Butantan / ESALQ-USP (Piracicaba) | Instituto Butantan: Dimas Tadeu Covas, Sandra Coccuzzo Sampaio, Maria Carolina Elias, José Salvatore Leister Patané, Vincent Louis Viala, Antonio Jorge Martins, Ricardo Haddad, Claudia Renata dos Santos Barros, Elaine Cristina Marqueze, Raul Machado Neto, Debora Botequio Moretti, Jardelina de Souza Todao Bernardino, Loyze Paola Oliveira de Lima, Luiz Aurelio de Campos Crispin. Centro de Genômica Funcional da ESALQ: Luiz Lehmann Coutinho, Ricardo Augusto Brassaloti, Raquel de Lello Rocha Campos Cassano. NGS Soluções Genômicas: Pilar Drummond Sampaio Corrêa Mariani. FZEA-USP Pirassununga: Mirele Daiana Poleti, Jessika Cristina Chagas Lesbon, Elisangela Chicaroni Mattos, Heidge Fukumasu. USP-Botucatu: Rejane Maria Tommasini Grotto, Jayme A. Souza-Neto, Guilherme Targino Valente, Patricia Akemi Assato, Felipe Allan da Silva da Costa, Bianca Cechetto Carlos. Mendelics: Bibiana Santos, João Paulo Kitajima, Erika Freitas, David Schlesinger. Hemocentro Ribeirão Preto: Simone Kashima, Evandra Strazza Rodrigues, Svetoslav Nanev Slavov, Elaine Vieira dos Santos, Rafael dos Santos Bezerra, Luiz Carlos Junior de Alcantara, Marta Giovanetti, Vagner Fonseca, Flavia Aburjaile, Rodrigo Tocantins Calado. FAMERP-SJRP: Cecília Artico Banho, Lívia Sacchetto, Fábio Sossai Possebon, Leila Sabrina Ullmann, Cíntia Bittar, Guilherme Campos, Helena Lage Ferreira, Jorge A. Petroli Marchesi, Maisa C. Pereira Parra, Marília Moraes, Paula Rahal, Paulo Inacio da Costa, João Pessoa Araújo Jr., Maurício Lacerda Nogueira. Prefeitura de Sao Paulo: Melissa Palmieri. |
| EPI_ISL_1966858 | UNIDADE SENTINELA VILA APARECIDA | Instituto Butantan / ESALQ-USP (Piracicaba) | Instituto Butantan: Dimas Tadeu Covas, Sandra Coccuzzo Sampaio, Maria Carolina Elias, José Salvatore Leister Patané, Vincent Louis Viala, Antonio Jorge Martins, Ricardo Haddad, Claudia Renata dos Santos Barros, Elaine Cristina Marqueze, Raul Machado Neto, Debora Botequio Moretti, Jardelina de Souza Todao Bernardino, Loyze Paola Oliveira de Lima, Luiz Aurelio de Campos Crispin. Centro de Genômica Funcional da ESALQ: Luiz Lehmann Coutinho, Ricardo Augusto Brassaloti, Raquel de Lello Rocha Campos Cassano. NGS Soluções Genômicas: Pilar Drummond Sampaio Corrêa Mariani. FZEA-USP Pirassununga: Mirele Daiana Poleti, Jessika Cristina Chagas Lesbon, Elisangela Chicaroni Mattos, Heidge Fukumasu. USP-Botucatu: Rejane Maria Tommasini Grotto, Jayme A. Souza-Neto, Guilherme Targino Valente, Patricia Akemi Assato, Felipe Allan da Silva da Costa, Bianca Cechetto Carlos. Mendelics: Bibiana Santos, João Paulo Kitajima, Erika Freitas, David Schlesinger. Hemocentro Ribeirão Preto: Simone Kashima, Evandra Strazza Rodrigues, Svetoslav Nanev Slavov, Elaine Vieira dos Santos, Rafael dos Santos Bezerra, Luiz Carlos Junior de Alcantara, Marta Giovanetti, Vagner Fonseca, Flavia Aburjaile, Rodrigo Tocantins Calado. FAMERP-SJRP: Cecília Artico Banho, Lívia Sacchetto, Fábio Sossai Possebon, Leila Sabrina Ullmann, Cíntia Bittar, Guilherme Campos, Helena Lage Ferreira, Jorge A. Petroli Marchesi, Maisa C. Pereira Parra, Marília Moraes, Paula Rahal, Paulo Inacio da Costa, João Pessoa Araújo Jr., Maurício Lacerda Nogueira. Prefeitura de Sao Paulo: Melissa Palmieri. |
| EPI_ISL_1966868, EPI_ISL_1966869 | CS DE CATIGUA | Instituto Butantan / ESALQ-USP (Piracicaba) | Instituto Butantan: Dimas Tadeu Covas, Sandra Coccuzzo Sampaio, Maria Carolina Elias, José Salvatore Leister Patané, Vincent Louis Viala, Antonio Jorge Martins, Ricardo Haddad, Claudia Renata dos Santos Barros, Elaine Cristina Marqueze, Raul Machado Neto, Debora Botequio Moretti, Jardelina de Souza Todao Bernardino, Loyze Paola Oliveira de Lima, Luiz Aurelio de Campos Crispin. Centro de Genômica Funcional da ESALQ: Luiz Lehmann Coutinho, Ricardo Augusto Brassaloti, Raquel de Lello Rocha Campos Cassano. NGS Soluções Genômicas: Pilar Drummond Sampaio Corrêa Mariani. FZEA-USP Pirassununga: Mirele Daiana Poleti, Jessika Cristina Chagas Lesbon, Elisangela Chicaroni Mattos, Heidge Fukumasu. USP-Botucatu: Rejane Maria Tommasini Grotto, Jayme A. Souza-Neto, Guilherme Targino Valente, Patricia Akemi Assato, Felipe Allan da Silva da Costa, Bianca Cechetto Carlos. Mendelics: Bibiana Santos, João Paulo Kitajima, Erika Freitas, David Schlesinger. Hemocentro Ribeirão Preto: Simone Kashima, Evandra Strazza Rodrigues, Svetoslav Nanev Slavov, Elaine Vieira dos Santos, Rafael dos Santos Bezerra, Luiz Carlos Junior de Alcantara, Marta Giovanetti, Vagner Fonseca, Flavia Aburjaile, Rodrigo Tocantins Calado. FAMERP-SJRP: Cecília Artico Banho, Lívia Sacchetto, Fábio Sossai Possebon, Leila Sabrina Ullmann, Cíntia Bittar, Guilherme Campos, Helena Lage Ferreira, Jorge A. Petroli Marchesi, Maisa C. Pereira Parra, Marília Moraes, Paula Rahal, Paulo Inacio da Costa, João Pessoa Araújo Jr., Maurício Lacerda Nogueira. Prefeitura de Sao Paulo: Melissa Palmieri. |
| EPI_ISL_1966910, EPI_ISL_1966911 | PR S DA FAMILIA UNIDADE DE SAUDE ADALBERTO ROCHA GUAREI | Instituto Butantan / ESALQ-USP (Piracicaba) | Instituto Butantan: Dimas Tadeu Covas, Sandra Coccuzzo Sampaio, Maria Carolina Elias, José Salvatore Leister Patané, Vincent Louis Viala, Antonio Jorge Martins, Ricardo Haddad, Claudia Renata dos Santos Barros, Elaine Cristina Marqueze, Raul Machado Neto, Debora Botequio Moretti, Jardelina de Souza Todao Bernardino, Loyze Paola Oliveira de Lima, Luiz Aurelio de Campos Crispin. Centro de Genômica Funcional da ESALQ: Luiz Lehmann Coutinho, Ricardo Augusto Brassaloti, Raquel de Lello Rocha Campos Cassano. NGS Soluções Genômicas: Pilar Drummond Sampaio Corrêa Mariani. FZEA-USP Pirassununga: Mirele Daiana Poleti, Jessika Cristina Chagas Lesbon, Elisangela Chicaroni Mattos, Heidge Fukumasu. USP-Botucatu: Rejane Maria Tommasini Grotto, Jayme A. Souza-Neto, Guilherme Targino Valente, Patricia Akemi Assato, Felipe Allan da Silva da Costa, Bianca Cechetto Carlos. Mendelics: Bibiana Santos, João Paulo Kitajima, Erika Freitas, David Schlesinger. Hemocentro Ribeirão Preto: Simone Kashima, Evandra Strazza Rodrigues, Svetoslav Nanev Slavov, Elaine Vieira dos Santos, Rafael dos Santos Bezerra, Luiz Carlos Junior de Alcantara, Marta Giovanetti, Vagner Fonseca, Flavia Aburjaile, Rodrigo Tocantins Calado. FAMERP-SJRP: Cecília Artico Banho, Lívia Sacchetto, Fábio Sossai Possebon, Leila Sabrina Ullmann, Cíntia Bittar, Guilherme Campos, Helena Lage Ferreira, Jorge A. Petroli Marchesi, Maisa C. Pereira Parra, Marília Moraes, Paula Rahal, Paulo Inacio da Costa, João Pessoa Araújo Jr., Maurício Lacerda Nogueira. Prefeitura de Sao Paulo: Melissa Palmieri. |
| EPI_ISL_1966924 | HOSPITAL REGIONAL DE ITAPETININGA | Instituto Butantan / ESALQ-USP (Piracicaba) | Instituto Butantan: Dimas Tadeu Covas, Sandra Coccuzzo Sampaio, Maria Carolina Elias, José Salvatore Leister Patané, Vincent Louis Viala, Antonio Jorge Martins, Ricardo Haddad, Claudia Renata dos Santos Barros, Elaine Cristina Marqueze, Raul Machado Neto, Debora Botequio Moretti, Jardelina de Souza Todao Bernardino, Loyze Paola Oliveira de Lima, Luiz Aurelio de Campos Crispin. Centro de Genômica Funcional da ESALQ: Luiz Lehmann Coutinho, Ricardo Augusto Brassaloti, Raquel de Lello Rocha Campos Cassano. NGS Soluções Genômicas: Pilar Drummond Sampaio Corrêa Mariani. FZEA-USP Pirassununga: Mirele Daiana Poleti, Jessika Cristina Chagas Lesbon, Elisangela Chicaroni Mattos, Heidge Fukumasu. USP-Botucatu: Rejane Maria Tommasini Grotto, Jayme A. Souza-Neto, Guilherme Targino Valente, Patricia Akemi Assato, Felipe Allan da Silva da Costa, Bianca Cechetto Carlos. Mendelics: Bibiana Santos, João Paulo Kitajima, Erika Freitas, David Schlesinger. Hemocentro Ribeirão Preto: Simone Kashima, Evandra Strazza Rodrigues, Svetoslav Nanev Slavov, Elaine Vieira dos Santos, Rafael dos Santos Bezerra, Luiz Carlos Junior de Alcantara, Marta Giovanetti, Vagner Fonseca, Flavia Aburjaile, Rodrigo Tocantins Calado. FAMERP-SJRP: Cecília Artico Banho, Lívia Sacchetto, Fábio Sossai Possebon, Leila Sabrina Ullmann, Cíntia Bittar, Guilherme Campos, Helena Lage Ferreira, Jorge A. Petroli Marchesi, Maisa C. Pereira Parra, Marília Moraes, Paula Rahal, Paulo Inacio da Costa, João Pessoa Araújo Jr., Maurício Lacerda Nogueira. Prefeitura de Sao Paulo: Melissa Palmieri. |
| EPI_ISL_1966969 | CS II EGIDIO BRUNHARA MORRO AGUDO | Instituto Butantan / ESALQ-USP (Piracicaba) | Instituto Butantan: Dimas Tadeu Covas, Sandra Coccuzzo Sampaio, Maria Carolina Elias, José Salvatore Leister Patané, Vincent Louis Viala, Antonio Jorge Martins, Ricardo Haddad, Claudia Renata dos Santos Barros, Elaine Cristina Marqueze, Raul Machado Neto, Debora Botequio Moretti, Jardelina de Souza Todao Bernardino, Loyze Paola Oliveira de Lima, Luiz Aurelio de Campos Crispin. Centro de Genômica Funcional da ESALQ: Luiz Lehmann Coutinho, Ricardo Augusto Brassaloti, Raquel de Lello Rocha Campos Cassano. NGS Soluções Genômicas: Pilar Drummond Sampaio Corrêa Mariani. FZEA-USP Pirassununga: Mirele Daiana Poleti, Jessika Cristina Chagas Lesbon, Elisangela Chicaroni Mattos, Heidge Fukumasu. USP-Botucatu: Rejane Maria Tommasini Grotto, Jayme A. Souza-Neto, Guilherme Targino Valente, Patricia Akemi Assato, Felipe Allan da Silva da Costa, Bianca Cechetto Carlos. Mendelics: Bibiana Santos, João Paulo Kitajima, Erika Freitas, David Schlesinger. Hemocentro Ribeirão Preto: Simone Kashima, Evandra Strazza Rodrigues, Svetoslav Nanev Slavov, Elaine Vieira dos Santos, Rafael dos Santos Bezerra, Luiz Carlos Junior de Alcantara, Marta Giovanetti, Vagner |

| | | | |
|---|---|---|---|
| | | | Fonseca, Flavia Aburjaile, Rodrigo Tocantins Calado. FAMERP-SJRP: Cecília Artico Banho, Lívia Sacchetto, Fábio Sossai Possebon, Leila Sabrina Ullmann, Cíntia Bittar, Guilherme Campos, Helena Lage Ferreira, Jorge A. Petroli Marchesi, Maisa C. Pereira Parra, Marília Moraes, Paula Rahal, Paulo Inacio da Costa, João Pessoa Araújo Jr., Maurício Lacerda Nogueira. Prefeitura de Sao Paulo: Melissa Palmieri. |
| EPI_ISL_1966995 | UBS III DE PARIQUERA ACU PARIQUERA ACU | Instituto Butantan / ESALQ-USP (Piracicaba) | Instituto Butantan: Dimas Tadeu Covas, Sandra Coccuzzo Sampaio, Maria Carolina Elias, José Salvatore Leister Patané, Vincent Louis Viala, Antonio Jorge Martins, Ricardo Haddad, Claudia Renata dos Santos Barros, Elaine Cristina Marqueze, Raul Machado Neto, Debora Botequio Moretti, Jardelina de Souza Todao Bernardino, Loyze Paola Oliveira de Lima, Luiz Aurelio de Campos Crispin. Centro de Genômica Funcional da ESALQ: Luiz Lehmann Coutinho, Ricardo Augusto Brassaloti, Raquel de Lello Rocha Campos Cassano. NGS Soluções Genômicas: Pilar Drummond Sampaio Corrêa Mariani. FZEA-USP Pirassununga: Mirele Daiana Poleti, Jessika Cristina Chagas Lesbon, Elisangela Chicaroni Mattos, Heidge Fukumasu. USP-Botucatu: Rejane Maria Tommasini Grotto, Jayme A. Souza-Neto, Guilherme Targino Valente, Patricia Akemi Assato, Felipe Allan da Silva da Costa, Bianca Cechetto Carlos. Mendelics: Bibiana Santos, João Paulo Kitajima, Erika Freitas, David Schlesinger. Hemocentro Ribeirão Preto: Simone Kashima, Evandra Strazza Rodrigues, Svetoslav Nanev Slavov, Elaine Vieira dos Santos, Rafael dos Santos Bezerra, Luiz Carlos Junior de Alcantara, Marta Giovanetti, Vagner Fonseca, Flavia Aburjaile, Rodrigo Tocantins Calado. FAMERP-SJRP: Cecília Artico Banho, Lívia Sacchetto, Fábio Sossai Possebon, Leila Sabrina Ullmann, Cíntia Bittar, Guilherme Campos, Helena Lage Ferreira, Jorge A. Petroli Marchesi, Maisa C. Pereira Parra, Marília Moraes, Paula Rahal, Paulo Inacio da Costa, João Pessoa Araújo Jr., Maurício Lacerda Nogueira. Prefeitura de Sao Paulo: Melissa Palmieri. |
| EPI_ISL_1967061 | CENTRO DE SAUDE II SAO MIGUEL ARCANJO | Instituto Butantan / ESALQ-USP (Piracicaba) | Instituto Butantan: Dimas Tadeu Covas, Sandra Coccuzzo Sampaio, Maria Carolina Elias, José Salvatore Leister Patané, Vincent Louis Viala, Antonio Jorge Martins, Ricardo Haddad, Claudia Renata dos Santos Barros, Elaine Cristina Marqueze, Raul Machado Neto, Debora Botequio Moretti, Jardelina de Souza Todao Bernardino, Loyze Paola Oliveira de Lima, Luiz Aurelio de Campos Crispin. Centro de Genômica Funcional da ESALQ: Luiz Lehmann Coutinho, Ricardo Augusto Brassaloti, Raquel de Lello Rocha Campos Cassano. NGS Soluções Genômicas: Pilar Drummond Sampaio Corrêa Mariani. FZEA-USP Pirassununga: Mirele Daiana Poleti, Jessika Cristina Chagas Lesbon, Elisangela Chicaroni Mattos, Heidge Fukumasu. USP-Botucatu: Rejane Maria Tommasini Grotto, Jayme A. Souza-Neto, Guilherme Targino Valente, Patricia Akemi Assato, Felipe Allan da Silva da Costa, Bianca Cechetto Carlos. Mendelics: Bibiana Santos, João Paulo Kitajima, Erika Freitas, David Schlesinger. Hemocentro Ribeirão Preto: Simone Kashima, Evandra Strazza Rodrigues, Svetoslav Nanev Slavov, Elaine Vieira dos Santos, Rafael dos Santos Bezerra, Luiz Carlos Junior de Alcantara, Marta Giovanetti, Vagner Fonseca, Flavia Aburjaile, Rodrigo Tocantins Calado. FAMERP-SJRP: Cecília Artico Banho, Lívia Sacchetto, Fábio Sossai Possebon, Leila Sabrina Ullmann, Cíntia Bittar, Guilherme Campos, Helena Lage Ferreira, Jorge A. Petroli Marchesi, Maisa C. Pereira Parra, Marília Moraes, Paula Rahal, Paulo Inacio da Costa, João Pessoa Araújo Jr., Maurício Lacerda Nogueira. Prefeitura de Sao Paulo: Melissa Palmieri. |
| EPI_ISL_1967149 | LABORATORIO MUNICIPAL DE SUZANO | Instituto Butantan / ESALQ-USP (Piracicaba) | Instituto Butantan: Dimas Tadeu Covas, Sandra Coccuzzo Sampaio, Maria Carolina Elias, José Salvatore Leister Patané, Vincent Louis Viala, Antonio Jorge Martins, Ricardo Haddad, Claudia Renata dos Santos Barros, Elaine Cristina Marqueze, Raul Machado Neto, Debora Botequio Moretti, Jardelina de Souza Todao Bernardino, Loyze Paola Oliveira de Lima, Luiz Aurelio de Campos Crispin. Centro de Genômica Funcional da ESALQ: Luiz Lehmann Coutinho, Ricardo Augusto Brassaloti, Raquel de Lello Rocha Campos Cassano. NGS Soluções Genômicas: Pilar Drummond Sampaio Corrêa Mariani. FZEA-USP Pirassununga: Mirele Daiana Poleti, Jessika Cristina Chagas Lesbon, Elisangela Chicaroni Mattos, Heidge Fukumasu. USP-Botucatu: Rejane Maria Tommasini Grotto, Jayme A. Souza-Neto, Guilherme Targino Valente, Patricia Akemi Assato, Felipe Allan da Silva da Costa, Bianca Cechetto Carlos. Mendelics: Bibiana Santos, João Paulo Kitajima, Erika Freitas, David Schlesinger. Hemocentro Ribeirão Preto: Simone Kashima, Evandra Strazza Rodrigues, Svetoslav Nanev Slavov, Elaine Vieira dos Santos, Rafael dos Santos Bezerra, Luiz Carlos Junior de Alcantara, Marta Giovanetti, Vagner Fonseca, Flavia Aburjaile, Rodrigo Tocantins Calado. FAMERP-SJRP: Cecília Artico Banho, Lívia Sacchetto, Fábio Sossai Possebon, Leila Sabrina Ullmann, Cíntia Bittar, Guilherme Campos, Helena Lage Ferreira, Jorge A. Petroli Marchesi, Maisa C. Pereira Parra, Marília Moraes, Paula Rahal, Paulo Inacio da Costa, João Pessoa Araújo Jr., Maurício Lacerda Nogueira. Prefeitura de Sao Paulo: Melissa Palmieri. |
| EPI_ISL_1967180 | HOSPITAL DE CAMPANHA COVID 19 MUNICIPIO DE TAUBATE | Instituto Butantan / ESALQ-USP (Piracicaba) | Instituto Butantan: Dimas Tadeu Covas, Sandra Coccuzzo Sampaio, Maria Carolina Elias, José Salvatore Leister Patané, Vincent Louis Viala, Antonio Jorge Martins, Ricardo Haddad, Claudia Renata dos Santos Barros, Elaine Cristina Marqueze, Raul Machado Neto, Debora Botequio Moretti, Jardelina de Souza Todao Bernardino, Loyze Paola Oliveira de Lima, Luiz Aurelio de Campos Crispin. Centro de Genômica Funcional da ESALQ: Luiz Lehmann Coutinho, Ricardo Augusto Brassaloti, Raquel de Lello Rocha Campos Cassano. NGS Soluções Genômicas: Pilar Drummond Sampaio Corrêa Mariani. FZEA-USP Pirassununga: Mirele Daiana Poleti, Jessika Cristina Chagas Lesbon, Elisangela Chicaroni Mattos, Heidge Fukumasu. USP-Botucatu: Rejane Maria Tommasini Grotto, Jayme A. Souza-Neto, Guilherme Targino Valente, Patricia Akemi Assato, Felipe Allan da Silva da Costa, Bianca Cechetto Carlos. Mendelics: Bibiana Santos, João Paulo Kitajima, Erika Freitas, David Schlesinger. Hemocentro Ribeirão Preto: Simone Kashima, Evandra Strazza Rodrigues, Svetoslav Nanev Slavov, Elaine Vieira dos Santos, Rafael dos Santos Bezerra, Luiz Carlos Junior de Alcantara, Marta Giovanetti, Vagner Fonseca, Flavia Aburjaile, Rodrigo Tocantins Calado. FAMERP-SJRP: Cecília Artico Banho, Lívia Sacchetto, Fábio Sossai Possebon, Leila Sabrina Ullmann, Cíntia Bittar, Guilherme Campos, Helena Lage Ferreira, Jorge A. Petroli Marchesi, Maisa C. Pereira Parra, Marília Moraes, Paula Rahal, Paulo Inacio da Costa, João Pessoa Araújo Jr., Maurício Lacerda Nogueira. Prefeitura de Sao Paulo: Melissa Palmieri. |
| EPI_ISL_1967244 | LABORATORIO MUNICIPAL DE ANALISES CLINICAS DE ITANHAEM | Instituto Butantan / Mendelics | Instituto Butantan: Dimas Tadeu Covas, Sandra Coccuzzo Sampaio, Maria Carolina Elias, José Salvatore Leister Patané, Vincent Louis Viala, Antonio Jorge Martins, Ricardo Haddad, Claudia Renata dos Santos Barros, Elaine Cristina Marqueze, Raul Machado Neto, Debora Botequio Moretti, Jardelina de Souza Todao Bernardino, Loyze Paola Oliveira de Lima, Luiz Aurelio de Campos Crispin. Centro de Genômica Funcional da ESALQ: Luiz Lehmann Coutinho, Ricardo Augusto Brassaloti, Raquel de Lello Rocha Campos Cassano. NGS Soluções Genômicas: Pilar Drummond Sampaio Corrêa Mariani. FZEA-USP Pirassununga: Mirele Daiana Poleti, Jessika Cristina Chagas Lesbon, Elisangela Chicaroni Mattos, Heidge Fukumasu. USP-Botucatu: Rejane Maria Tommasini Grotto, Jayme A. Souza-Neto, Guilherme Targino Valente, Patricia Akemi Assato, Felipe Allan da Silva da Costa, Bianca Cechetto Carlos. Mendelics: Bibiana Santos, João Paulo Kitajima, Erika Freitas, David Schlesinger. Hemocentro Ribeirão Preto: Simone Kashima, Evandra Strazza Rodrigues, Svetoslav Nanev Slavov, Elaine Vieira dos Santos, Rafael dos Santos Bezerra, Luiz Carlos Junior de Alcantara, Marta Giovanetti, Vagner Fonseca, Flavia Aburjaile, Rodrigo Tocantins Calado. FAMERP-SJRP: Cecília Artico Banho, Lívia Sacchetto, Fábio Sossai Possebon, Leila Sabrina Ullmann, Cíntia Bittar, Guilherme Campos, Helena Lage Ferreira, Jorge A. Petroli Marchesi, Maisa C. Pereira Parra, Marília Moraes, Paula Rahal, Paulo Inacio da Costa, João Pessoa Araújo Jr., Maurício Lacerda Nogueira. Prefeitura de Sao Paulo: Melissa Palmieri. |
| EPI_ISL_2003128 | Instituto Adolfo Lutz Central | Instituto Adolfo Lutz, Interdiciplinary Procedures Center, Strategic Laboratory | Claudio Tavares Sacchi, Claudia Regina Gonçalves, Erica Valessa Ramos Gomes, Karoline Rodrigues Campos, Caio Vinicius Dias Lopes, Leonardo Jose Tadeu de Araujo |
| EPI_ISL_2017265, EPI_ISL_2017316 | HLAGYN - Laboratorio de Imunologia de Transplantes de Goias | HLAGYN - Laboratorio de Imunologia de Transplantes de Goias | Fernando Antonio Vinhal dos Santos, Erika Lopes Rocha Batista, Alessandro Leonardo Alvares Magalhaes, Raphael Bessa Parmigiane, Frederico Rodrigues Vinhal, Sabrina Sara Moreira Duarte, Danielle de Paiva Rezende, Lucas Carlos Gomes Pereira, Paola Cristina Resende Silva |
| EPI_ISL_2017432, EPI_ISL_2102521 | HLAGYN - Laboratorio de Imunologia de Transplantes de Goias | HLAGYN - Laboratorio de Imunologia de Transplantes de Goias | Fernando Antonio Vinhal dos Santos, Erika Lopes Rocha Batista, Alessandro Leonardo Alvares Magalhaes, Frederico Rodrigues Vinhal, Sabrina Sara Moreira Duarte, Danielle de Paiva Rezende, Lucas Carlos Gomes Pereira, Paola Cristina Resende Silva |
| EPI_ISL_2170973 | UPA DE BEBEDOURO | Instituto Butantan / Mendelics | Instituto Butantan: Dimas Tadeu Covas, Sandra Coccuzzo Sampaio, Maria Carolina Elias, José Salvatore Leister Patané, Vincent Louis Viala, Antonio Jorge Martins, Ricardo Haddad, Claudia Renata dos Santos Barros, Elaine Cristina Marqueze, Raul Machado Neto, Debora Botequio Moretti, Jardelina de Souza Todao Bernardino, Loyze Paola Oliveira de Lima, Luiz Aurelio de Campos Crispin. Centro de Genômica Funcional da ESALQ: Luiz Lehmann Coutinho, Ricardo Augusto Brassaloti, Raquel de Lello Rocha Campos Cassano. NGS Soluções Genômicas: Pilar Drummond Sampaio Corrêa Mariani. FZEA-USP Pirassununga: Mirele Daiana Poleti, Jessika Cristina Chagas Lesbon, Elisangela Chicaroni Mattos, Heidge Fukumasu. USP-Botucatu: Rejane Maria Tommasini Grotto, Jayme A. Souza-Neto, Guilherme Targino Valente, Patricia Akemi Assato, Felipe Allan da Silva da Costa, Bianca Cechetto Carlos. Mendelics: Bibiana Santos, João Paulo Kitajima, Erika Freitas, David Schlesinger. Hemocentro Ribeirão Preto: Simone Kashima, Evandra Strazza Rodrigues, Svetoslav Nanev Slavov, Elaine Vieira dos Santos, Rafael dos Santos Bezerra, Luiz Carlos Junior de Alcantara, Marta Giovanetti, Vagner Fonseca, Flavia Aburjaile, Rodrigo Tocantins Calado. FAMERP-SJRP: Cecília Artico Banho, Lívia Sacchetto, Fábio Sossai Possebon, Leila Sabrina Ullmann, Cíntia Bittar, Guilherme Campos, Helena Lage Ferreira, Jorge A. Petroli Marchesi, Maisa C. Pereira Parra, Marília Moraes, Paula Rahal, Paulo Inacio da Costa, João Pessoa Araújo Jr., Maurício Lacerda Nogueira. Prefeitura de Sao Paulo: Melissa Palmieri. |
| EPI_ISL_2187857, EPI_ISL_2187882, EPI_ISL_2187981 | HLAGYN - Laboratorio de Imunologia de Transplantes de Goias | HLAGYN - Laboratorio de Imunologia de Transplantes de Goias | Fernando Antonio Vinhal dos Santos, Erika Lopes Rocha Batista, Alessandro Leonardo Alvares Magalhaes, Frederico Rodrigues Vinhal, Sabrina Sara Moreira Duarte, Lucas Carlos Gomes Pereira, Daniel Ferreira de Sousa |
| EPI_ISL_2208958 | VIGILANCIA EPIDEMIOLOGICA JARDINOPOLIS SP | Instituto Butantan / Mendelics | Dimas Tadeu Covas, Antonio Jorge Martins, Claudia Renata dos Santos Barros, David Schlesinger, Debora Botequio Moretti, Elaine Cristina Marqueze, Elaine Vieira Santos, Evandra Strazza Rodrigues, Heidge Fukumasu, Jayme Augusto de Souza-Neto, José Salvatore Leister Patané, Luiz Alcantara, Luiz Lehmann Coutinho, Maria Carolina Elias, Maurício Lacerda Nogueira, Rafael dos Santos Bezerra, Raul Machado Neto, Rejane Maria Tommasini Grotto, Ricardo Haddad, Sandra Coccuzzo Sampaio Vessoni, Simone Kashima, Svetoslav Nanev Slavov, Vincent Louis Viala |

| | | | |
|---|---|---|---|
| EPI_ISL_2209692 | LABORATORIO DR PAULO EMILIO DALESSANDRO PINDAMONHANGABA | Instituto Butantan / Mendelics | Dimas Tadeu Covas, Antonio Jorge Martins, Claudia Renata dos Santos Barros, David Schlesinger, Debora Botequio Moretti, Elaine Cristina Marqueze, Elaine Vieira Santos, Evandra Strazza Rodrigues, Heidge Fukumasu, Jayme Augusto de Souza-Neto, José Salvatore Leister Patané, Luiz Alcantara, Luiz Lehmann Coutinho, Maria Carolina Elias, Maurício Lacerda Nogueira, Rafael dos Santos Bezerra, Raul Machado Neto, Rejane Maria Tommasini Grotto, Ricardo Haddad, Sandra Coccuzzo Sampaio Vessoni, Simone Kashima, Svetoslav Nanev Slavov, Vincent Louis Viala |
| EPI_ISL_2274049 | Laboratorio Central de Saude Publica do Estado de Alagoas (LACEN/AL) | Laboratory of Respiratory Viruses and Measles, Oswaldo Cruz Institute, FIOCRUZ | Paola Resende, Luciana Appolinario, Fernando Motta, Anna Carolina Paixao, Ana Carolina Mendonca, Alice Sampaio Rocha, Taina Venas, Elisa Cavalcante Pereira, Renata Serrano Lopes, Anderson Brandao Leite, Marilda Siqueira on behalf of the Fiocruz COVID-19 Genomic Surveillance Network |
| EPI_ISL_2295405 | Instituto de Biotecnologia - UNESP-Botucatu-SP | Instituto de Biotecnologia - UNESP-Botucatu-SP | Fábio Sossai Possebon; Leila Sabrina Ullmann; Cecília Artico Banho; Síntia Bittar; Guilherme Campos; Helena Lage Ferreira; Jorge A. Petroli Marchesi; Lívia Sacchetto; Maisa C. Pereira Parra; Marília Moraes; Maurício L. Nogueira; Paula Rahal; Paulo Inacio da Costa; João Pessoa Araújo Jr. |
| EPI_ISL_2298842 | Laboratório Central de Saúde Pública do Maranhão | Coordenação Geral de Laboratórios de Saúde Pública (CGLAB/DAEVS/SVS/MS) | Vagner Fonseca, et al. |
| EPI_ISL_2344823 | CS DE CATIGUA | Instituto Butantan / ESALQ-Piracicaba | Dimas Tadeu Covas, Antonio Jorge Martins, Claudia Renata dos Santos Barros, David Schlesinger, Debora Botequio Moretti, Elaine Cristina Marqueze, Elaine Vieira Santos, Evandra Strazza Rodrigues, Heidge Fukumasu, Jayme Augusto de Souza-Neto, José Salvatore Leister Patané, Luiz Alcantara, Luiz Lehmann Coutinho, Maria Carolina Elias, Maurício Lacerda Nogueira, Rafael dos Santos Bezerra, Raul Machado Neto, Rejane Maria Tommasini Grotto, Ricardo Haddad, Sandra Coccuzzo Sampaio Vessoni, Simone Kashima, Svetoslav Nanev Slavov, Vincent Louis Viala |
| EPI_ISL_2344997 | HOSPITAL DE CAMPANHA COVID 19 MUNICIPIO DE TAUBATE | Instituto Butantan / ESALQ-Piracicaba | Dimas Tadeu Covas, Antonio Jorge Martins, Claudia Renata dos Santos Barros, David Schlesinger, Debora Botequio Moretti, Elaine Cristina Marqueze, Elaine Vieira Santos, Evandra Strazza Rodrigues, Heidge Fukumasu, Jayme Augusto de Souza-Neto, José Salvatore Leister Patané, Luiz Alcantara, Luiz Lehmann Coutinho, Maria Carolina Elias, Maurício Lacerda Nogueira, Rafael dos Santos Bezerra, Raul Machado Neto, Rejane Maria Tommasini Grotto, Ricardo Haddad, Sandra Coccuzzo Sampaio Vessoni, Simone Kashima, Svetoslav Nanev Slavov, Vincent Louis Viala |
| EPI_ISL_2345297 | LABORATORIO MUNICIPAL DE PIRACICABA | Instituto Butantan / ESALQ-Piracicaba | Dimas Tadeu Covas, Antonio Jorge Martins, Claudia Renata dos Santos Barros, David Schlesinger, Debora Botequio Moretti, Elaine Cristina Marqueze, Elaine Vieira Santos, Evandra Strazza Rodrigues, Heidge Fukumasu, Jayme Augusto de Souza-Neto, José Salvatore Leister Patané, Luiz Alcantara, Luiz Lehmann Coutinho, Maria Carolina Elias, Maurício Lacerda Nogueira, Rafael dos Santos Bezerra, Raul Machado Neto, Rejane Maria Tommasini Grotto, Ricardo Haddad, Sandra Coccuzzo Sampaio Vessoni, Simone Kashima, Svetoslav Nanev Slavov, Vincent Louis Viala |
| EPI_ISL_2345415 | USF SALERNO | Instituto Butantan / ESALQ-Piracicaba | Dimas Tadeu Covas, Antonio Jorge Martins, Claudia Renata dos Santos Barros, David Schlesinger, Debora Botequio Moretti, Elaine Cristina Marqueze, Elaine Vieira Santos, Evandra Strazza Rodrigues, Heidge Fukumasu, Jayme Augusto de Souza-Neto, José Salvatore Leister Patané, Luiz Alcantara, Luiz Lehmann Coutinho, Maria Carolina Elias, Maurício Lacerda Nogueira, Rafael dos Santos Bezerra, Raul Machado Neto, Rejane Maria Tommasini Grotto, Ricardo Haddad, Sandra Coccuzzo Sampaio Vessoni, Simone Kashima, Svetoslav Nanev Slavov, Vincent Louis Viala |
| EPI_ISL_2345449 | PRONTO ATENDIMENTO UNIDADE SAUDE ADALBERTO ROCHA GUAREI | Instituto Butantan / ESALQ-Piracicaba | Dimas Tadeu Covas, Antonio Jorge Martins, Claudia Renata dos Santos Barros, David Schlesinger, Debora Botequio Moretti, Elaine Cristina Marqueze, Elaine Vieira Santos, Evandra Strazza Rodrigues, Heidge Fukumasu, Jayme Augusto de Souza-Neto, José Salvatore Leister Patané, Luiz Alcantara, Luiz Lehmann Coutinho, Maria Carolina Elias, Maurício Lacerda Nogueira, Rafael dos Santos Bezerra, Raul Machado Neto, Rejane Maria Tommasini Grotto, Ricardo Haddad, Sandra Coccuzzo Sampaio Vessoni, Simone Kashima, Svetoslav Nanev Slavov, Vincent Louis Viala |
| EPI_ISL_2345451 | HOSPITAL REGIONAL DE ITAPETININGA | Instituto Butantan / ESALQ-Piracicaba | Dimas Tadeu Covas, Antonio Jorge Martins, Claudia Renata dos Santos Barros, David Schlesinger, Debora Botequio Moretti, Elaine Cristina Marqueze, Elaine Vieira Santos, Evandra Strazza Rodrigues, Heidge Fukumasu, Jayme Augusto de Souza-Neto, José Salvatore Leister Patané, Luiz Alcantara, Luiz Lehmann Coutinho, Maria Carolina Elias, Maurício Lacerda Nogueira, Rafael dos Santos Bezerra, Raul Machado Neto, Rejane Maria Tommasini Grotto, Ricardo Haddad, Sandra Coccuzzo Sampaio Vessoni, Simone Kashima, Svetoslav Nanev Slavov, Vincent Louis Viala |
| EPI_ISL_2345463 | UNIDADE DE SAUDE DA FAMILIA JOSE ADALBERTO LELLIS GARCIA | Instituto Butantan / ESALQ-Piracicaba | Dimas Tadeu Covas, Antonio Jorge Martins, Claudia Renata dos Santos Barros, David Schlesinger, Debora Botequio Moretti, Elaine Cristina Marqueze, Elaine Vieira Santos, Evandra Strazza Rodrigues, Heidge Fukumasu, Jayme Augusto de Souza-Neto, José Salvatore Leister Patané, Luiz Alcantara, Luiz Lehmann Coutinho, Maria Carolina Elias, Maurício Lacerda Nogueira, Rafael dos Santos Bezerra, Raul Machado Neto, Rejane Maria Tommasini Grotto, Ricardo Haddad, Sandra Coccuzzo Sampaio Vessoni, Simone Kashima, Svetoslav Nanev Slavov, Vincent Louis Viala |
| EPI_ISL_2345483, EPI_ISL_2345484 | ESF NOVA TANABI II | Instituto Butantan / ESALQ-Piracicaba | Dimas Tadeu Covas, Antonio Jorge Martins, Claudia Renata dos Santos Barros, David Schlesinger, Debora Botequio Moretti, Elaine Cristina Marqueze, Elaine Vieira Santos, Evandra Strazza Rodrigues, Heidge Fukumasu, Jayme Augusto de Souza-Neto, José Salvatore Leister Patané, Luiz Alcantara, Luiz Lehmann Coutinho, Maria Carolina Elias, Maurício Lacerda Nogueira, Rafael dos Santos Bezerra, Raul Machado Neto, Rejane Maria Tommasini Grotto, Ricardo Haddad, Sandra Coccuzzo Sampaio Vessoni, Simone Kashima, Svetoslav Nanev Slavov, Vincent Louis Viala |
| EPI_ISL_2345491 | UBS II DE TANABI MILTON MARTINS PERCHES | Instituto Butantan / ESALQ-Piracicaba | Dimas Tadeu Covas, Antonio Jorge Martins, Claudia Renata dos Santos Barros, David Schlesinger, Debora Botequio Moretti, Elaine Cristina Marqueze, Elaine Vieira Santos, Evandra Strazza Rodrigues, Heidge Fukumasu, Jayme Augusto de Souza-Neto, José Salvatore Leister Patané, Luiz Alcantara, Luiz Lehmann Coutinho, Maria Carolina Elias, Maurício Lacerda Nogueira, Rafael dos Santos Bezerra, Raul Machado Neto, Rejane Maria Tommasini Grotto, Ricardo Haddad, Sandra Coccuzzo Sampaio Vessoni, Simone Kashima, Svetoslav Nanev Slavov, Vincent Louis Viala |
| EPI_ISL_2345604 | HOSPITAL MUNICIPAL REYNALDO GUERRA CAJATI | Instituto Butantan / ESALQ-Piracicaba | Dimas Tadeu Covas, Antonio Jorge Martins, Claudia Renata dos Santos Barros, David Schlesinger, Debora Botequio Moretti, Elaine Cristina Marqueze, Elaine Vieira Santos, Evandra Strazza Rodrigues, Heidge Fukumasu, Jayme Augusto de Souza-Neto, José Salvatore Leister Patané, Luiz Alcantara, Luiz Lehmann Coutinho, Maria Carolina Elias, Maurício Lacerda Nogueira, Rafael dos Santos Bezerra, Raul Machado Neto, Rejane Maria Tommasini Grotto, Ricardo Haddad, Sandra Coccuzzo Sampaio Vessoni, Simone Kashima, Svetoslav Nanev Slavov, Vincent Louis Viala |
| EPI_ISL_2345669 | CENTRO DE REFERENCIA DO IDOSO DR HUMBERTO MENDES DE CARVALHO | Instituto Butantan / Mendelics | Dimas Tadeu Covas, Antonio Jorge Martins, Claudia Renata dos Santos Barros, David Schlesinger, Debora Botequio Moretti, Elaine Cristina Marqueze, Elaine Vieira Santos, Evandra Strazza Rodrigues, Heidge Fukumasu, Jayme Augusto de Souza-Neto, José Salvatore Leister Patané, Luiz Alcantara, Luiz Lehmann Coutinho, Maria Carolina Elias, Maurício Lacerda Nogueira, Rafael dos Santos Bezerra, Raul Machado Neto, Rejane Maria Tommasini Grotto, Ricardo Haddad, Sandra Coccuzzo Sampaio Vessoni, Simone Kashima, Svetoslav Nanev Slavov, Vincent Louis Viala |
| EPI_ISL_2345771 | CSII DR WASHINGTON LUIS M RODRIGUES DA SILVA PITANGUEIRAS | Instituto Butantan / Mendelics | Dimas Tadeu Covas, Antonio Jorge Martins, Claudia Renata dos Santos Barros, David Schlesinger, Debora Botequio Moretti, Elaine Cristina Marqueze, Elaine Vieira Santos, Evandra Strazza Rodrigues, Heidge Fukumasu, Jayme Augusto de Souza-Neto, José Salvatore Leister Patané, Luiz Alcantara, Luiz Lehmann Coutinho, Maria Carolina Elias, Maurício Lacerda Nogueira, Rafael dos Santos Bezerra, Raul Machado Neto, Rejane Maria Tommasini Grotto, Ricardo Haddad, Sandra Coccuzzo Sampaio Vessoni, Simone Kashima, Svetoslav Nanev Slavov, Vincent Louis Viala |
| EPI_ISL_2345859 | POLICLINICA COVID 19 ITAPETININGA | Instituto Butantan / Mendelics | Dimas Tadeu Covas, Antonio Jorge Martins, Claudia Renata dos Santos Barros, David Schlesinger, Debora Botequio Moretti, Elaine Cristina Marqueze, Elaine Vieira Santos, Evandra Strazza Rodrigues, Heidge Fukumasu, Jayme Augusto de Souza-Neto, José Salvatore Leister Patané, Luiz Alcantara, Luiz Lehmann Coutinho, Maria Carolina Elias, Maurício Lacerda Nogueira, Rafael dos Santos Bezerra, Raul Machado Neto, Rejane Maria Tommasini Grotto, Ricardo Haddad, Sandra Coccuzzo Sampaio Vessoni, Simone Kashima, Svetoslav Nanev Slavov, Vincent Louis Viala |
| EPI_ISL_2345865 | CENTRO DE REFERENCIA DO IDOSO DR HUMBERTO MENDES DE CARVALHO | Instituto Butantan / Mendelics | Dimas Tadeu Covas, Antonio Jorge Martins, Claudia Renata dos Santos Barros, David Schlesinger, Debora Botequio Moretti, Elaine Cristina Marqueze, Elaine Vieira Santos, Evandra Strazza Rodrigues, Heidge Fukumasu, Jayme Augusto de Souza-Neto, José Salvatore Leister Patané, Luiz Alcantara, Luiz Lehmann Coutinho, Maria Carolina Elias, Maurício Lacerda Nogueira, Rafael dos Santos Bezerra, Raul Machado Neto, Rejane Maria Tommasini Grotto, Ricardo Haddad, Sandra Coccuzzo Sampaio Vessoni, Simone Kashima, Svetoslav Nanev Slavov, Vincent Louis Viala |
| EPI_ISL_2345868 | CSII DR WASHINGTON LUIS M RODRIGUES DA SILVA PITANGUEIRAS | Instituto Butantan / Mendelics | Dimas Tadeu Covas, Antonio Jorge Martins, Claudia Renata dos Santos Barros, David Schlesinger, Debora Botequio Moretti, Elaine Cristina Marqueze, Elaine Vieira Santos, Evandra Strazza Rodrigues, Heidge Fukumasu, Jayme Augusto de Souza-Neto, José Salvatore Leister Patané, Luiz Alcantara, Luiz Lehmann Coutinho, Maria Carolina Elias, Maurício Lacerda Nogueira, Rafael dos Santos Bezerra, Raul Machado Neto, Rejane Maria Tommasini Grotto, Ricardo Haddad, Sandra Coccuzzo Sampaio Vessoni, Simone Kashima, Svetoslav Nanev Slavov, Vincent Louis Viala |
| EPI_ISL_2345943 | CENTRO INTEGRADO DE SAUDE | Instituto Butantan / ESALQ-Piracicaba | Dimas Tadeu Covas, Antonio Jorge Martins, Claudia Renata dos Santos Barros, David Schlesinger, Debora Botequio Moretti, Elaine Cristina Marqueze, Elaine Vieira Santos, Evandra Strazza Rodrigues, Heidge Fukumasu, Jayme Augusto de Souza-Neto, José Salvatore Leister Patané, Luiz Alcantara, Luiz Lehmann Coutinho, Maria Carolina Elias, Maurício Lacerda Nogueira, Rafael dos Santos Bezerra, Raul Machado Neto, Rejane Maria Tommasini Grotto, Ricardo Haddad, Sandra Coccuzzo Sampaio Vessoni, Simone Kashima, Svetoslav Nanev Slavov, Vincent Louis Viala |
| EPI_ISL_2346050, EPI_ISL_2346052, EPI_ISL_2346053, EPI_ISL_2346054, EPI_ISL_2346056 | SERRANA | Instituto Butantan / Mendelics | Dimas Tadeu Covas, Antonio Jorge Martins, Claudia Renata dos Santos Barros, David Schlesinger, Debora Botequio Moretti, Elaine Cristina Marqueze, Elaine Vieira Santos, Evandra Strazza Rodrigues, Heidge Fukumasu, Jayme Augusto de Souza-Neto, José Salvatore Leister Patané, Luiz Alcantara, Luiz Lehmann Coutinho, Maria Carolina Elias, Maurício Lacerda Nogueira, Rafael dos Santos Bezerra, Raul Machado Neto, Rejane Maria Tommasini Grotto, |

| | | | |
|---|---|---|---|
| | | | Ricardo Haddad, Sandra Coccuzzo Sampaio Vessoni, Simone Kashima, Svetoslav Nanev Slavov, Vincent Louis Viala |
| EPI_ISL_2382540 | Laboratório de Microbiologia Molecular - Universidade FEEVALE | Molecular Microbiology Laboratory | Alana Witt Hansen, Fágner Henrique Heldt, Fernando Rosado Spilki, Flávio Silveira, Juliana Schons Gularte, Juliane Deise Fleck, Mariana Soares da Silva, Meriane Demoliner, Matheus Nunes Weber, Paula Rodrigues de Almeida, Micheli Filippi. |
| EPI_ISL_2443551 | Labortorio Central de Saude Publica do Estado de Santa Catarina (LACEN/SC) | Laboratory of Respiratory Viruses and Measles, Oswaldo Cruz Institute, FIOCRUZ | Paola Resende, Luciana Appolinario, Fernando Motta, Anna Carolina Paixao, Ana Carolina Mendonca, Alice Sampaio Rocha, Taina Venas, Elisa Cavalcante Pereira, Renata Serrano Lopes, Darcita Buerger Rovaris, Sandra Bianchini Fernandes, Marilda Siqueira on behalf of the Fiocruz COVID-19 Genomic Surveillance Network |
| EPI_ISL_2444197, EPI_ISL_2444209 | SAO JOSE DO RIO PRETO | Instituto Butantan / FAMERP | Dimas Tadeu Covas, Antonio Jorge Martins, Claudia Renata dos Santos Barros, David Schlesinger, Debora Botequio Moretti, Elaine Cristina Marqueze, Elaine Vieira Santos, Evandra Strazza Rodrigues, Heidge Fukumasu, Jayme Augusto de Souza-Neto, José Salvatore Leister Patané, Luiz Alcantara, Luiz Lehmann Coutinho, Maria Carolina Elias, Maurício Lacerda Nogueira, Rafael dos Santos Bezerra, Raul Machado Neto, Rejane Maria Tommasini Grotto, Ricardo Haddad, Sandra Coccuzzo Sampaio Vessoni, Simone Kashima, Svetoslav Nanev Slavov, Vincent Louis Viala |
| EPI_ISL_2445221, EPI_ISL_2445222 | UNIDADE MISTA DE IGUAPE | Instituto Butantan | Dimas Tadeu Covas, Antonio Jorge Martins, Claudia Renata dos Santos Barros, David Schlesinger, Debora Botequio Moretti, Elaine Cristina Marqueze, Elaine Vieira Santos, Evandra Strazza Rodrigues, Heidge Fukumasu, Jayme Augusto de Souza-Neto, José Salvatore Leister Patané, Luiz Alcantara, Luiz Lehmann Coutinho, Maria Carolina Elias, Maurício Lacerda Nogueira, Rafael dos Santos Bezerra, Raul Machado Neto, Rejane Maria Tommasini Grotto, Ricardo Haddad, Sandra Coccuzzo Sampaio Vessoni, Simone Kashima, Svetoslav Nanev Slavov, Vincent Louis Viala |
| EPI_ISL_2445539 | CENTRO DE ESPECIALIDADES MARIAS DA GLORIA | Instituto Butantan | Dimas Tadeu Covas, Antonio Jorge Martins, Claudia Renata dos Santos Barros, David Schlesinger, Debora Botequio Moretti, Elaine Cristina Marqueze, Elaine Vieira Santos, Evandra Strazza Rodrigues, Heidge Fukumasu, Jayme Augusto de Souza-Neto, José Salvatore Leister Patané, Luiz Alcantara, Luiz Lehmann Coutinho, Maria Carolina Elias, Maurício Lacerda Nogueira, Rafael dos Santos Bezerra, Raul Machado Neto, Rejane Maria Tommasini Grotto, Ricardo Haddad, Sandra Coccuzzo Sampaio Vessoni, Simone Kashima, Svetoslav Nanev Slavov, Vincent Louis Viala |
| EPI_ISL_2445540 | HOSPITAL REGIONAL DE REGISTRO REGISTRO | Instituto Butantan | Dimas Tadeu Covas, Antonio Jorge Martins, Claudia Renata dos Santos Barros, David Schlesinger, Debora Botequio Moretti, Elaine Cristina Marqueze, Elaine Vieira Santos, Evandra Strazza Rodrigues, Heidge Fukumasu, Jayme Augusto de Souza-Neto, José Salvatore Leister Patané, Luiz Alcantara, Luiz Lehmann Coutinho, Maria Carolina Elias, Maurício Lacerda Nogueira, Rafael dos Santos Bezerra, Raul Machado Neto, Rejane Maria Tommasini Grotto, Ricardo Haddad, Sandra Coccuzzo Sampaio Vessoni, Simone Kashima, Svetoslav Nanev Slavov, Vincent Louis Viala |
| EPI_ISL_2466455 | HLAGYN - Laboratorio de Imunologia de Transplantes de Goias | HLAGYN - Laboratorio de Imunologia de Transplantes de Goias | Fernando Antonio Vinhal dos Santos, Erika Lopes Rocha Batista, Alessandro Leonardo Alvares Magalhaes, Frederico Rodrigues Vinhal, Sabrina Sara Moreira Duarte, Lucas Carlos Gomes Pereira, Daniel Ferreira de Sousa |
| EPI_ISL_2493285 | AFIP | Instituto Butantan | Dimas Tadeu Covas, Antonio Jorge Martins, Claudia Renata dos Santos Barros, David Schlesinger, Debora Botequio Moretti, Elaine Cristina Marqueze, Elaine Vieira Santos, Evandra Strazza Rodrigues, Heidge Fukumasu, Jayme Augusto de Souza-Neto, José Salvatore Leister Patané, Luiz Alcantara, Luiz Lehmann Coutinho, Maria Carolina Elias, Maurício Lacerda Nogueira, Rafael dos Santos Bezerra, Raul Machado Neto, Rejane Maria Tommasini Grotto, Ricardo Haddad, Sandra Coccuzzo Sampaio Vessoni, Simone Kashima, Svetoslav Nanev Slavov, Vincent Louis Viala |
| EPI_ISL_2493473 | UNIDADE DE PRONTO ATENDIMENTO | Instituto Butantan | Dimas Tadeu Covas, Antonio Jorge Martins, Claudia Renata dos Santos Barros, David Schlesinger, Debora Botequio Moretti, Elaine Cristina Marqueze, Elaine Vieira Santos, Evandra Strazza Rodrigues, Heidge Fukumasu, Jayme Augusto de Souza-Neto, José Salvatore Leister Patané, Luiz Alcantara, Luiz Lehmann Coutinho, Maria Carolina Elias, Maurício Lacerda Nogueira, Rafael dos Santos Bezerra, Raul Machado Neto, Rejane Maria Tommasini Grotto, Ricardo Haddad, Sandra Coccuzzo Sampaio Vessoni, Simone Kashima, Svetoslav Nanev Slavov, Vincent Louis Viala |
| EPI_ISL_2493539 | CENTRO DE SAUDE III BORBOREMA | Instituto Butantan | Dimas Tadeu Covas, Antonio Jorge Martins, Claudia Renata dos Santos Barros, David Schlesinger, Debora Botequio Moretti, Elaine Cristina Marqueze, Elaine Vieira Santos, Evandra Strazza Rodrigues, Heidge Fukumasu, Jayme Augusto de Souza-Neto, José Salvatore Leister Patané, Luiz Alcantara, Luiz Lehmann Coutinho, Maria Carolina Elias, Maurício Lacerda Nogueira, Rafael dos Santos Bezerra, Raul Machado Neto, Rejane Maria Tommasini Grotto, Ricardo Haddad, Sandra Coccuzzo Sampaio Vessoni, Simone Kashima, Svetoslav Nanev Slavov, Vincent Louis Viala |
| EPI_ISL_2493579, EPI_ISL_2493581 | SECRETARIA MUNICIPAL DE SAUDE | Instituto Butantan | Dimas Tadeu Covas, Antonio Jorge Martins, Claudia Renata dos Santos Barros, David Schlesinger, Debora Botequio Moretti, Elaine Cristina Marqueze, Elaine Vieira Santos, Evandra Strazza Rodrigues, Heidge Fukumasu, Jayme Augusto de Souza-Neto, José Salvatore Leister Patané, Luiz Alcantara, Luiz Lehmann Coutinho, Maria Carolina Elias, Maurício Lacerda Nogueira, Rafael dos Santos Bezerra, Raul Machado Neto, Rejane Maria Tommasini Grotto, Ricardo Haddad, Sandra Coccuzzo Sampaio Vessoni, Simone Kashima, Svetoslav Nanev Slavov, Vincent Louis Viala |
| EPI_ISL_2493592 | HOSPITAL MUNICIPAL DE ILHABELA GOV MARIO COVAS JR | Instituto Butantan | Dimas Tadeu Covas, Antonio Jorge Martins, Claudia Renata dos Santos Barros, David Schlesinger, Debora Botequio Moretti, Elaine Cristina Marqueze, Elaine Vieira Santos, Evandra Strazza Rodrigues, Heidge Fukumasu, Jayme Augusto de Souza-Neto, José Salvatore Leister Patané, Luiz Alcantara, Luiz Lehmann Coutinho, Maria Carolina Elias, Maurício Lacerda Nogueira, Rafael dos Santos Bezerra, Raul Machado Neto, Rejane Maria Tommasini Grotto, Ricardo Haddad, Sandra Coccuzzo Sampaio Vessoni, Simone Kashima, Svetoslav Nanev Slavov, Vincent Louis Viala |
| EPI_ISL_2493603 | SECRETARIA MUNICIPAL DA SAUDE DE GUARIBA | Instituto Butantan | Dimas Tadeu Covas, Antonio Jorge Martins, Claudia Renata dos Santos Barros, David Schlesinger, Debora Botequio Moretti, Elaine Cristina Marqueze, Elaine Vieira Santos, Evandra Strazza Rodrigues, Heidge Fukumasu, Jayme Augusto de Souza-Neto, José Salvatore Leister Patané, Luiz Alcantara, Luiz Lehmann Coutinho, Maria Carolina Elias, Maurício Lacerda Nogueira, Rafael dos Santos Bezerra, Raul Machado Neto, Rejane Maria Tommasini Grotto, Ricardo Haddad, Sandra Coccuzzo Sampaio Vessoni, Simone Kashima, Svetoslav Nanev Slavov, Vincent Louis Viala |
| EPI_ISL_2493651 | HOSPITAL MUNICIPAL DE ILHABELA GOV MARIO COVAS JR | Instituto Butantan | Dimas Tadeu Covas, Antonio Jorge Martins, Claudia Renata dos Santos Barros, David Schlesinger, Debora Botequio Moretti, Elaine Cristina Marqueze, Elaine Vieira Santos, Evandra Strazza Rodrigues, Heidge Fukumasu, Jayme Augusto de Souza-Neto, José Salvatore Leister Patané, Luiz Alcantara, Luiz Lehmann Coutinho, Maria Carolina Elias, Maurício Lacerda Nogueira, Rafael dos Santos Bezerra, Raul Machado Neto, Rejane Maria Tommasini Grotto, Ricardo Haddad, Sandra Coccuzzo Sampaio Vessoni, Simone Kashima, Svetoslav Nanev Slavov, Vincent Louis Viala |
| EPI_ISL_2493667 | POLICLINICA COVID 19 ITAPETININGA | Instituto Butantan | Dimas Tadeu Covas, Antonio Jorge Martins, Claudia Renata dos Santos Barros, David Schlesinger, Debora Botequio Moretti, Elaine Cristina Marqueze, Elaine Vieira Santos, Evandra Strazza Rodrigues, Heidge Fukumasu, Jayme Augusto de Souza-Neto, José Salvatore Leister Patané, Luiz Alcantara, Luiz Lehmann Coutinho, Maria Carolina Elias, Maurício Lacerda Nogueira, Rafael dos Santos Bezerra, Raul Machado Neto, Rejane Maria Tommasini Grotto, Ricardo Haddad, Sandra Coccuzzo Sampaio Vessoni, Simone Kashima, Svetoslav Nanev Slavov, Vincent Louis Viala |
| EPI_ISL_2493677 | CENTRO DE SAUDE III TABATINGA | Instituto Butantan | Dimas Tadeu Covas, Antonio Jorge Martins, Claudia Renata dos Santos Barros, David Schlesinger, Debora Botequio Moretti, Elaine Cristina Marqueze, Elaine Vieira Santos, Evandra Strazza Rodrigues, Heidge Fukumasu, Jayme Augusto de Souza-Neto, José Salvatore Leister Patané, Luiz Alcantara, Luiz Lehmann Coutinho, Maria Carolina Elias, Maurício Lacerda Nogueira, Rafael dos Santos Bezerra, Raul Machado Neto, Rejane Maria Tommasini Grotto, Ricardo Haddad, Sandra Coccuzzo Sampaio Vessoni, Simone Kashima, Svetoslav Nanev Slavov, Vincent Louis Viala |
| EPI_ISL_2493698 | SECRETARIA MUNICIPAL DE SAUDE SOROCABA | Instituto Butantan | Dimas Tadeu Covas, Antonio Jorge Martins, Claudia Renata dos Santos Barros, David Schlesinger, Debora Botequio Moretti, Elaine Cristina Marqueze, Elaine Vieira Santos, Evandra Strazza Rodrigues, Heidge Fukumasu, Jayme Augusto de Souza-Neto, José Salvatore Leister Patané, Luiz Alcantara, Luiz Lehmann Coutinho, Maria Carolina Elias, Maurício Lacerda Nogueira, Rafael dos Santos Bezerra, Raul Machado Neto, Rejane Maria Tommasini Grotto, Ricardo Haddad, Sandra Coccuzzo Sampaio Vessoni, Simone Kashima, Svetoslav Nanev Slavov, Vincent Louis Viala |
| EPI_ISL_2493831 | USAFA SAO JORGE | Instituto Butantan | Dimas Tadeu Covas, Antonio Jorge Martins, Claudia Renata dos Santos Barros, David Schlesinger, Debora Botequio Moretti, Elaine Cristina Marqueze, Elaine Vieira Santos, Evandra Strazza Rodrigues, Heidge Fukumasu, Jayme Augusto de Souza-Neto, José Salvatore Leister Patané, Luiz Alcantara, Luiz Lehmann Coutinho, Maria Carolina Elias, Maurício Lacerda Nogueira, Rafael dos Santos Bezerra, Raul Machado Neto, Rejane Maria Tommasini Grotto, Ricardo Haddad, Sandra Coccuzzo Sampaio Vessoni, Simone Kashima, Svetoslav Nanev Slavov, Vincent Louis Viala |
| EPI_ISL_2493858 | UPA SADAKO SEDOGUTI | Instituto Butantan | Dimas Tadeu Covas, Antonio Jorge Martins, Claudia Renata dos Santos Barros, David Schlesinger, Debora Botequio Moretti, Elaine Cristina Marqueze, Elaine Vieira Santos, Evandra Strazza Rodrigues, Heidge Fukumasu, Jayme Augusto de Souza-Neto, José Salvatore Leister Patané, Luiz Alcantara, Luiz Lehmann Coutinho, Maria Carolina Elias, Maurício Lacerda Nogueira, Rafael dos Santos Bezerra, Raul Machado Neto, Rejane Maria Tommasini Grotto, Ricardo Haddad, Sandra Coccuzzo Sampaio Vessoni, Simone Kashima, Svetoslav Nanev Slavov, Vincent Louis Viala |
| EPI_ISL_2494112 | VIGILANCIA EM SAUDE SAE VIGILANCIA EPIDEMIOLOGICA | Instituto Butantan | Dimas Tadeu Covas, Antonio Jorge Martins, Claudia Renata dos Santos Barros, David Schlesinger, Debora Botequio Moretti, Elaine Cristina Marqueze, Elaine Vieira Santos, Evandra Strazza Rodrigues, Heidge Fukumasu, Jayme Augusto de Souza-Neto, José Salvatore Leister Patané, Luiz Alcantara, Luiz Lehmann Coutinho, Maria Carolina Elias, Maurício Lacerda Nogueira, Rafael dos Santos Bezerra, Raul Machado Neto, Rejane Maria Tommasini Grotto, Ricardo Haddad, Sandra Coccuzzo Sampaio Vessoni, Simone Kashima, Svetoslav Nanev Slavov, Vincent Louis Viala |
| EPI_ISL_2494127 | CENTRO DE REFERENCIA DO IDOSO DR HUMBERTO | Instituto Butantan | Dimas Tadeu Covas, Antonio Jorge Martins, Claudia Renata dos Santos Barros, David Schlesinger, Debora Botequio Moretti, Elaine Cristina Marqueze, |

| | | | |
|---|---|---|---|
| | MENDES DE CARVALHO | | Elaine Vieira Santos, Evandra Strazza Rodrigues, Heidge Fukumasu, Jayme Augusto de Souza-Neto, José Salvatore Leister Patané, Luiz Alcantara, Luiz Lehmann Coutinho, Maria Carolina Elias, Maurício Lacerda Nogueira, Rafael dos Santos Bezerra, Raul Machado Neto, Rejane Maria Tommasini Grotto, Ricardo Haddad, Sandra Coccuzzo Sampaio Vessoni, Simone Kashima, Svetoslav Nanev Slavov, Vincent Louis Viala |
| EPI_ISL_2494321 | CSII DR WASHINGTON LUIS M RODRIGUES DA SILVA PITANGUEIRAS | Instituto Butantan | Dimas Tadeu Covas, Antonio Jorge Martins, Claudia Renata dos Santos Barros, David Schlesinger, Debora Botequio Moretti, Elaine Cristina Marqueze, Elaine Vieira Santos, Evandra Strazza Rodrigues, Heidge Fukumasu, Jayme Augusto de Souza-Neto, José Salvatore Leister Patané, Luiz Alcantara, Luiz Lehmann Coutinho, Maria Carolina Elias, Maurício Lacerda Nogueira, Rafael dos Santos Bezerra, Raul Machado Neto, Rejane Maria Tommasini Grotto, Ricardo Haddad, Sandra Coccuzzo Sampaio Vessoni, Simone Kashima, Svetoslav Nanev Slavov, Vincent Louis Viala |
| EPI_ISL_2497464 | HLAGYN - Laboratorio de Imunologia de Transplantes de Goias | HLAGYN - Laboratorio de Imunologia de Transplantes de Goias | Fernando Antonio Vinhal dos Santos, Erika Lopes Rocha Batista, Alessandro Leonardo Alvares Magalhaes, Frederico Rodrigues Vinhal, Sabrina Sara Moreira Duarte, Lucas Carlos Gomes Pereira, Daniel Ferreira de Sousa |
| EPI_ISL_2544855, EPI_ISL_2544859 | Laboratorio de Pesquisa em Virologia, FAMERP, SJRP | Laboratorio de Pesquisa em Virologia, FAMERP, SJRP | Cecília Artico Banho; Lívia Sacchetto; Guilherme Campos; Fábio Sossai Possebon; Leila Sabrina Ullmann; Cíntia Bittar; Helena Lage Ferreira; Jorge A. Petroli Marchesi; Maisa C. Pereira Parra; Marília Moraes; Paula Rahal; Paulo Inacio da Costa; João Pessoa Araújo Jr.; Maurício L. Nogueira. |
| EPI_ISL_2614117 | Laboratory of Respiratory Viruses and Measles, Oswaldo Cruz Institute, FIOCRUZ | Laboratory of Respiratory Viruses and Measles, Oswaldo Cruz Institute, FIOCRUZ | Paola Resende, Luciana Appolinario, Fernando Motta, Anna Carolina Paixao, Ana Carolina Mendonca, Alice Sampaio Rocha, Taina Venas, Elisa Cavalcante Pereira, Renata Serrano Lopes, Marilda Siqueira on behalf of the Fiocruz COVID-19 Genomic Surveillance Network |
| EPI_ISL_2614590 | Hospital Municipal Dr. Guido Guida | Instituto Adolfo Lutz, Interdiciplinary Procedures Center, Strategic Laboratory | Claudio Tavares Sacchi, Claudia Regina Gonçalves, Erica Valessa Ramos Gomes, Karoline Rodrigues Campos, Caio Vinicius Dias Lopes, Leonardo Jose Tadeu de Araujo |
| EPI_ISL_2661768 | Laboratorio Central de Saude Publica do Estado do Rio Grande do Sul (LACEN-RS) | Laboratory of Respiratory Viruses and Measles, Oswaldo Cruz Institute, FIOCRUZ | Paola Resende, Luciana Appolinario, Fernando Motta, Anna Carolina Paixao, Ana Carolina Mendonca, Alice Sampaio Rocha, Taina Venas, Elisa Cavalcante Pereira, Renata Serrano Lopes, Tatiana Schaffer Gregianini, Richard Salvato, Marilda Siqueira on behalf of the Fiocruz COVID-19 Genomic Surveillance Network |
| EPI_ISL_2680903 | HLAGYN - Laboratorio de Imunologia de Transplantes de Goias | HLAGYN - Laboratorio de Imunologia de Transplantes de Goias | Fernando Antonio Vinhal dos Santos, Erika Lopes Rocha Batista, Alessandro Leonardo Alvares Magalhaes, Frederico Rodrigues Vinhal, Sabrina Sara Moreira Duarte, Lucas Carlos Gomes Pereira, Daniel Ferreira de Sousa, Danielle de Paiva Rezende |
| EPI_ISL_2691097 | Instituto Adolfo Lutz - Regional de Marilia | Instituto Adolfo Lutz, Interdiciplinary Procedures Center, Strategic Laboratory | Claudio Tavares Sacchi, Claudia Regina Gonçalves, Erica Valessa Ramos Gomes, Karoline Rodrigues Campos, Caio Vinicius Dias Lopes, Leonardo Jose Tadeu de Araujo |
| EPI_ISL_2691129 | Instituto Adolfo Lutz Central | Instituto Adolfo Lutz, Interdiciplinary Procedures Center, Strategic Laboratory | Claudio Tavares Sacchi, Claudia Regina Gonçalves, Erica Valessa Ramos Gomes, Karoline Rodrigues Campos, Caio Vinicius Dias Lopes, Leonardo Jose Tadeu de Araujo |
| EPI_ISL_2691168 | Instituo Adolfo Lutz Central | Instituto Adolfo Lutz, Interdiciplinary Procedures Center, Strategic Laboratory | Claudio Tavares Sacchi, Claudia Regina Gonçalves, Erica Valessa Ramos Gomes, Karoline Rodrigues Campos, Caio Vinicius Dias Lopes, Leonardo Jose Tadeu de Araujo |
| EPI_ISL_2691578 | Unidade de apoio ao diagnostico da COVID - UNADIG | Bioinformatics Laboratory / LNCC | Luiz G P de Almeida, Alessandra P Lamarca, Ronaldo da Silva F Jr, Liliane Cavalcante, Alexandra L Gerber, Ana Paula de C Guimaraes, Douglas Terra Machado, Cassia Alves, Diana Mariani, Cintia Policarpo, Gleidson da Silva de Oliveira, Mario Sergio Ribeiro, Silvia Carvalho, Flavio Dias da Silva, Marcio Henrique de Oliveira Garcia, Leandro Magalhaes de Souza, Cristiane Gomes da Silva, Caio Luiz Pereira Ribeiro, Andrea Cony Cavalcanti, Claudia Maria Braga de Mello, Amilcar Tanuri, Ana Tereza R Vasconcelos |
| EPI_ISL_717808 | Laboratorio de Virologia Molecular / UFRJ | Bioinformatics Laboratory / LNCC | Carolina M Voloch, Ronaldo da Silva F Jr, Luiz G P de Almeida, Cynthia C Cardoso, Otavio Bustrolini, Alexandra L Gerber, Ana Paula de C Guimarães, Diana Mariani, Andréa Cony Cavalcanti, Claudia dos Santos Rodrigues, Terezinha M P P Castiñeira, Amilcar Tanuri, Ana Tereza R de Vasconcelos |
| EPI_ISL_735403 | Instituto Adolfo Lutz - Regional de Rio Claro | Instituto Adolfo Lutz, Interdiciplinary Procedures Center, Strategic Laboratory | Claudio Tavares Sacchi, Claudia Regina Gonçalves, Erica Valessa Ramos Gomes, Karoline Rodrigues Campos |
| EPI_ISL_940628 | Unidade Mista de Iguape | Instituto Adolfo Lutz, Interdiciplinary Procedures Center, Strategic Laboratory | Claudio Tavares Sacchi, Claudia Regina Gonçalves, Erica Valessa Ramos Gomes, Karoline Rodrigues Campos |

We gratefully acknowledge the following Authors from the Originating laboratories responsible for obtaining the specimens, as well as the Submitting laboratories where the genome data were generated and shared via GISAID, on which this research is based.

All Submitters of data may be contacted directly via www.gisaid.org

Authors are sorted alphabetically.

| Accession ID | Originating Laboratory | Submitting Laboratory | Authors |
|---|---|---|---|
| EPI_ISL_1795385, EPI_ISL_1795386 | UNIDADE DE VIGILANCIA EM SAUDE | Instituto Butantan / ESALQ-Piracicaba | Instituto Butantan: Alexander Roberto Precioso, Dimas Tadeu Covas, Sandra Coccuzzo Sampaio, Maria Carolina Elias, José Salvatore Leister Patané, Vincent Louis Viala, Antonio Jorge Martins, Ricardo Haddad, Claudia Renata dos Santos Barros, Elaine Cristina Marqueze, Raul Machado Neto, Debora Botequio Moretti. Centro de Genômica Funcional da ESALQ: Luiz Lehmann Coutinho, Ricardo Augusto Brassaloti, Raquel de Lello Rocha Campos Cassano. NGS Soluções Genômicas: Pilar Drummond Sampaio Corrêa Mariani. FZEA-USP Pirassununga: Mirele Daiana Poleti, Jessika Cristina Chagas Lesbon, Elisangela Chicaroni Mattos, Heidge Fukumasu. USP-Botucatu: Rejane Maria Tommasini Grotto, Jayme A. Souza-Neto, Guilherme Targino Valente, Patricia Akemi Assato, Felipe Allan da Silva da Costa, Bianca Cechetto Carlos. Mendelics: Bibiana Santos, João Paulo Kitajima, Erika Freitas, David Schlesinger. Hemocentro Ribeirão Preto: Simone Kashima, Evandra Strazza Rodrigues, Svetoslav Nanev Slavov, Elaine Vieira dos Santos, Rafael dos Santos Bezerra, Luiz Carlos Junior de Alcantara, Marta Giovanetti, Vagner Fonseca, Flavia Aburjaile, Rodrigo Tocantins Calado. |
| EPI_ISL_2086590, EPI_ISL_2086764 | LABCOVID_HCPA | LABRESIS_HCPA | Wink PL, Martins AF, Volpato F, Monteiro F, Zavascki AP, Barth AL |
| EPI_ISL_2102518, EPI_ISL_2102519, EPI_ISL_2102520 | HLAGYN - Laboratorio de Imunologia de Transplantes de Goias | HLAGYN - Laboratorio de Imunologia de Transplantes de Goias | Fernando Antonio Vinhal dos Santos, Erika Lopes Rocha Batista, Alessandro Leonardo Alvares Magalhaes, Frederico Rodrigues Vinhal, Sabrina Sara Moreira Duarte, Danielle de Paiva Rezende, Lucas Carlos Gomes Pereira, Paola Cristina Resende Silva |
| EPI_ISL_2245112 | Laboratório Central de Saúde Pública de Roraima | Coordenação Geral de Laboratórios de Saúde Pública (CGLAB/DAEVS/SVS/MS) | Vagner Fonseca, et al. |
| EPI_ISL_2345272, EPI_ISL_2345275, EPI_ISL_2345277, EPI_ISL_2345279, EPI_ISL_2345323, EPI_ISL_2345336 | UNIDADE DE VIGILANCIA EM SAUDE | Instituto Butantan / ESALQ-Piracicaba | Dimas Tadeu Covas, Antonio Jorge Martins, Claudia Renata dos Santos Barros, David Schlesinger, Debora Botequio Moretti, Elaine Cristina Marqueze, Elaine Vieira Santos, Evandra Strazza Rodrigues, Heidge Fukumasu, Jayme Augusto de Souza-Neto, José Salvatore Leister Patané, Luiz Alcantara, Luiz Lehmann Coutinho, Maria Carolina Elias, Maurício Lacerda Nogueira, Rafael dos Santos Bezerra, Raul Machado Neto, Rejane Maria Tommasini Grotto, Ricardo Haddad, Sandra Coccuzzo Sampaio Vessoni, Simone Kashima, Svetoslav Nanev Slavov, Vincent Louis Viala |
| EPI_ISL_2345696, EPI_ISL_2345748 | CENTRO DE SAUDE III SANTA ERNESTINA | Instituto Butantan / Mendelics | Dimas Tadeu Covas, Antonio Jorge Martins, Claudia Renata dos Santos Barros, David Schlesinger, Debora Botequio Moretti, Elaine Cristina Marqueze, Elaine Vieira Santos, Evandra Strazza Rodrigues, Heidge Fukumasu, Jayme Augusto de Souza-Neto, José Salvatore Leister Patané, Luiz Alcantara, Luiz Lehmann Coutinho, Maria Carolina Elias, Maurício Lacerda Nogueira, Rafael dos Santos Bezerra, Raul Machado Neto, Rejane Maria Tommasini Grotto, Ricardo Haddad, Sandra Coccuzzo Sampaio Vessoni, Simone Kashima, Svetoslav Nanev Slavov, Vincent Louis Viala |
| EPI_ISL_2385704, EPI_ISL_2385705, EPI_ISL_2385770 | Unidade de apoio ao diagnóstico da COVID - UNADIG | Bioinformatics Laboratory / LNCC | Luiz G P de Almeida, Alessandra P Lamarca, Ronaldo da Silva F Jr, Liliane Cavalcante, Alexandra L Gerber, Ana Paula de C Guimaraes, Douglas Terra Machado, Cassia Alves, Diana Mariani, Cintia Policarpo, Gleidson da Silva de Oliveira, Mario Sergio Ribeiro, Silvia Carvalho, Flavio Dias da Silva, Marcio Henrique de Oliveira Garcia, Leandro Magalhaes de Souza, Cristiane Gomes da Silva, Caio Luiz Pereira Ribeiro, Andrea Cony Cavalcanti, Claudia Maria Braga de Mello, Amilcar Tanuri, Ana Tereza R Vasconcelos |
| EPI_ISL_2445233, EPI_ISL_2445238 | HOSPITAL MUNICIPAL REYNALDO GUERRA CAJATI | Instituto Butantan | Dimas Tadeu Covas, Antonio Jorge Martins, Claudia Renata dos Santos Barros, David Schlesinger, Debora Botequio Moretti, Elaine Cristina Marqueze, Elaine Vieira Santos, Evandra Strazza Rodrigues, Heidge Fukumasu, Jayme Augusto de Souza-Neto, José Salvatore Leister Patané, Luiz Alcantara, Luiz Lehmann Coutinho, Maria Carolina Elias, Maurício Lacerda Nogueira, Rafael dos Santos Bezerra, Raul Machado Neto, Rejane Maria Tommasini Grotto, Ricardo Haddad, Sandra Coccuzzo Sampaio Vessoni, Simone Kashima, Svetoslav Nanev Slavov, Vincent Louis Viala |
| EPI_ISL_2445497 | LABORATORIO DE FRANCA | Instituto Butantan | Dimas Tadeu Covas, Antonio Jorge Martins, Claudia Renata dos Santos Barros, David Schlesinger, Debora Botequio Moretti, Elaine Cristina Marqueze, Elaine Vieira Santos, Evandra Strazza Rodrigues, Heidge Fukumasu, Jayme Augusto de Souza-Neto, José Salvatore Leister Patané, Luiz Alcantara, Luiz Lehmann Coutinho, Maria Carolina Elias, Maurício Lacerda Nogueira, Rafael dos Santos Bezerra, Raul Machado Neto, Rejane Maria Tommasini Grotto, Ricardo Haddad, Sandra Coccuzzo Sampaio Vessoni, Simone Kashima, Svetoslav Nanev Slavov, Vincent Louis Viala |
| EPI_ISL_2473819 | UNIDADE DE PRONTO ATENDIMENTO UPA | Instituto Butantan | Dimas Tadeu Covas, Antonio Jorge Martins, Claudia Renata dos Santos Barros, David Schlesinger, Debora Botequio Moretti, Elaine Cristina Marqueze, Elaine Vieira Santos, Evandra Strazza Rodrigues, Heidge Fukumasu, Jayme Augusto de Souza-Neto, José Salvatore Leister Patané, Luiz Alcantara, Luiz Lehmann Coutinho, Maria Carolina Elias, Maurício Lacerda Nogueira, Rafael dos Santos Bezerra, Raul Machado Neto, Rejane Maria Tommasini Grotto, Ricardo Haddad, Sandra Coccuzzo Sampaio Vessoni, Simone Kashima, Svetoslav Nanev Slavov, Vincent Louis Viala |
| EPI_ISL_2493720 | INSIDE DIAGNÓSTICOS | Instituto Butantan | Dimas Tadeu Covas, Antonio Jorge Martins, Claudia Renata dos Santos Barros, David Schlesinger, Debora Botequio Moretti, Elaine Cristina Marqueze, Elaine Vieira Santos, Evandra Strazza Rodrigues, Heidge Fukumasu, Jayme Augusto de Souza-Neto, José Salvatore Leister Patané, Luiz Alcantara, Luiz Lehmann Coutinho, Maria Carolina Elias, Maurício Lacerda Nogueira, Rafael dos Santos Bezerra, Raul Machado Neto, Rejane Maria Tommasini Grotto, Ricardo Haddad, Sandra Coccuzzo Sampaio Vessoni, Simone Kashima, Svetoslav Nanev Slavov, Vincent Louis Viala |
| EPI_ISL_2493774 | BIOFAST | Instituto Butantan | Dimas Tadeu Covas, Antonio Jorge Martins, Claudia Renata dos Santos Barros, David Schlesinger, Debora Botequio Moretti, Elaine Cristina Marqueze, Elaine Vieira Santos, Evandra Strazza Rodrigues, Heidge Fukumasu, Jayme Augusto de Souza-Neto, José Salvatore Leister Patané, Luiz Alcantara, Luiz Lehmann Coutinho, Maria Carolina Elias, Maurício Lacerda Nogueira, Rafael dos Santos Bezerra, Raul Machado Neto, Rejane Maria Tommasini Grotto, Ricardo Haddad, Sandra Coccuzzo Sampaio Vessoni, Simone Kashima, Svetoslav Nanev Slavov, Vincent Louis Viala |
| EPI_ISL_2614328 | Laboratory of Respiratory Viruses and Measles, Oswaldo Cruz Institute, FIOCRUZ | Laboratory of Respiratory Viruses and Measles, Oswaldo Cruz Institute, FIOCRUZ | Paola Resende, Luciana Appolinario, Fernando Motta, Anna Carolina Paixao, Ana Carolina Mendonca, Alice Sampaio Rocha, Taina Venas, Elisa Cavalcante Pereira, Renata Serrano Lopes, Marilda Siqueira on behalf of the Fiocruz COVID-19 Genomic Surveillance Network |
| EPI_ISL_2645663 | Laboratorio Central de Saude Publica do Estado de Alagoas (LACEN/AL) | Laboratory of Respiratory Viruses and Measles, Oswaldo Cruz Institute, FIOCRUZ | Paola Resende, Luciana Appolinario, Fernando Motta, Anna Carolina Paixao, Ana Carolina Mendonca, Alice Sampaio Rocha, Taina Venas, Elisa Cavalcante Pereira, Renata Serrano Lopes, Anderson Brandao Leite, Marilda Siqueira on behalf of the Fiocruz COVID-19 Genomic Surveillance Network |
| EPI_ISL_2660533 | Laboratorio Central de Saude Publica do Estado de Minas Gerais (LACEN/MG) | Laboratory of Respiratory Viruses and Measles, Oswaldo Cruz Institute, FIOCRUZ | Paola Resende, Luciana Appolinario, Fernando Motta, Anna Carolina Paixao, Ana Carolina Mendonca, Alice Sampaio Rocha, Taina Venas, Elisa Cavalcante Pereira, Renata Serrano Lopes, Andre Felipe Leal Bernardes, Marilda Siqueira on behalf of the Fiocruz COVID-19 Genomic Surveillance Network |
| EPI_ISL_2691132, EPI_ISL_2691135 | Instituto Adolfo Lutz Central | Instituto Adolfo Lutz, Interdiciplinary Procedures Center, Strategic Laboratory | Claudio Tavares Sacchi, Claudia Regina Gonçalves, Erica Valessa Ramos Gomes, Karoline Rodrigues Campos, Caio Vinicius Dias Lopes, Leonardo Jose Tadeu de Araujo |
| EPI_ISL_2691664 | Unidade de apoio ao diagnostico da COVID - UNADIG | Bioinformatics Laboratory / LNCC | Luiz G P de Almeida, Alessandra P Lamarca, Ronaldo da Silva F Jr, Liliane Cavalcante, Alexandra L Gerber, Ana Paula de C Guimaraes, Douglas Terra Machado, Cassia Alves, Diana Mariani, Cintia Policarpo, Gleidson da Silva de Oliveira, Mario Sergio Ribeiro, Silvia Carvalho, Flavio Dias da Silva, Marcio Henrique de Oliveira Garcia, Leandro Magalhaes de Souza, Cristiane Gomes da Silva, Caio Luiz Pereira Ribeiro, Andrea Cony Cavalcanti, Claudia Maria Braga de Mello, Amilcar Tanuri, Ana Tereza R Vasconcelos |

We gratefully acknowledge the following Authors from the Originating laboratories responsible for obtaining the specimens, as well as the Submitting laboratories where the genome data were generated and shared via GISAID, on which this research is based.

All Submitters of data may be contacted directly via www.gisaid.org

Authors are sorted alphabetically.

| Accession ID | Originating Laboratory | Submitting Laboratory | Authors |
|---|---|---|---|
| EPI_ISL_1133124 | LABCOVID_HCPA | LABRESIS_HCPA | Martins AF, Wink PL, Volpato F, Rosset C, de Paris F, Monteiro F, Barth AL |
| EPI_ISL_1445142 | CS II EGIDIO BRUNHARA MORRO AGUDO | Instituto Butantan / Mendelics | Dimas Tadeu Covas, Sandra Coccuzzo Sampaio, Maria Carolina Elias, José Salvatore Leister Patané, Vincent Louis Viala, Antonio Jorge Martins, Ricardo Haddad, Claudia Renata dos Santos Barros, Elaine Cristina Marqueze, Raul Machado Neto, Debora Botequio Moretti, Bibiana Santos, João Paulo Kitajima, Erika Freitas, David Schlesinger, Simone Kashima, Evandra Strazza Rodrigues, Svetoslav Nanev Slavov, Elaine Vieira dos Santos, Rafael dos Santos Bezerra, Luiz Carlos Junior de Alcantara, Marta Giovanetti, Vagner Fonseca, Flavia Aburjaile, Rodrigo Tocantins Calado. |
| EPI_ISL_1583681 | Central Public Health Laboratory - LACEN -Bahia, Salvador, Brazil | Central Public Health Laboratory - LACEN -Bahia, Salvador, Brazil | Stephane Tosta, Luciana Oliveira, Vanessa Nardy,Patrícia Cajado,Marcela Gómez, Breno Dominguez, Jaqueline Gomes, Vagner Fonseca,Marta Giovanetti,Luiz Alcantara, Felicidade Pereira, Arabela Leal |
| EPI_ISL_1664164, EPI_ISL_1664165, EPI_ISL_1664166, EPI_ISL_1664167, EPI_ISL_1664168, EPI_ISL_1664169, EPI_ISL_1664170, EPI_ISL_1664171, EPI_ISL_1664172, EPI_ISL_1664173 | Laboratorio Central Noel Nutels | Bioinformatics Laboratory / LNCC | Luiz G P de Almeida, Alessandra P Lamarca, Ronaldo da Silva F Jr, Liliane Cavalcante, Alexandra L Gerber, Ana Paula de C Guimarães, Douglas Terra Machado, Cassia Alves, Diana Mariani, Thais Felix Cruz, Mario Sergio Ribeiro, Silvia Carvalho, Flávio Dias da Silva, Marcio Henrique de Oliveira Garcia, Leandro Magalhães de Souza, Cristiane Gomes da Silva, Caio Luiz Pereira Ribeiro, Andréa Cony Cavalcanti, Claudia Maria Braga de Mello, Amilcar Tanuri, Ana Tereza R Vasconcelos |
| EPI_ISL_1715149, EPI_ISL_1715159 | Instituto Adolfo Lutz Central | Instituto Adolfo Lutz, Interdiciplinary Procedures Center, Strategic Laboratory | Claudio Tavares Sacchi, Claudia Regina Gonçalves, Erica Valessa Ramos Gomes, Karoline Rodrigues Campos, Caio Vinicius Dias Lopes, Leonardo Jose Tadeu de Araujo, Katia Correa de Oliveira Santos |
| EPI_ISL_1752639 | UPA Dr Luis Atilio Losi Viana Ribeirao Preto | Instituto Adolfo Lutz, Interdiciplinary Procedures Center, Strategic Laboratory | Claudio Tavares Sacchi, Claudia Regina Gonçalves, Erica Valessa Ramos Gomes, Karoline Rodrigues Campos, Caio Vinicius Dias Lopes, Leonardo Jose Tadeu de Araujo, Katia Correa de Oliveira Santos |
| EPI_ISL_1795103 | HOSPITAL DOS FORNCEDORES | Instituto Butantan / ESALQ-Piracicaba | Instituto Butantan: Alexander Roberto Precioso, Dimas Tadeu Covas, Sandra Coccuzzo Sampaio, Maria Carolina Elias, José Salvatore Leister Patané, Vincent Louis Viala, Antonio Jorge Martins, Ricardo Haddad, Claudia Renata dos Santos Barros, Elaine Cristina Marqueze, Raul Machado Neto, Debora Botequio Moretti. Centro de Genômica Funcional da ESALQ: Luiz Lehmann Coutinho, Ricardo Augusto Brassaloti, Raquel de Lello Rocha Campos Cassano. NGS Soluções Genômicas: Pilar Drummond Sampaio Corrêa Mariani. FZEA-USP Pirassununga: Mirele Daiana Poleti, Jessika Cristina Chagas Lesbon, Elisangela Chicaroni Mattos, Heidge Fukumasu. USP-Botucatu: Rejane Maria Tommasini Grotto, Jayme A. Souza-Neto, Guilherme Targino Valente, Patricia Akemi Assato, Felipe Allan da Silva da Costa, Bianca Cechetto Carlos. Mendelics: Bibiana Santos, João Paulo Kitajima, Erika Freitas, David Schlesinger. Hemocentro Ribeirão Preto: Simone Kashima, Evandra Strazza Rodrigues, Svetoslav Nanev Slavov, Elaine Vieira dos Santos, Rafael dos Santos Bezerra, Luiz Carlos Junior de Alcantara, Marta Giovanetti, Vagner Fonseca, Flavia Aburjaile, Rodrigo Tocantins Calado. |
| EPI_ISL_1795104, EPI_ISL_1795105 | LABORATORIO MUNICIPAL DE PIRACICABA | Instituto Butantan / ESALQ-Piracicaba | Instituto Butantan: Alexander Roberto Precioso, Dimas Tadeu Covas, Sandra Coccuzzo Sampaio, Maria Carolina Elias, José Salvatore Leister Patané, Vincent Louis Viala, Antonio Jorge Martins, Ricardo Haddad, Claudia Renata dos Santos Barros, Elaine Cristina Marqueze, Raul Machado Neto, Debora Botequio Moretti. Centro de Genômica Funcional da ESALQ: Luiz Lehmann Coutinho, Ricardo Augusto Brassaloti, Raquel de Lello Rocha Campos Cassano. NGS Soluções Genômicas: Pilar Drummond Sampaio Corrêa Mariani. FZEA-USP Pirassununga: Mirele Daiana Poleti, Jessika Cristina Chagas Lesbon, Elisangela Chicaroni Mattos, Heidge Fukumasu. USP-Botucatu: Rejane Maria Tommasini Grotto, Jayme A. Souza-Neto, Guilherme Targino Valente, Patricia Akemi Assato, Felipe Allan da Silva da Costa, Bianca Cechetto Carlos. Mendelics: Bibiana Santos, João Paulo Kitajima, Erika Freitas, David Schlesinger. Hemocentro Ribeirão Preto: Simone Kashima, Evandra Strazza Rodrigues, Svetoslav Nanev Slavov, Elaine Vieira dos Santos, Rafael dos Santos Bezerra, Luiz Carlos Junior de Alcantara, Marta Giovanetti, Vagner Fonseca, Flavia Aburjaile, Rodrigo Tocantins Calado. |
| EPI_ISL_1795108 | SECRETARIA DE SAUDE DE SAO PEDRO | Instituto Butantan / ESALQ-Piracicaba | Instituto Butantan: Alexander Roberto Precioso, Dimas Tadeu Covas, Sandra Coccuzzo Sampaio, Maria Carolina Elias, José Salvatore Leister Patané, Vincent Louis Viala, Antonio Jorge Martins, Ricardo Haddad, Claudia Renata dos Santos Barros, Elaine Cristina Marqueze, Raul Machado Neto, Debora Botequio Moretti. Centro de Genômica Funcional da ESALQ: Luiz Lehmann Coutinho, Ricardo Augusto Brassaloti, Raquel de Lello Rocha Campos Cassano. NGS Soluções Genômicas: Pilar Drummond Sampaio Corrêa Mariani. FZEA-USP Pirassununga: Mirele Daiana Poleti, Jessika Cristina Chagas Lesbon, Elisangela Chicaroni Mattos, Heidge Fukumasu. USP-Botucatu: Rejane Maria Tommasini Grotto, Jayme A. Souza-Neto, Guilherme Targino Valente, Patricia Akemi Assato, Felipe Allan da Silva da Costa, Bianca Cechetto Carlos. Mendelics: Bibiana Santos, João Paulo Kitajima, Erika Freitas, David Schlesinger. Hemocentro Ribeirão Preto: Simone Kashima, Evandra Strazza Rodrigues, Svetoslav Nanev Slavov, Elaine Vieira dos Santos, Rafael dos Santos Bezerra, Luiz Carlos Junior de Alcantara, Marta Giovanetti, Vagner Fonseca, Flavia Aburjaile, Rodrigo Tocantins Calado. |
| EPI_ISL_1795223 | HOSPITAL REGIONAL DE ITAPETININGA | Instituto Butantan / ESALQ-Piracicaba | Instituto Butantan: Alexander Roberto Precioso, Dimas Tadeu Covas, Sandra Coccuzzo Sampaio, Maria Carolina Elias, José Salvatore Leister Patané, Vincent Louis Viala, Antonio Jorge Martins, Ricardo Haddad, Claudia Renata dos Santos Barros, Elaine Cristina Marqueze, Raul Machado Neto, Debora Botequio Moretti. Centro de Genômica Funcional da ESALQ: Luiz Lehmann Coutinho, Ricardo Augusto Brassaloti, Raquel de Lello Rocha Campos Cassano. NGS Soluções Genômicas: Pilar Drummond Sampaio Corrêa Mariani. FZEA-USP Pirassununga: Mirele Daiana Poleti, Jessika Cristina Chagas Lesbon, Elisangela Chicaroni Mattos, Heidge Fukumasu. USP-Botucatu: Rejane Maria Tommasini Grotto, Jayme A. Souza-Neto, Guilherme Targino Valente, Patricia Akemi Assato, Felipe Allan da Silva da Costa, Bianca Cechetto Carlos. Mendelics: Bibiana Santos, João Paulo Kitajima, Erika Freitas, David Schlesinger. Hemocentro Ribeirão Preto: Simone Kashima, Evandra Strazza Rodrigues, Svetoslav Nanev Slavov, Elaine Vieira dos Santos, Rafael dos Santos Bezerra, Luiz Carlos Junior de Alcantara, Marta Giovanetti, Vagner Fonseca, Flavia Aburjaile, Rodrigo Tocantins Calado. |
| EPI_ISL_1795329 | CENTRO DE ESPECIALIDADES DE PRIMAVERA | Instituto Butantan / ESALQ-Piracicaba | Instituto Butantan: Alexander Roberto Precioso, Dimas Tadeu Covas, Sandra Coccuzzo Sampaio, Maria Carolina Elias, José Salvatore Leister Patané, Vincent Louis Viala, Antonio Jorge Martins, Ricardo Haddad, Claudia Renata dos Santos Barros, Elaine Cristina Marqueze, Raul Machado Neto, Debora Botequio Moretti. Centro de Genômica Funcional da ESALQ: Luiz Lehmann Coutinho, Ricardo Augusto Brassaloti, Raquel de Lello Rocha Campos Cassano. NGS Soluções Genômicas: Pilar Drummond Sampaio Corrêa Mariani. FZEA-USP Pirassununga: Mirele Daiana Poleti, Jessika Cristina Chagas Lesbon, Elisangela Chicaroni Mattos, Heidge Fukumasu. USP-Botucatu: Rejane Maria Tommasini Grotto, Jayme A. Souza-Neto, Guilherme Targino Valente, Patricia Akemi Assato, Felipe Allan da Silva da Costa, Bianca Cechetto Carlos. Mendelics: Bibiana Santos, João Paulo Kitajima, Erika Freitas, David Schlesinger. Hemocentro Ribeirão Preto: Simone Kashima, Evandra Strazza Rodrigues, Svetoslav Nanev Slavov, Elaine Vieira dos Santos, Rafael dos Santos Bezerra, Luiz Carlos Junior de Alcantara, Marta Giovanetti, Vagner Fonseca, Flavia Aburjaile, Rodrigo Tocantins Calado. |
| EPI_ISL_1858359, EPI_ISL_1858367, EPI_ISL_1858369, EPI_ISL_1858377, EPI_ISL_1858380, EPI_ISL_1858383, EPI_ISL_1858387, EPI_ISL_1858395, EPI_ISL_1858462, EPI_ISL_1858470, EPI_ISL_1858486, EPI_ISL_1858623, EPI_ISL_1858627, EPI_ISL_1858631, EPI_ISL_1858633, EPI_ISL_1858636, EPI_ISL_1858637, EPI_ISL_1858639, EPI_ISL_1858670, EPI_ISL_1858672, EPI_ISL_1858741 | | | |
| see above | Laboratorio Central Noel Nutels | Bioinformatics Laboratory / LNCC | Luiz G P de Almeida, Alessandra P Lamarca, Ronaldo da Silva F Jr, Liliane Cavalcante, Alexandra L Gerber, Ana Paula de C Guimarães, Douglas Terra Machado, Cassia Alves, Diana Mariani, Thais Felix Cruz, Mario Sergio Ribeiro, Silvia Carvalho, Flávio Dias da Silva, Marcio Henrique de Oliveira Garcia, Leandro Magalhães de Souza, Cristiane Gomes da Silva, Caio Luiz Pereira Ribeiro, Andréa Cony Cavalcanti, Claudia Maria Braga de Mello, Amilcar Tanuri, Ana Tereza R Vasconcelos |
| EPI_ISL_1858840 | Unidade de apoio ao diagnóstico da COVID - UNADIG | Bioinformatics Laboratory / LNCC | Luiz G P de Almeida, Alessandra P Lamarca, Ronaldo da Silva F Jr, Liliane Cavalcante, Alexandra L Gerber, Ana Paula de C Guimarães, Douglas Terra Machado, Cassia Alves, Diana Mariani, Thais Felix Cruz, Mario Sergio Ribeiro, Silvia Carvalho, Flávio Dias da Silva, Marcio Henrique de Oliveira Garcia, Leandro Magalhães de Souza, Cristiane Gomes da Silva, Caio Luiz Pereira Ribeiro, Andréa Cony Cavalcanti, Claudia Maria Braga de Mello, Amilcar Tanuri, Ana Tereza R Vasconcelos |
| EPI_ISL_1966113 | CS II EGIDIO BRUNHARA MORRO AGUDO | Instituto Butantan / Mendelics | Instituto Butantan: Dimas Tadeu Covas, Sandra Coccuzzo Sampaio, Maria Carolina Elias, José Salvatore Leister Patané, Vincent Louis Viala, Antonio Jorge Martins, Ricardo Haddad, Claudia Renata dos Santos Barros, Elaine Cristina Marqueze, Raul Machado Neto, Debora Botequio Moretti, Jardelina de |

| | | | |
|---|---|---|---|
| | | | Souza Todao Bernardino, Loyze Paola Oliveira de Lima, Luiz Aurelio de Campos Crispin. Centro de Genômica Funcional da ESALQ: Luiz Lehmann Coutinho, Ricardo Augusto Brassaloti, Raquel de Lello Rocha Campos Cassano. NGS Soluções Genômicas: Pilar Drummond Sampaio Corrêa Mariani. FZEA-USP Pirassununga: Mirele Daiana Poleti, Jessika Cristina Chagas Lesbon, Elisangela Chicaroni Mattos, Heidge Fukumasu. USP-Botucatu: Rejane Maria Tommasini Grotto, Jayme A. Souza-Neto, Guilherme Targino Valente, Patricia Akemi Assato, Felipe Allan da Silva da Costa, Bianca Cechetto Carlos. Mendelics: Bibiana Santos, João Paulo Kitajima, Erika Freitas, David Schlesinger. Hemocentro Ribeirão Preto: Simone Kashima, Evandra Strazza Rodrigues, Svetoslav Nanev Slavov, Elaine Vieira dos Santos, Rafael dos Santos Bezerra, Luiz Carlos Junior de Alcantara, Marta Giovanetti, Vagner Fonseca, Flavia Aburjaile, Rodrigo Tocantins Calado. FAMERP-SJRP: Cecília Artico Banho, Lívia Sacchetto, Fábio Sossai Possebon, Leila Sabrina Ullmann, Cíntia Bittar, Guilherme Campos, Helena Lage Ferreira, Jorge A. Petroli Marchesi, Maisa C. Pereira Parra, Marília Moraes, Paula Rahal, Paulo Inacio da Costa, João Pessoa Araújo Jr., Maurício Lacerda Nogueira. Prefeitura de Sao Paulo: Melissa Palmieri. |
| EPI_ISL_1966358 | SMS SECRETARIA MUNICIPAL DE SAUDE DE BOITUVA | Instituto Butantan / Mendelics | Instituto Butantan: Dimas Tadeu Covas, Sandra Coccuzzo Sampaio, Maria Carolina Elias, José Salvatore Leister Patané, Vincent Louis Viala, Antonio Jorge Martins, Ricardo Haddad, Claudia Renata dos Santos Barros, Elaine Cristina Marqueze, Raul Machado Neto, Debora Botequio Moretti, Jardelina de Souza Todao Bernardino, Loyze Paola Oliveira de Lima, Luiz Aurelio de Campos Crispin. Centro de Genômica Funcional da ESALQ: Luiz Lehmann Coutinho, Ricardo Augusto Brassaloti, Raquel de Lello Rocha Campos Cassano. NGS Soluções Genômicas: Pilar Drummond Sampaio Corrêa Mariani. FZEA-USP Pirassununga: Mirele Daiana Poleti, Jessika Cristina Chagas Lesbon, Elisangela Chicaroni Mattos, Heidge Fukumasu. USP-Botucatu: Rejane Maria Tommasini Grotto, Jayme A. Souza-Neto, Guilherme Targino Valente, Patricia Akemi Assato, Felipe Allan da Silva da Costa, Bianca Cechetto Carlos. Mendelics: Bibiana Santos, João Paulo Kitajima, Erika Freitas, David Schlesinger. Hemocentro Ribeirão Preto: Simone Kashima, Evandra Strazza Rodrigues, Svetoslav Nanev Slavov, Elaine Vieira dos Santos, Rafael dos Santos Bezerra, Luiz Carlos Junior de Alcantara, Marta Giovanetti, Vagner Fonseca, Flavia Aburjaile, Rodrigo Tocantins Calado. FAMERP-SJRP: Cecília Artico Banho, Lívia Sacchetto, Fábio Sossai Possebon, Leila Sabrina Ullmann, Cíntia Bittar, Guilherme Campos, Helena Lage Ferreira, Jorge A. Petroli Marchesi, Maisa C. Pereira Parra, Marília Moraes, Paula Rahal, Paulo Inacio da Costa, João Pessoa Araújo Jr., Maurício Lacerda Nogueira. Prefeitura de Sao Paulo: Melissa Palmieri. |
| EPI_ISL_1966448 | NUCLEO DE SAUDE VILA FALCAO DE BAURU | Instituto Butantan / Mendelics | Instituto Butantan: Dimas Tadeu Covas, Sandra Coccuzzo Sampaio, Maria Carolina Elias, José Salvatore Leister Patané, Vincent Louis Viala, Antonio Jorge Martins, Ricardo Haddad, Claudia Renata dos Santos Barros, Elaine Cristina Marqueze, Raul Machado Neto, Debora Botequio Moretti, Jardelina de Souza Todao Bernardino, Loyze Paola Oliveira de Lima, Luiz Aurelio de Campos Crispin. Centro de Genômica Funcional da ESALQ: Luiz Lehmann Coutinho, Ricardo Augusto Brassaloti, Raquel de Lello Rocha Campos Cassano. NGS Soluções Genômicas: Pilar Drummond Sampaio Corrêa Mariani. FZEA-USP Pirassununga: Mirele Daiana Poleti, Jessika Cristina Chagas Lesbon, Elisangela Chicaroni Mattos, Heidge Fukumasu. USP-Botucatu: Rejane Maria Tommasini Grotto, Jayme A. Souza-Neto, Guilherme Targino Valente, Patricia Akemi Assato, Felipe Allan da Silva da Costa, Bianca Cechetto Carlos. Mendelics: Bibiana Santos, João Paulo Kitajima, Erika Freitas, David Schlesinger. Hemocentro Ribeirão Preto: Simone Kashima, Evandra Strazza Rodrigues, Svetoslav Nanev Slavov, Elaine Vieira dos Santos, Rafael dos Santos Bezerra, Luiz Carlos Junior de Alcantara, Marta Giovanetti, Vagner Fonseca, Flavia Aburjaile, Rodrigo Tocantins Calado. FAMERP-SJRP: Cecília Artico Banho, Lívia Sacchetto, Fábio Sossai Possebon, Leila Sabrina Ullmann, Cíntia Bittar, Guilherme Campos, Helena Lage Ferreira, Jorge A. Petroli Marchesi, Maisa C. Pereira Parra, Marília Moraes, Paula Rahal, Paulo Inacio da Costa, João Pessoa Araújo Jr., Maurício Lacerda Nogueira. Prefeitura de Sao Paulo: Melissa Palmieri. |
| EPI_ISL_1966526 | USF PAULISTA FERNANDOPOLIS ANTONIO PIVATO | Instituto Butantan / FZEA-USP (Pirassununga) | Instituto Butantan: Dimas Tadeu Covas, Sandra Coccuzzo Sampaio, Maria Carolina Elias, José Salvatore Leister Patané, Vincent Louis Viala, Antonio Jorge Martins, Ricardo Haddad, Claudia Renata dos Santos Barros, Elaine Cristina Marqueze, Raul Machado Neto, Debora Botequio Moretti, Jardelina de Souza Todao Bernardino, Loyze Paola Oliveira de Lima, Luiz Aurelio de Campos Crispin. Centro de Genômica Funcional da ESALQ: Luiz Lehmann Coutinho, Ricardo Augusto Brassaloti, Raquel de Lello Rocha Campos Cassano. NGS Soluções Genômicas: Pilar Drummond Sampaio Corrêa Mariani. FZEA-USP Pirassununga: Mirele Daiana Poleti, Jessika Cristina Chagas Lesbon, Elisangela Chicaroni Mattos, Heidge Fukumasu. USP-Botucatu: Rejane Maria Tommasini Grotto, Jayme A. Souza-Neto, Guilherme Targino Valente, Patricia Akemi Assato, Felipe Allan da Silva da Costa, Bianca Cechetto Carlos. Mendelics: Bibiana Santos, João Paulo Kitajima, Erika Freitas, David Schlesinger. Hemocentro Ribeirão Preto: Simone Kashima, Evandra Strazza Rodrigues, Svetoslav Nanev Slavov, Elaine Vieira dos Santos, Rafael dos Santos Bezerra, Luiz Carlos Junior de Alcantara, Marta Giovanetti, Vagner Fonseca, Flavia Aburjaile, Rodrigo Tocantins Calado. FAMERP-SJRP: Cecília Artico Banho, Lívia Sacchetto, Fábio Sossai Possebon, Leila Sabrina Ullmann, Cíntia Bittar, Guilherme Campos, Helena Lage Ferreira, Jorge A. Petroli Marchesi, Maisa C. Pereira Parra, Marília Moraes, Paula Rahal, Paulo Inacio da Costa, João Pessoa Araújo Jr., Maurício Lacerda Nogueira. Prefeitura de Sao Paulo: Melissa Palmieri. |
| EPI_ISL_1966567, EPI_ISL_1966576, EPI_ISL_1966578, EPI_ISL_1966580, EPI_ISL_1966581, EPI_ISL_1966582, EPI_ISL_1966585 | NUCLEO DE SAUDE VILA FALCAO DE BAURU | Instituto Butantan / Mendelics | Instituto Butantan: Dimas Tadeu Covas, Sandra Coccuzzo Sampaio, Maria Carolina Elias, José Salvatore Leister Patané, Vincent Louis Viala, Antonio Jorge Martins, Ricardo Haddad, Claudia Renata dos Santos Barros, Elaine Cristina Marqueze, Raul Machado Neto, Debora Botequio Moretti, Jardelina de Souza Todao Bernardino, Loyze Paola Oliveira de Lima, Luiz Aurelio de Campos Crispin. Centro de Genômica Funcional da ESALQ: Luiz Lehmann Coutinho, Ricardo Augusto Brassaloti, Raquel de Lello Rocha Campos Cassano. NGS Soluções Genômicas: Pilar Drummond Sampaio Corrêa Mariani. FZEA-USP Pirassununga: Mirele Daiana Poleti, Jessika Cristina Chagas Lesbon, Elisangela Chicaroni Mattos, Heidge Fukumasu. USP-Botucatu: Rejane Maria Tommasini Grotto, Jayme A. Souza-Neto, Guilherme Targino Valente, Patricia Akemi Assato, Felipe Allan da Silva da Costa, Bianca Cechetto Carlos. Mendelics: Bibiana Santos, João Paulo Kitajima, Erika Freitas, David Schlesinger. Hemocentro Ribeirão Preto: Simone Kashima, Evandra Strazza Rodrigues, Svetoslav Nanev Slavov, Elaine Vieira dos Santos, Rafael dos Santos Bezerra, Luiz Carlos Junior de Alcantara, Marta Giovanetti, Vagner Fonseca, Flavia Aburjaile, Rodrigo Tocantins Calado. FAMERP-SJRP: Cecília Artico Banho, Lívia Sacchetto, Fábio Sossai Possebon, Leila Sabrina Ullmann, Cíntia Bittar, Guilherme Campos, Helena Lage Ferreira, Jorge A. Petroli Marchesi, Maisa C. Pereira Parra, Marília Moraes, Paula Rahal, Paulo Inacio da Costa, João Pessoa Araújo Jr., Maurício Lacerda Nogueira. Prefeitura de Sao Paulo: Melissa Palmieri. |
| EPI_ISL_1966658 | UPA UNIDADE DE PRONTO ATENDIMENTO 24 HORAS BOM JESUS | Instituto Butantan / Mendelics | Instituto Butantan: Dimas Tadeu Covas, Sandra Coccuzzo Sampaio, Maria Carolina Elias, José Salvatore Leister Patané, Vincent Louis Viala, Antonio Jorge Martins, Ricardo Haddad, Claudia Renata dos Santos Barros, Elaine Cristina Marqueze, Raul Machado Neto, Debora Botequio Moretti, Jardelina de Souza Todao Bernardino, Loyze Paola Oliveira de Lima, Luiz Aurelio de Campos Crispin. Centro de Genômica Funcional da ESALQ: Luiz Lehmann Coutinho, Ricardo Augusto Brassaloti, Raquel de Lello Rocha Campos Cassano. NGS Soluções Genômicas: Pilar Drummond Sampaio Corrêa Mariani. FZEA-USP Pirassununga: Mirele Daiana Poleti, Jessika Cristina Chagas Lesbon, Elisangela Chicaroni Mattos, Heidge Fukumasu. USP-Botucatu: Rejane Maria Tommasini Grotto, Jayme A. Souza-Neto, Guilherme Targino Valente, Patricia Akemi Assato, Felipe Allan da Silva da Costa, Bianca Cechetto Carlos. Mendelics: Bibiana Santos, João Paulo Kitajima, Erika Freitas, David Schlesinger. Hemocentro Ribeirão Preto: Simone Kashima, Evandra Strazza Rodrigues, Svetoslav Nanev Slavov, Elaine Vieira dos Santos, Rafael dos Santos Bezerra, Luiz Carlos Junior de Alcantara, Marta Giovanetti, Vagner Fonseca, Flavia Aburjaile, Rodrigo Tocantins Calado. FAMERP-SJRP: Cecília Artico Banho, Lívia Sacchetto, Fábio Sossai Possebon, Leila Sabrina Ullmann, Cíntia Bittar, Guilherme Campos, Helena Lage Ferreira, Jorge A. Petroli Marchesi, Maisa C. Pereira Parra, Marília Moraes, Paula Rahal, Paulo Inacio da Costa, João Pessoa Araújo Jr., Maurício Lacerda Nogueira. Prefeitura de Sao Paulo: Melissa Palmieri. |
| EPI_ISL_1966877 | UBS ATALAIA DR MARIO AGUIAR FILHO | Instituto Butantan / ESALQ-USP (Piracicaba) | Instituto Butantan: Dimas Tadeu Covas, Sandra Coccuzzo Sampaio, Maria Carolina Elias, José Salvatore Leister Patané, Vincent Louis Viala, Antonio Jorge Martins, Ricardo Haddad, Claudia Renata dos Santos Barros, Elaine Cristina Marqueze, Raul Machado Neto, Debora Botequio Moretti, Jardelina de Souza Todao Bernardino, Loyze Paola Oliveira de Lima, Luiz Aurelio de Campos Crispin. Centro de Genômica Funcional da ESALQ: Luiz Lehmann Coutinho, Ricardo Augusto Brassaloti, Raquel de Lello Rocha Campos Cassano. NGS Soluções Genômicas: Pilar Drummond Sampaio Corrêa Mariani. FZEA-USP Pirassununga: Mirele Daiana Poleti, Jessika Cristina Chagas Lesbon, Elisangela Chicaroni Mattos, Heidge Fukumasu. USP-Botucatu: Rejane Maria Tommasini Grotto, Jayme A. Souza-Neto, Guilherme Targino Valente, Patricia Akemi Assato, Felipe Allan da Silva da Costa, Bianca Cechetto Carlos. Mendelics: Bibiana Santos, João Paulo Kitajima, Erika Freitas, David Schlesinger. Hemocentro Ribeirão Preto: Simone Kashima, Evandra Strazza Rodrigues, Svetoslav Nanev Slavov, Elaine Vieira dos Santos, Rafael dos Santos Bezerra, Luiz Carlos Junior de Alcantara, Marta Giovanetti, Vagner Fonseca, Flavia Aburjaile, Rodrigo Tocantins Calado. FAMERP-SJRP: Cecília Artico Banho, Lívia Sacchetto, Fábio Sossai Possebon, Leila Sabrina Ullmann, Cíntia Bittar, Guilherme Campos, Helena Lage Ferreira, Jorge A. Petroli Marchesi, Maisa C. Pereira Parra, Marília Moraes, Paula Rahal, Paulo Inacio da Costa, João Pessoa Araújo Jr., Maurício Lacerda Nogueira. Prefeitura de Sao Paulo: Melissa Palmieri. |
| EPI_ISL_1966926 | HOSPITAL REGIONAL DE ITAPETININGA | Instituto Butantan / ESALQ-USP (Piracicaba) | Instituto Butantan: Dimas Tadeu Covas, Sandra Coccuzzo Sampaio, Maria Carolina Elias, José Salvatore Leister Patané, Vincent Louis Viala, Antonio Jorge Martins, Ricardo Haddad, Claudia Renata dos Santos Barros, Elaine Cristina Marqueze, Raul Machado Neto, Debora Botequio Moretti, Jardelina de Souza Todao Bernardino, Loyze Paola Oliveira de Lima, Luiz Aurelio de Campos Crispin. Centro de Genômica Funcional da ESALQ: Luiz Lehmann Coutinho, Ricardo Augusto Brassaloti, Raquel de Lello Rocha Campos Cassano. NGS Soluções Genômicas: Pilar Drummond Sampaio Corrêa Mariani. FZEA-USP Pirassununga: Mirele Daiana Poleti, Jessika Cristina Chagas Lesbon, Elisangela Chicaroni Mattos, Heidge Fukumasu. USP-Botucatu: Rejane Maria Tommasini Grotto, Jayme A. Souza-Neto, Guilherme Targino Valente, Patricia Akemi Assato, Felipe Allan da Silva da Costa, Bianca Cechetto Carlos. Mendelics: Bibiana Santos, João Paulo Kitajima, Erika Freitas, David Schlesinger. Hemocentro Ribeirão Preto: Simone Kashima, Evandra Strazza Rodrigues, Svetoslav Nanev Slavov, Elaine Vieira dos Santos, Rafael dos Santos Bezerra, Luiz Carlos Junior de Alcantara, Marta Giovanetti, Vagner Fonseca, Flavia Aburjaile, Rodrigo Tocantins Calado. FAMERP-SJRP: Cecília Artico Banho, Lívia Sacchetto, Fábio Sossai Possebon, Leila Sabrina |

| | | | |
|---|---|---|---|
| | | | Ullmann, Cíntia Bittar, Guilherme Campos, Helena Lage Ferreira, Jorge A. Petroli Marchesi, Maisa C. Pereira Parra, Marília Moraes, Paula Rahal, Paulo Inacio da Costa, João Pessoa Araújo Jr., Maurício Lacerda Nogueira. Prefeitura de Sao Paulo: Melissa Palmieri. |
| EPI_ISL_1967232 | UBS 09 JOAO CREVELARO BIRIGUI | Instituto Butantan / Mendelics | Instituto Butantan: Dimas Tadeu Covas, Sandra Coccuzzo Sampaio, Maria Carolina Elias, José Salvatore Leister Patané, Vincent Louis Viala, Antonio Jorge Martins, Ricardo Haddad, Claudia Renata dos Santos Barros, Elaine Cristina Marqueze, Raul Machado Neto, Debora Botequio Moretti, Jardelina de Souza Todao Bernardino, Loyze Paola Oliveira de Lima, Luiz Aurelio de Campos Crispin. Centro de Genômica Funcional da ESALQ: Luiz Lehmann Coutinho, Ricardo Augusto Brassaloti, Raquel de Lello Rocha Campos Cassano. NGS Soluções Genômicas: Pilar Drummond Sampaio Corrêa Mariani. FZEA-USP Pirassununga: Mirele Daiana Poleti, Jessika Cristina Chagas Lesbon, Elisangela Chicaroni Mattos, Heidge Fukumasu. USP-Botucatu: Rejane Maria Tommasini Grotto, Jayme A. Souza-Neto, Guilherme Targino Valente, Patricia Akemi Assato, Felipe Allan da Silva da Costa, Bianca Cechetto Carlos. Mendelics: Bibiana Santos, João Paulo Kitajima, Erika Freitas, David Schlesinger. Hemocentro Ribeirão Preto: Simone Kashima, Evandra Strazza Rodrigues, Svetoslav Nanev Slavov, Elaine Vieira dos Santos, Rafael dos Santos Bezerra, Luiz Carlos Junior de Alcantara, Marta Giovanetti, Vagner Fonseca, Flavia Aburjaile, Rodrigo Tocantins Calado. FAMERP-SJRP: Cecília Artico Banho, Lívia Sacchetto, Fábio Sossai Possebon, Leila Sabrina Ullmann, Cíntia Bittar, Guilherme Campos, Helena Lage Ferreira, Jorge A. Petroli Marchesi, Maisa C. Pereira Parra, Marília Moraes, Paula Rahal, Paulo Inacio da Costa, João Pessoa Araújo Jr., Maurício Lacerda Nogueira. Prefeitura de Sao Paulo: Melissa Palmieri. |
| EPI_ISL_1967284 | UBS RECREIO SAO JORGE | Instituto Butantan / Mendelics | Instituto Butantan: Dimas Tadeu Covas, Sandra Coccuzzo Sampaio, Maria Carolina Elias, José Salvatore Leister Patané, Vincent Louis Viala, Antonio Jorge Martins, Ricardo Haddad, Claudia Renata dos Santos Barros, Elaine Cristina Marqueze, Raul Machado Neto, Debora Botequio Moretti, Jardelina de Souza Todao Bernardino, Loyze Paola Oliveira de Lima, Luiz Aurelio de Campos Crispin. Centro de Genômica Funcional da ESALQ: Luiz Lehmann Coutinho, Ricardo Augusto Brassaloti, Raquel de Lello Rocha Campos Cassano. NGS Soluções Genômicas: Pilar Drummond Sampaio Corrêa Mariani. FZEA-USP Pirassununga: Mirele Daiana Poleti, Jessika Cristina Chagas Lesbon, Elisangela Chicaroni Mattos, Heidge Fukumasu. USP-Botucatu: Rejane Maria Tommasini Grotto, Jayme A. Souza-Neto, Guilherme Targino Valente, Patricia Akemi Assato, Felipe Allan da Silva da Costa, Bianca Cechetto Carlos. Mendelics: Bibiana Santos, João Paulo Kitajima, Erika Freitas, David Schlesinger. Hemocentro Ribeirão Preto: Simone Kashima, Evandra Strazza Rodrigues, Svetoslav Nanev Slavov, Elaine Vieira dos Santos, Rafael dos Santos Bezerra, Luiz Carlos Junior de Alcantara, Marta Giovanetti, Vagner Fonseca, Flavia Aburjaile, Rodrigo Tocantins Calado. FAMERP-SJRP: Cecília Artico Banho, Lívia Sacchetto, Fábio Sossai Possebon, Leila Sabrina Ullmann, Cíntia Bittar, Guilherme Campos, Helena Lage Ferreira, Jorge A. Petroli Marchesi, Maisa C. Pereira Parra, Marília Moraes, Paula Rahal, Paulo Inacio da Costa, João Pessoa Araújo Jr., Maurício Lacerda Nogueira. Prefeitura de Sao Paulo: Melissa Palmieri. |
| EPI_ISL_2003120, EPI_ISL_2003123 | Instituto Adolfo Lutz Central | Instituto Adolfo Lutz, Interdiciplinary Procedures Center, Strategic Laboratory | Claudio Tavares Sacchi, Claudia Regina Gonçalves, Erica Valessa Ramos Gomes, Karoline Rodrigues Campos, Caio Vinicius Dias Lopes, Leonardo Jose Tadeu de Araujo |
| EPI_ISL_2017480 | HLAGYN - Laboratorio de Imunologia de Transplantes de Goias | HLAGYN - Laboratorio de Imunologia de Transplantes de Goias | Fernando Antonio Vinhal dos Santos, Erika Lopes Rocha Batista, Alessandro Leonardo Alvares Magalhaes, Frederico Rodrigues Vinhal, Sabrina Sara Moreira Duarte, Danielle de Paiva Rezende, Lucas Carlos Gomes Pereira, Paola Cristina Resende Silva |
| EPI_ISL_2086603, EPI_ISL_2086611 | LABCOVID_HCPA | LABRESIS_HCPA | Wink PL, Martins AF, Volpato F, Monteiro F, Zavascki AP, Barth AL |
| EPI_ISL_2101444, EPI_ISL_2101450, EPI_ISL_2101587, EPI_ISL_2101589, EPI_ISL_2101593, EPI_ISL_2101607, EPI_ISL_2101689, EPI_ISL_2101720, EPI_ISL_2101739 | Laboratorio Central Noel Nutels | Bioinformatics Laboratory / LNCC | Luiz G P de Almeida, Alessandra P Lamarca, Ronaldo da Silva F Jr, Liliane Cavalcante, Alexandra L Gerber, Ana Paula de C Guimaraes, Douglas Terra Machado, Cassia Alves, Diana Mariani, Cintia Policarpo, Gleidson da Silva de Oliveira, Mario Sergio Ribeiro, Silvia Carvalho, Flavio Dias da Silva, Marcio Henrique de Oliveira Garcia, Leandro Magalhaes de Souza, Cristiane Gomes da Silva, Caio Luiz Pereira Ribeiro, Andrea Cony Cavalcanti, Claudia Maria Braga de Mello, Amilcar Tanuri, Ana Tereza R Vasconcelos |
| EPI_ISL_2139495, EPI_ISL_2139508, EPI_ISL_2139513, EPI_ISL_2139515, EPI_ISL_2139519, EPI_ISL_2139523, EPI_ISL_2139526, EPI_ISL_2139527, EPI_ISL_2139531, EPI_ISL_2139536 | Laboratorio Exame | Universidade Federal de Ciencias da Saude de Porto Alegre | Vinícius Bonetti Franceschi, Gabriel Dickin Caldana et al. |
| EPI_ISL_2157382 | Laboratorio Central de Saude Publica do Estado de Sergipe (LACEN/SE) | Laboratory of Respiratory Viruses and Measles, Oswaldo Cruz Institute, FIOCRUZ | Paola Resende, Luciana Appolinario, Fernando Motta, Anna Carolina Paixao, Ana Carolina Mendonca, Alice Sampaio Rocha, Tainá Moreira Martins Venas, Elisa Cavalcante Pereira, Renata Serrano Lopes, Cliomar Alves dos Santos, Marilda Siqueira on behalf of the Fiocruz COVID-19 Genomic Surveillance Network |
| EPI_ISL_2157396, EPI_ISL_2157399 | Laboratório Central de Saude Publica do Estado de Santa Catarina (LACEN/SC) | Laboratory of Respiratory Viruses and Measles, Oswaldo Cruz Institute, FIOCRUZ | Paola Resende, Luciana Appolinario, Fernando Motta, Anna Carolina Paixao, Ana Carolina Mendonca, Alice Sampaio Rocha, Taina Venas, Elisa Cavalcante Pereira, Renata Serrano Lopes, Darcita Buerger Rovaris, Sandra Bianchini Fernandes, Marilda Siqueira on behalf of the Fiocruz COVID-19 Genomic Surveillance Network |
| EPI_ISL_2157404 | Lboratorio Central de Saude Publica do Estado do Parana (LACEN/PR) | Laboratory of Respiratory Viruses and Measles, Oswaldo Cruz Institute, FIOCRUZ | Paola Resende, Luciana Appolinario, Fernando Motta, Anna Carolina Paixao, Ana Carolina Mendonca, Alice Sampaio Rocha, Taina Venas, Elisa Cavalcante Pereira, Renata Serrano Lopes, Irina Riediger, Marilda Siqueira on behalf of the Fiocruz COVID-19 Genomic Surveillance Network |
| EPI_ISL_2157407 | Laboratório Central de Saude Publica do Estado de Santa Catarina (LACEN/SC) | Laboratory of Respiratory Viruses and Measles, Oswaldo Cruz Institute, FIOCRUZ | Paola Resende, Luciana Appolinario, Fernando Motta, Anna Carolina Paixao, Ana Carolina Mendonca, Alice Sampaio Rocha, Taina Venas, Elisa Cavalcante Pereira, Renata Serrano Lopes, Darcita Buerger Rovaris, Sandra Bianchini Fernandes, Marilda Siqueira on behalf of the Fiocruz COVID-19 Genomic Surveillance Network |
| EPI_ISL_2157423 | UNIVERSIDADE FEDERAL DE VIÇOSA | Laboratory of Respiratory Viruses and Measles, Oswaldo Cruz Institute, FIOCRUZ | Paola Resende, Luciana Appolinario, Fernando Motta, Anna Carolina Paixao, Ana Carolina Mendonca, Alice Sampaio Rocha, Taina Venas, Elisa Cavalcante Pereira, Renata Serrano Lopes, Rubens Pasa, Marilda Siqueira on behalf of the Fiocruz COVID-19 Genomic Surveillance Network |
| EPI_ISL_2157426 | Laboratório Central de Saude Publica do Estado de Santa Catarina (LACEN/SC) | Laboratory of Respiratory Viruses and Measles, Oswaldo Cruz Institute, FIOCRUZ | Paola Resende, Luciana Appolinario, Fernando Motta, Anna Carolina Paixao, Ana Carolina Mendonca, Alice Sampaio Rocha, Taina Venas, Elisa Cavalcante Pereira, Renata Serrano Lopes, Darcita Buerger Rovaris, Sandra Bianchini Fernandes, Marilda Siqueira on behalf of the Fiocruz COVID-19 Genomic Surveillance Network |
| EPI_ISL_2157428, EPI_ISL_2157429, EPI_ISL_2157435, EPI_ISL_2157442, EPI_ISL_2157444 | Laboratorio Central de Saude Publica do Estado de Sergipe (LACEN/SE) | Laboratory of Respiratory Viruses and Measles, Oswaldo Cruz Institute, FIOCRUZ | Paola Resende, Luciana Appolinario, Fernando Motta, Anna Carolina Paixao, Ana Carolina Mendonca, Alice Sampaio Rocha, Tainá Moreira Martins Venas, Elisa Cavalcante Pereira, Renata Serrano Lopes, Cliomar Alves dos Santos, Marilda Siqueira on behalf of the Fiocruz COVID-19 Genomic Surveillance Network |
| EPI_ISL_2157447 | Laboratório Central de Saude Publica do Estado de Santa Catarina (LACEN/SC) | Laboratory of Respiratory Viruses and Measles, Oswaldo Cruz Institute, FIOCRUZ | Paola Resende, Luciana Appolinario, Fernando Motta, Anna Carolina Paixao, Ana Carolina Mendonca, Alice Sampaio Rocha, Taina Venas, Elisa Cavalcante Pereira, Renata Serrano Lopes, Darcita Buerger Rovaris, Sandra Bianchini Fernandes, Marilda Siqueira on behalf of the Fiocruz COVID-19 Genomic Surveillance Network |
| EPI_ISL_2157449, EPI_ISL_2157453, EPI_ISL_2157455, EPI_ISL_2157471, EPI_ISL_2157473, EPI_ISL_2157474, EPI_ISL_2157478 | Laboratorio Central de Saude Publica do Estado de Sergipe (LACEN/SE) | Laboratory of Respiratory Viruses and Measles, Oswaldo Cruz Institute, FIOCRUZ | Paola Resende, Luciana Appolinario, Fernando Motta, Anna Carolina Paixao, Ana Carolina Mendonca, Alice Sampaio Rocha, Tainá Moreira Martins Venas, Elisa Cavalcante Pereira, Renata Serrano Lopes, Cliomar Alves dos Santos, Marilda Siqueira on behalf of the Fiocruz COVID-19 Genomic Surveillance Network |
| EPI_ISL_2157487 | Lboratorio Central de Saude Publica do Estado do Parana (LACEN/PR) | Laboratory of Respiratory Viruses and Measles, Oswaldo Cruz Institute, FIOCRUZ | Paola Resende, Luciana Appolinario, Fernando Motta, Anna Carolina Paixao, Ana Carolina Mendonca, Alice Sampaio Rocha, Taina Venas, Elisa Cavalcante Pereira, Renata Serrano Lopes, Irina Riediger, Marilda Siqueira on behalf of the Fiocruz COVID-19 Genomic Surveillance Network |
| EPI_ISL_2157497, EPI_ISL_2157518 | Laboratório Central de Saude Publica do Estado de Santa Catarina (LACEN/SC) | Laboratory of Respiratory Viruses and Measles, Oswaldo Cruz Institute, FIOCRUZ | Paola Resende, Luciana Appolinario, Fernando Motta, Anna Carolina Paixao, Ana Carolina Mendonca, Alice Sampaio Rocha, Taina Venas, Elisa Cavalcante Pereira, Renata Serrano Lopes, Darcita Buerger Rovaris, Sandra Bianchini Fernandes, Marilda Siqueira on behalf of the Fiocruz COVID-19 Genomic Surveillance Network |
| EPI_ISL_2187879, EPI_ISL_2187911, EPI_ISL_2187952, EPI_ISL_2187964, EPI_ISL_2188002, EPI_ISL_2188008 | HLAGYN - Laboratorio de Imunologia de Transplantes de Goias | HLAGYN - Laboratorio de Imunologia de Transplantes de Goias | Fernando Antonio Vinhal dos Santos, Erika Lopes Rocha Batista, Alessandro Leonardo Alvares Magalhaes, Frederico Rodrigues Vinhal, Sabrina Sara Moreira Duarte, Lucas Carlos Gomes Pereira, Daniel Ferreira de Sousa |
| EPI_ISL_2196211, EPI_ISL_2196213, EPI_ISL_2196215, EPI_ISL_2196220 | Laboratorio Central de Saude Publica do Estado de Sergipe (LACEN/SE) | Laboratory of Respiratory Viruses and Measles, Oswaldo Cruz Institute, FIOCRUZ | Paola Resende, Luciana Appolinario, Fernando Motta, Anna Carolina Paixao, Ana Carolina Mendonca, Alice Sampaio Rocha, Tainá Moreira Martins Venas, Elisa Cavalcante Pereira, Renata Serrano Lopes, Cliomar Alves dos Santos, Marilda Siqueira on behalf of the Fiocruz COVID-19 Genomic Surveillance Network |
| EPI_ISL_2196267 | Laboratorio Central de Saude Publica do Estado Maranhao | Laboratory of Respiratory Viruses and Measles, Oswaldo Cruz | Paola Resende, Luciana Appolinario, Fernando Motta, Anna Carolina Paixao, Ana Carolina Mendonca, Alice Sampaio Rocha, Taina Venas, Elisa |

| | | | |
|---|---|---|---|
| | (LACEN-MA) | Institute, FIOCRUZ | Cavalcante Pereira, Renata Serrano Lopes, Lidio Gonçalves Lima Neto, Marilda Siqueira on behalf of the Fiocruz COVID-19 Genomic Surveillance Network |
| EPI_ISL_2209955 | SECRETARIA DE SAUDE DE SAO PEDRO | Instituto Butantan / FZEA-USP-Pirassununga | Dimas Tadeu Covas, Antonio Jorge Martins, Claudia Renata dos Santos Barros, David Schlesinger, Debora Botequio Moretti, Elaine Cristina Marqueze, Elaine Vieira Santos, Evandra Strazza Rodrigues, Heidge Fukumasu, Jayme Augusto de Souza-Neto, José Salvatore Leister Patané, Luiz Alcantara, Luiz Lehmann Coutinho, Maria Carolina Elias, Maurício Lacerda Nogueira, Rafael dos Santos Bezerra, Raul Machado Neto, Rejane Maria Tommasini Grotto, Ricardo Haddad, Sandra Coccuzzo Sampaio Vessoni, Simone Kashima, Svetoslav Nanev Slavov, Vincent Louis Viala |
| EPI_ISL_2241579 | Laboratório Central de Saúde Pública de Sergipe | Coordenação Geral de Laboratórios de Saúde Pública (CGLAB/DAEVS/SVS/MS) | Vagner Fonseca, et al. |
| EPI_ISL_2241581 | Laboratório Central de Saúde Pública da Paraiba | Coordenação Geral de Laboratórios de Saúde Pública (CGLAB/DAEVS/SVS/MS) | Vagner Fonseca, et al. |
| EPI_ISL_2241589, EPI_ISL_2241590 | Laboratório Central de Saúde Pública de Sergipe | Coordenação Geral de Laboratórios de Saúde Pública (CGLAB/DAEVS/SVS/MS) | Vagner Fonseca, et al. |
| EPI_ISL_2249357, EPI_ISL_2249361, EPI_ISL_2249365 | Laboratório Central de Saúde Pública do Rio Grande do Sul | Coordenação Geral de Laboratórios de Saúde Pública (CGLAB/DAEVS/SVS/MS) | Vagner Fonseca, et al. |
| EPI_ISL_2274080 | Laboratorio Central de Saude Publica do Estado Maranhao (LACEN-MA) | Laboratory of Respiratory Viruses and Measles, Oswaldo Cruz Institute, FIOCRUZ | Paola Resende, Luciana Appolinario, Fernando Motta, Anna Carolina Paixao, Ana Carolina Mendonca, Alice Sampaio Rocha, Taina Venas, Elisa Cavalcante Pereira, Renata Serrano Lopes, Lidio Gonçalves Lima Neto, Marilda Siqueira on behalf of the Fiocruz COVID-19 Genomic Surveillance Network |
| EPI_ISL_2274096 | Lboratorio Central de Saude Publica do Estado do Parana (LACEN/PR) | Laboratory of Respiratory Viruses and Measles, Oswaldo Cruz Institute, FIOCRUZ | Paola Resende, Luciana Appolinario, Fernando Motta, Anna Carolina Paixao, Ana Carolina Mendonca, Alice Sampaio Rocha, Taina Venas, Elisa Cavalcante Pereira, Renata Serrano Lopes, Irina Riediger, Marilda Siqueira on behalf of the Fiocruz COVID-19 Genomic Surveillance Network |
| EPI_ISL_2274108, EPI_ISL_2274109, EPI_ISL_2274115 | Laboratorio Central de Saude Publica do Estado do Rio Grande do Sul (LACEN-RS) | Laboratory of Respiratory Viruses and Measles, Oswaldo Cruz Institute, FIOCRUZ | Paola Resende, Luciana Appolinario, Fernando Motta, Anna Carolina Paixao, Ana Carolina Mendonca, Alice Sampaio Rocha, Taina Venas, Elisa Cavalcante Pereira, Renata Serrano Lopes, Tatiana Schaffer Gregianini, Richard Salvato, Marilda Siqueira on behalf of the Fiocruz COVID-19 Genomic Surveillance Network |
| EPI_ISL_2308409, EPI_ISL_2308456, EPI_ISL_2308471, EPI_ISL_2308473, EPI_ISL_2308475, EPI_ISL_2308478 | Laboratório Central de Saúde Pública de Sergipe | Coordenação Geral de Laboratórios de Saúde Pública (CGLAB/DAEVS/SVS/MS) | Vagner Fonseca, et al. |
| EPI_ISL_2344552 | HOSPITAL DOS FORNCEDORES | Instituto Butantan / ESALQ-Piracicaba | Dimas Tadeu Covas, Antonio Jorge Martins, Claudia Renata dos Santos Barros, David Schlesinger, Debora Botequio Moretti, Elaine Cristina Marqueze, Elaine Vieira Santos, Evandra Strazza Rodrigues, Heidge Fukumasu, Jayme Augusto de Souza-Neto, José Salvatore Leister Patané, Luiz Alcantara, Luiz Lehmann Coutinho, Maria Carolina Elias, Maurício Lacerda Nogueira, Rafael dos Santos Bezerra, Raul Machado Neto, Rejane Maria Tommasini Grotto, Ricardo Haddad, Sandra Coccuzzo Sampaio Vessoni, Simone Kashima, Svetoslav Nanev Slavov, Vincent Louis Viala |
| EPI_ISL_2344553, EPI_ISL_2344554 | LABORATORIO MUNICIPAL DE PIRACICABA | Instituto Butantan / ESALQ-Piracicaba | Dimas Tadeu Covas, Antonio Jorge Martins, Claudia Renata dos Santos Barros, David Schlesinger, Debora Botequio Moretti, Elaine Cristina Marqueze, Elaine Vieira Santos, Evandra Strazza Rodrigues, Heidge Fukumasu, Jayme Augusto de Souza-Neto, José Salvatore Leister Patané, Luiz Alcantara, Luiz Lehmann Coutinho, Maria Carolina Elias, Maurício Lacerda Nogueira, Rafael dos Santos Bezerra, Raul Machado Neto, Rejane Maria Tommasini Grotto, Ricardo Haddad, Sandra Coccuzzo Sampaio Vessoni, Simone Kashima, Svetoslav Nanev Slavov, Vincent Louis Viala |
| EPI_ISL_2344662 | SECRETARIA DE SAUDE DE SAO PEDRO | Instituto Butantan / ESALQ-Piracicaba | Dimas Tadeu Covas, Antonio Jorge Martins, Claudia Renata dos Santos Barros, David Schlesinger, Debora Botequio Moretti, Elaine Cristina Marqueze, Elaine Vieira Santos, Evandra Strazza Rodrigues, Heidge Fukumasu, Jayme Augusto de Souza-Neto, José Salvatore Leister Patané, Luiz Alcantara, Luiz Lehmann Coutinho, Maria Carolina Elias, Maurício Lacerda Nogueira, Rafael dos Santos Bezerra, Raul Machado Neto, Rejane Maria Tommasini Grotto, Ricardo Haddad, Sandra Coccuzzo Sampaio Vessoni, Simone Kashima, Svetoslav Nanev Slavov, Vincent Louis Viala |
| EPI_ISL_2344851 | LABORATORIO MUNICIPAL DE PIRACICABA | Instituto Butantan / ESALQ-Piracicaba | Dimas Tadeu Covas, Antonio Jorge Martins, Claudia Renata dos Santos Barros, David Schlesinger, Debora Botequio Moretti, Elaine Cristina Marqueze, Elaine Vieira Santos, Evandra Strazza Rodrigues, Heidge Fukumasu, Jayme Augusto de Souza-Neto, José Salvatore Leister Patané, Luiz Alcantara, Luiz Lehmann Coutinho, Maria Carolina Elias, Maurício Lacerda Nogueira, Rafael dos Santos Bezerra, Raul Machado Neto, Rejane Maria Tommasini Grotto, Ricardo Haddad, Sandra Coccuzzo Sampaio Vessoni, Simone Kashima, Svetoslav Nanev Slavov, Vincent Louis Viala |
| EPI_ISL_2344900 | HOSPITAL E PRONTO SOCORRO PORTINARI | Instituto Butantan | Dimas Tadeu Covas, Antonio Jorge Martins, Claudia Renata dos Santos Barros, David Schlesinger, Debora Botequio Moretti, Elaine Cristina Marqueze, Elaine Vieira Santos, Evandra Strazza Rodrigues, Heidge Fukumasu, Jayme Augusto de Souza-Neto, José Salvatore Leister Patané, Luiz Alcantara, Luiz Lehmann Coutinho, Maria Carolina Elias, Maurício Lacerda Nogueira, Rafael dos Santos Bezerra, Raul Machado Neto, Rejane Maria Tommasini Grotto, Ricardo Haddad, Sandra Coccuzzo Sampaio Vessoni, Simone Kashima, Svetoslav Nanev Slavov, Vincent Louis Viala |
| EPI_ISL_2344915, EPI_ISL_2344985 | LABORATORIO MUNICIPAL DE PIRACICABA | Instituto Butantan / ESALQ-Piracicaba | Dimas Tadeu Covas, Antonio Jorge Martins, Claudia Renata dos Santos Barros, David Schlesinger, Debora Botequio Moretti, Elaine Cristina Marqueze, Elaine Vieira Santos, Evandra Strazza Rodrigues, Heidge Fukumasu, Jayme Augusto de Souza-Neto, José Salvatore Leister Patané, Luiz Alcantara, Luiz Lehmann Coutinho, Maria Carolina Elias, Maurício Lacerda Nogueira, Rafael dos Santos Bezerra, Raul Machado Neto, Rejane Maria Tommasini Grotto, Ricardo Haddad, Sandra Coccuzzo Sampaio Vessoni, Simone Kashima, Svetoslav Nanev Slavov, Vincent Louis Viala |
| EPI_ISL_2345026 | HOSP MUN JOSANIAS CASTANHA BRAGA | Instituto Butantan | Dimas Tadeu Covas, Antonio Jorge Martins, Claudia Renata dos Santos Barros, David Schlesinger, Debora Botequio Moretti, Elaine Cristina Marqueze, Elaine Vieira Santos, Evandra Strazza Rodrigues, Heidge Fukumasu, Jayme Augusto de Souza-Neto, José Salvatore Leister Patané, Luiz Alcantara, Luiz Lehmann Coutinho, Maria Carolina Elias, Maurício Lacerda Nogueira, Rafael dos Santos Bezerra, Raul Machado Neto, Rejane Maria Tommasini Grotto, Ricardo Haddad, Sandra Coccuzzo Sampaio Vessoni, Simone Kashima, Svetoslav Nanev Slavov, Vincent Louis Viala |
| EPI_ISL_2345181 | NUCLEO SAUDE III N HABITACIONAL PRESIDENTE GEISEL BAURU | Instituto Butantan / ESALQ-Piracicaba | Dimas Tadeu Covas, Antonio Jorge Martins, Claudia Renata dos Santos Barros, David Schlesinger, Debora Botequio Moretti, Elaine Cristina Marqueze, Elaine Vieira Santos, Evandra Strazza Rodrigues, Heidge Fukumasu, Jayme Augusto de Souza-Neto, José Salvatore Leister Patané, Luiz Alcantara, Luiz Lehmann Coutinho, Maria Carolina Elias, Maurício Lacerda Nogueira, Rafael dos Santos Bezerra, Raul Machado Neto, Rejane Maria Tommasini Grotto, Ricardo Haddad, Sandra Coccuzzo Sampaio Vessoni, Simone Kashima, Svetoslav Nanev Slavov, Vincent Louis Viala |
| EPI_ISL_2345207 | NUCLEO DE SAUDE VILA FALCAO DE BAURU | Instituto Butantan / ESALQ-Piracicaba | Dimas Tadeu Covas, Antonio Jorge Martins, Claudia Renata dos Santos Barros, David Schlesinger, Debora Botequio Moretti, Elaine Cristina Marqueze, Elaine Vieira Santos, Evandra Strazza Rodrigues, Heidge Fukumasu, Jayme Augusto de Souza-Neto, José Salvatore Leister Patané, Luiz Alcantara, Luiz Lehmann Coutinho, Maria Carolina Elias, Maurício Lacerda Nogueira, Rafael dos Santos Bezerra, Raul Machado Neto, Rejane Maria Tommasini Grotto, Ricardo Haddad, Sandra Coccuzzo Sampaio Vessoni, Simone Kashima, Svetoslav Nanev Slavov, Vincent Louis Viala |
| EPI_ISL_2345267 | LABORATORIO MUNICIPAL DE PIRACICABA | Instituto Butantan / ESALQ-Piracicaba | Dimas Tadeu Covas, Antonio Jorge Martins, Claudia Renata dos Santos Barros, David Schlesinger, Debora Botequio Moretti, Elaine Cristina Marqueze, Elaine Vieira Santos, Evandra Strazza Rodrigues, Heidge Fukumasu, Jayme Augusto de Souza-Neto, José Salvatore Leister Patané, Luiz Alcantara, Luiz Lehmann Coutinho, Maria Carolina Elias, Maurício Lacerda Nogueira, Rafael dos Santos Bezerra, Raul Machado Neto, Rejane Maria Tommasini Grotto, Ricardo Haddad, Sandra Coccuzzo Sampaio Vessoni, Simone Kashima, Svetoslav Nanev Slavov, Vincent Louis Viala |
| EPI_ISL_2345268 | UPA IPIRANGA | Instituto Butantan / ESALQ-Piracicaba | Dimas Tadeu Covas, Antonio Jorge Martins, Claudia Renata dos Santos Barros, David Schlesinger, Debora Botequio Moretti, Elaine Cristina Marqueze, Elaine Vieira Santos, Evandra Strazza Rodrigues, Heidge Fukumasu, Jayme Augusto de Souza-Neto, José Salvatore Leister Patané, Luiz Alcantara, Luiz Lehmann Coutinho, Maria Carolina Elias, Maurício Lacerda Nogueira, Rafael dos Santos Bezerra, Raul Machado Neto, Rejane Maria Tommasini Grotto, Ricardo Haddad, Sandra Coccuzzo Sampaio Vessoni, Simone Kashima, Svetoslav Nanev Slavov, Vincent Louis Viala |
| EPI_ISL_2345280, EPI_ISL_2345282 | NUCLEO DE SAUDE VILA FALCAO DE BAURU | Instituto Butantan / ESALQ-Piracicaba | Dimas Tadeu Covas, Antonio Jorge Martins, Claudia Renata dos Santos Barros, David Schlesinger, Debora Botequio Moretti, Elaine Cristina Marqueze, Elaine Vieira Santos, Evandra Strazza Rodrigues, Heidge Fukumasu, Jayme Augusto de Souza-Neto, José Salvatore Leister Patané, Luiz Alcantara, Luiz Lehmann Coutinho, Maria Carolina Elias, Maurício Lacerda Nogueira, Rafael dos Santos Bezerra, Raul Machado Neto, Rejane Maria Tommasini Grotto, Ricardo Haddad, Sandra Coccuzzo Sampaio Vessoni, Simone Kashima, Svetoslav Nanev Slavov, Vincent Louis Viala |
| EPI_ISL_2345453 | HOSPITAL REGIONAL DE ITAPETININGA | Instituto Butantan / ESALQ-Piracicaba | Dimas Tadeu Covas, Antonio Jorge Martins, Claudia Renata dos Santos Barros, David Schlesinger, Debora Botequio Moretti, Elaine Cristina Marqueze, Elaine Vieira Santos, Evandra Strazza Rodrigues, Heidge Fukumasu, Jayme Augusto de Souza-Neto, José Salvatore Leister Patané, Luiz Alcantara, Luiz Lehmann Coutinho, Maria Carolina Elias, Maurício Lacerda Nogueira, Rafael dos Santos Bezerra, Raul Machado Neto, Rejane Maria Tommasini Grotto, |

| | | | |
|---|---|---|---|
| | | | Ricardo Haddad, Sandra Coccuzzo Sampaio Vessoni, Simone Kashima, Svetoslav Nanev Slavov, Vincent Louis Viala |
| EPI_ISL_2345590 | CENTRO DE ESPECIALIDADES DE PRIMAVERA | Instituto Butantan / ESALQ-Piracicaba | Dimas Tadeu Covas, Antonio Jorge Martins, Claudia Renata dos Santos Barros, David Schlesinger, Debora Botequio Moretti, Elaine Cristina Marqueze, Elaine Vieira Santos, Evandra Strazza Rodrigues, Heidge Fukumasu, Jayme Augusto de Souza-Neto, José Salvatore Leister Patané, Luiz Alcantara, Luiz Lehmann Coutinho, Maria Carolina Elias, Maurício Lacerda Nogueira, Rafael dos Santos Bezerra, Raul Machado Neto, Rejane Maria Tommasini Grotto, Ricardo Haddad, Sandra Coccuzzo Sampaio Vessoni, Simone Kashima, Svetoslav Nanev Slavov, Vincent Louis Viala |
| EPI_ISL_2345708 | SMS SECRETARIA MUNICIPAL DE SAUDE DE BOITUVA | Instituto Butantan / Mendelics | Dimas Tadeu Covas, Antonio Jorge Martins, Claudia Renata dos Santos Barros, David Schlesinger, Debora Botequio Moretti, Elaine Cristina Marqueze, Elaine Vieira Santos, Evandra Strazza Rodrigues, Heidge Fukumasu, Jayme Augusto de Souza-Neto, José Salvatore Leister Patané, Luiz Alcantara, Luiz Lehmann Coutinho, Maria Carolina Elias, Maurício Lacerda Nogueira, Rafael dos Santos Bezerra, Raul Machado Neto, Rejane Maria Tommasini Grotto, Ricardo Haddad, Sandra Coccuzzo Sampaio Vessoni, Simone Kashima, Svetoslav Nanev Slavov, Vincent Louis Viala |
| EPI_ISL_2345715 | CENTRO DE ESPECIALIDADES DE PRIMAVERA | Instituto Butantan / Mendelics | Dimas Tadeu Covas, Antonio Jorge Martins, Claudia Renata dos Santos Barros, David Schlesinger, Debora Botequio Moretti, Elaine Cristina Marqueze, Elaine Vieira Santos, Evandra Strazza Rodrigues, Heidge Fukumasu, Jayme Augusto de Souza-Neto, José Salvatore Leister Patané, Luiz Alcantara, Luiz Lehmann Coutinho, Maria Carolina Elias, Maurício Lacerda Nogueira, Rafael dos Santos Bezerra, Raul Machado Neto, Rejane Maria Tommasini Grotto, Ricardo Haddad, Sandra Coccuzzo Sampaio Vessoni, Simone Kashima, Svetoslav Nanev Slavov, Vincent Louis Viala |
| EPI_ISL_2345723 | UNIDADE BASICA DE SAUDE JARDIM DO LAGO | Instituto Butantan / Mendelics | Dimas Tadeu Covas, Antonio Jorge Martins, Claudia Renata dos Santos Barros, David Schlesinger, Debora Botequio Moretti, Elaine Cristina Marqueze, Elaine Vieira Santos, Evandra Strazza Rodrigues, Heidge Fukumasu, Jayme Augusto de Souza-Neto, José Salvatore Leister Patané, Luiz Alcantara, Luiz Lehmann Coutinho, Maria Carolina Elias, Maurício Lacerda Nogueira, Rafael dos Santos Bezerra, Raul Machado Neto, Rejane Maria Tommasini Grotto, Ricardo Haddad, Sandra Coccuzzo Sampaio Vessoni, Simone Kashima, Svetoslav Nanev Slavov, Vincent Louis Viala |
| EPI_ISL_2345755 | SECRETARIA MUNICIPAL DE SAUDE SOROCABA | Instituto Butantan / Mendelics | Dimas Tadeu Covas, Antonio Jorge Martins, Claudia Renata dos Santos Barros, David Schlesinger, Debora Botequio Moretti, Elaine Cristina Marqueze, Elaine Vieira Santos, Evandra Strazza Rodrigues, Heidge Fukumasu, Jayme Augusto de Souza-Neto, José Salvatore Leister Patané, Luiz Alcantara, Luiz Lehmann Coutinho, Maria Carolina Elias, Maurício Lacerda Nogueira, Rafael dos Santos Bezerra, Raul Machado Neto, Rejane Maria Tommasini Grotto, Ricardo Haddad, Sandra Coccuzzo Sampaio Vessoni, Simone Kashima, Svetoslav Nanev Slavov, Vincent Louis Viala |
| EPI_ISL_2345811 | CENTRO DE ESPECIALIDADES DE PRIMAVERA | Instituto Butantan / Mendelics | Dimas Tadeu Covas, Antonio Jorge Martins, Claudia Renata dos Santos Barros, David Schlesinger, Debora Botequio Moretti, Elaine Cristina Marqueze, Elaine Vieira Santos, Evandra Strazza Rodrigues, Heidge Fukumasu, Jayme Augusto de Souza-Neto, José Salvatore Leister Patané, Luiz Alcantara, Luiz Lehmann Coutinho, Maria Carolina Elias, Maurício Lacerda Nogueira, Rafael dos Santos Bezerra, Raul Machado Neto, Rejane Maria Tommasini Grotto, Ricardo Haddad, Sandra Coccuzzo Sampaio Vessoni, Simone Kashima, Svetoslav Nanev Slavov, Vincent Louis Viala |
| EPI_ISL_2345828 | UPA MARY DOTA | Instituto Butantan / Mendelics | Dimas Tadeu Covas, Antonio Jorge Martins, Claudia Renata dos Santos Barros, David Schlesinger, Debora Botequio Moretti, Elaine Cristina Marqueze, Elaine Vieira Santos, Evandra Strazza Rodrigues, Heidge Fukumasu, Jayme Augusto de Souza-Neto, José Salvatore Leister Patané, Luiz Alcantara, Luiz Lehmann Coutinho, Maria Carolina Elias, Maurício Lacerda Nogueira, Rafael dos Santos Bezerra, Raul Machado Neto, Rejane Maria Tommasini Grotto, Ricardo Haddad, Sandra Coccuzzo Sampaio Vessoni, Simone Kashima, Svetoslav Nanev Slavov, Vincent Louis Viala |
| EPI_ISL_2345874 | CENTRO DE APOIO EPIDEMIOLOGICO | Instituto Butantan / Mendelics | Dimas Tadeu Covas, Antonio Jorge Martins, Claudia Renata dos Santos Barros, David Schlesinger, Debora Botequio Moretti, Elaine Cristina Marqueze, Elaine Vieira Santos, Evandra Strazza Rodrigues, Heidge Fukumasu, Jayme Augusto de Souza-Neto, José Salvatore Leister Patané, Luiz Alcantara, Luiz Lehmann Coutinho, Maria Carolina Elias, Maurício Lacerda Nogueira, Rafael dos Santos Bezerra, Raul Machado Neto, Rejane Maria Tommasini Grotto, Ricardo Haddad, Sandra Coccuzzo Sampaio Vessoni, Simone Kashima, Svetoslav Nanev Slavov, Vincent Louis Viala |
| EPI_ISL_2346088, EPI_ISL_2346090 | Prefeirtura de SP | Instituto Butantan | Dimas Tadeu Covas, Antonio Jorge Martins, Claudia Renata dos Santos Barros, David Schlesinger, Debora Botequio Moretti, Elaine Cristina Marqueze, Elaine Vieira Santos, Evandra Strazza Rodrigues, Heidge Fukumasu, Jayme Augusto de Souza-Neto, José Salvatore Leister Patané, Luiz Alcantara, Luiz Lehmann Coutinho, Maria Carolina Elias, Maurício Lacerda Nogueira, Rafael dos Santos Bezerra, Raul Machado Neto, Rejane Maria Tommasini Grotto, Ricardo Haddad, Sandra Coccuzzo Sampaio Vessoni, Simone Kashima, Svetoslav Nanev Slavov, Vincent Louis Viala |
| EPI_ISL_2378653, EPI_ISL_2378681 | Instituto de Biotecnologia - UNESP-Botucatu-SP | Instituto de Biotecnologia - UNESP-Botucatu-SP | Fábio Sossai Possebon; Leila Sabrina Ullmann; Cecília Artico Banho; Cíntia Bittar; Guilherme Campos; Helena Lage Ferreira; Jorge A. Petroli Marchesi; Lívia Sacchetto; Maisa C. Pereira Parra; Marília Moraes; Maurício L. Nogueira; Paula Rahal; Paulo Inacio da Costa; João Pessoa Araújo Jr. |
| EPI_ISL_2378745 | HOSPITAL DO RIM E HIPERTENSAO | Instituto Butantan | Dimas Tadeu Covas, Antonio Jorge Martins, Claudia Renata dos Santos Barros, David Schlesinger, Debora Botequio Moretti, Elaine Cristina Marqueze, Elaine Vieira Santos, Evandra Strazza Rodrigues, Heidge Fukumasu, Jayme Augusto de Souza-Neto, José Salvatore Leister Patané, Luiz Alcantara, Luiz Lehmann Coutinho, Maria Carolina Elias, Maurício Lacerda Nogueira, Rafael dos Santos Bezerra, Raul Machado Neto, Rejane Maria Tommasini Grotto, Ricardo Haddad, Sandra Coccuzzo Sampaio Vessoni, Simone Kashima, Svetoslav Nanev Slavov, Vincent Louis Viala |
| EPI_ISL_2383033 | LABORATORIO MUNICIPAL DE PIRACICABA | Instituto Butantan | Dimas Tadeu Covas, Antonio Jorge Martins, Claudia Renata dos Santos Barros, David Schlesinger, Debora Botequio Moretti, Elaine Cristina Marqueze, Elaine Vieira Santos, Evandra Strazza Rodrigues, Heidge Fukumasu, Jayme Augusto de Souza-Neto, José Salvatore Leister Patané, Luiz Alcantara, Luiz Lehmann Coutinho, Maria Carolina Elias, Maurício Lacerda Nogueira, Rafael dos Santos Bezerra, Raul Machado Neto, Rejane Maria Tommasini Grotto, Ricardo Haddad, Sandra Coccuzzo Sampaio Vessoni, Simone Kashima, Svetoslav Nanev Slavov, Vincent Louis Viala |
| EPI_ISL_2383035 | UNIDADE BASICA DE SAUDE DR MATHEUS GABRIEL BONASSA | Instituto Butantan | Dimas Tadeu Covas, Antonio Jorge Martins, Claudia Renata dos Santos Barros, David Schlesinger, Debora Botequio Moretti, Elaine Cristina Marqueze, Elaine Vieira Santos, Evandra Strazza Rodrigues, Heidge Fukumasu, Jayme Augusto de Souza-Neto, José Salvatore Leister Patané, Luiz Alcantara, Luiz Lehmann Coutinho, Maria Carolina Elias, Maurício Lacerda Nogueira, Rafael dos Santos Bezerra, Raul Machado Neto, Rejane Maria Tommasini Grotto, Ricardo Haddad, Sandra Coccuzzo Sampaio Vessoni, Simone Kashima, Svetoslav Nanev Slavov, Vincent Louis Viala |
| EPI_ISL_2385439, EPI_ISL_2385449, EPI_ISL_2385481, EPI_ISL_2385520, EPI_ISL_2385524, EPI_ISL_2385525, EPI_ISL_2385526, EPI_ISL_2385555, EPI_ISL_2385564, EPI_ISL_2385565, EPI_ISL_2385566, EPI_ISL_2385588, EPI_ISL_2385650, EPI_ISL_2385660 | | | |
| see above | Laboratorio Central Noel Nutels | Bioinformatics Laboratory / LNCC | Luiz G P de Almeida, Alessandra P Lamarca, Ronaldo da Silva F Jr, Liliane Cavalcante, Alexandra L Gerber, Ana Paula de C Guimaraes, Douglas Terra Machado, Cassia Alves, Diana Mariani, Cintia Policarpo, Gleidson da Silva de Oliveira, Mario Sergio Ribeiro, Silvia Carvalho, Flavio Dias da Silva, Marcio Henrique de Oliveira Garcia, Leandro Magalhaes de Souza, Cristiane Gomes da Silva, Caio Luiz Pereira Ribeiro, Andrea Cony Cavalcanti, Claudia Maria Braga de Mello, Amilcar Tanuri, Ana Tereza R Vasconcelos |
| EPI_ISL_2431437, EPI_ISL_2431843 | Laboratório de Microbiologia Molecular - Universidade FEEVALE | Molecular Microbiology Laboratory | Alana Witt Hansen, Fágner Henrique Heldt, Fernando Rosado Spilki, Flávio Silveira, Juliana Schons Gularte, Juliane Deise Fleck, Mariana Soares da Silva, Meriane Demoliner, Matheus Nunes Weber, Paula Rodrigues de Almeida, Micheli Filippi. |
| EPI_ISL_2443542 | Lboratorio Central de Saude Publica do Estado do Parana (LACEN/PR) | Laboratory of Respiratory Viruses and Measles, Oswaldo Cruz Institute, FIOCRUZ | Paola Resende, Luciana Appolinario, Fernando Motta, Anna Carolina Paixao, Ana Carolina Mendonca, Alice Sampaio Rocha, Taina Venas, Elisa Cavalcante Pereira, Renata Serrano Lopes, Irina Riediger, Marilda Siqueira on behalf of the Fiocruz COVID-19 Genomic Surveillance Network |
| EPI_ISL_2443591 | Laboratory of Respiratory Viruses and Measles, Oswaldo Cruz Institute, FIOCRUZ | Laboratory of Respiratory Viruses and Measles, Oswaldo Cruz Institute, FIOCRUZ | Paola Resende, Luciana Appolinario, Fernando Motta, Anna Carolina Paixao, Ana Carolina Mendonca, Alice Sampaio Rocha, Taina Venas, Elisa Cavalcante Pereira, Renata Serrano Lopes, Marilda Siqueira on behalf of the Fiocruz COVID-19 Genomic Surveillance Network |
| EPI_ISL_2443673 | Laboratorio Central de Saude Publica do Estado do Rio Grande do Sul (LACEN-RS) | Laboratory of Respiratory Viruses and Measles, Oswaldo Cruz Institute, FIOCRUZ | Paola Resende, Luciana Appolinario, Fernando Motta, Anna Carolina Paixao, Ana Carolina Mendonca, Alice Sampaio Rocha, Taina Venas, Elisa Cavalcante Pereira, Renata Serrano Lopes, Tatiana Schaffer Gregianini, Richard Salvato, Marilda Siqueira on behalf of the Fiocruz COVID-19 Genomic Surveillance Network |
| EPI_ISL_2443675, EPI_ISL_2443712 | Labortorio Central de Saude Publica do Estado de Santa Catarina (LACEN/SC) | Laboratory of Respiratory Viruses and Measles, Oswaldo Cruz Institute, FIOCRUZ | Paola Resende, Luciana Appolinario, Fernando Motta, Anna Carolina Paixao, Ana Carolina Mendonca, Alice Sampaio Rocha, Taina Venas, Elisa Cavalcante Pereira, Renata Serrano Lopes, Darcita Buerger Rovaris, Sandra Bianchini Fernandes, Marilda Siqueira on behalf of the Fiocruz COVID-19 Genomic Surveillance Network |
| EPI_ISL_2445168 | UNIDADE REFERENCIAL SUDOESTE | Instituto Butantan | Dimas Tadeu Covas, Antonio Jorge Martins, Claudia Renata dos Santos Barros, David Schlesinger, Debora Botequio Moretti, Elaine Cristina Marqueze, Elaine Vieira Santos, Evandra Strazza Rodrigues, Heidge Fukumasu, Jayme Augusto de Souza-Neto, José Salvatore Leister Patané, Luiz Alcantara, Luiz Lehmann Coutinho, Maria Carolina Elias, Maurício Lacerda Nogueira, Rafael dos Santos Bezerra, Raul Machado Neto, Rejane Maria Tommasini Grotto, Ricardo Haddad, Sandra Coccuzzo Sampaio Vessoni, Simone Kashima, Svetoslav Nanev Slavov, Vincent Louis Viala |
| EPI_ISL_2445275 | UPA VILA SANTA CATARINA | Instituto Butantan | Dimas Tadeu Covas, Antonio Jorge Martins, Claudia Renata dos Santos Barros, David Schlesinger, Debora Botequio Moretti, Elaine Cristina Marqueze, Elaine Vieira Santos, Evandra Strazza Rodrigues, Heidge Fukumasu, Jayme Augusto de Souza-Neto, José Salvatore Leister Patané, Luiz Alcantara, Luiz Lehmann Coutinho, Maria Carolina Elias, Maurício Lacerda Nogueira, Rafael dos Santos Bezerra, Raul Machado Neto, Rejane Maria Tommasini Grotto, Ricardo Haddad, Sandra Coccuzzo Sampaio Vessoni, Simone Kashima, Svetoslav Nanev Slavov, Vincent Louis Viala |

| | | | |
|---|---|---|---|
| EPI_ISL_2445289 | VIGILANCIA SANITARIA E VIG EPIDEMIOLOGICA DE ITAPEVI | Instituto Butantan | Dimas Tadeu Covas, Antonio Jorge Martins, Claudia Renata dos Santos Barros, David Schlesinger, Debora Botequio Moretti, Elaine Cristina Marqueze, Elaine Vieira Santos, Evandra Strazza Rodrigues, Heidge Fukumasu, Jayme Augusto de Souza-Neto, José Salvatore Leister Patané, Luiz Alcantara, Luiz Lehmann Coutinho, Maria Carolina Elias, Maurício Lacerda Nogueira, Rafael dos Santos Bezerra, Raul Machado Neto, Rejane Maria Tommasini Grotto, Ricardo Haddad, Sandra Coccuzzo Sampaio Vessoni, Simone Kashima, Svetoslav Nanev Slavov, Vincent Louis Viala |
| EPI_ISL_2445326 | UPA GEISEL | Instituto Butantan | Dimas Tadeu Covas, Antonio Jorge Martins, Claudia Renata dos Santos Barros, David Schlesinger, Debora Botequio Moretti, Elaine Cristina Marqueze, Elaine Vieira Santos, Evandra Strazza Rodrigues, Heidge Fukumasu, Jayme Augusto de Souza-Neto, José Salvatore Leister Patané, Luiz Alcantara, Luiz Lehmann Coutinho, Maria Carolina Elias, Maurício Lacerda Nogueira, Rafael dos Santos Bezerra, Raul Machado Neto, Rejane Maria Tommasini Grotto, Ricardo Haddad, Sandra Coccuzzo Sampaio Vessoni, Simone Kashima, Svetoslav Nanev Slavov, Vincent Louis Viala |
| EPI_ISL_2445406 | CS II DR MIGUEL VITALIANO ORLANDIA | Instituto Butantan | Dimas Tadeu Covas, Antonio Jorge Martins, Claudia Renata dos Santos Barros, David Schlesinger, Debora Botequio Moretti, Elaine Cristina Marqueze, Elaine Vieira Santos, Evandra Strazza Rodrigues, Heidge Fukumasu, Jayme Augusto de Souza-Neto, José Salvatore Leister Patané, Luiz Alcantara, Luiz Lehmann Coutinho, Maria Carolina Elias, Maurício Lacerda Nogueira, Rafael dos Santos Bezerra, Raul Machado Neto, Rejane Maria Tommasini Grotto, Ricardo Haddad, Sandra Coccuzzo Sampaio Vessoni, Simone Kashima, Svetoslav Nanev Slavov, Vincent Louis Viala |
| EPI_ISL_2445499 | UNIDADE DE SAUDE DR PHEBO DE OLIVEIRA ROGE FERREIRA | Instituto Butantan | Dimas Tadeu Covas, Antonio Jorge Martins, Claudia Renata dos Santos Barros, David Schlesinger, Debora Botequio Moretti, Elaine Cristina Marqueze, Elaine Vieira Santos, Evandra Strazza Rodrigues, Heidge Fukumasu, Jayme Augusto de Souza-Neto, José Salvatore Leister Patané, Luiz Alcantara, Luiz Lehmann Coutinho, Maria Carolina Elias, Maurício Lacerda Nogueira, Rafael dos Santos Bezerra, Raul Machado Neto, Rejane Maria Tommasini Grotto, Ricardo Haddad, Sandra Coccuzzo Sampaio Vessoni, Simone Kashima, Svetoslav Nanev Slavov, Vincent Louis Viala |
| EPI_ISL_2445564, EPI_ISL_2445570 | IRMANDADE DA SANTA CASA DE MISERICORDIA LORENA | Instituto Butantan | Dimas Tadeu Covas, Antonio Jorge Martins, Claudia Renata dos Santos Barros, David Schlesinger, Debora Botequio Moretti, Elaine Cristina Marqueze, Elaine Vieira Santos, Evandra Strazza Rodrigues, Heidge Fukumasu, Jayme Augusto de Souza-Neto, José Salvatore Leister Patané, Luiz Alcantara, Luiz Lehmann Coutinho, Maria Carolina Elias, Maurício Lacerda Nogueira, Rafael dos Santos Bezerra, Raul Machado Neto, Rejane Maria Tommasini Grotto, Ricardo Haddad, Sandra Coccuzzo Sampaio Vessoni, Simone Kashima, Svetoslav Nanev Slavov, Vincent Louis Viala |
| EPI_ISL_2445581 | HOSPITAL DE CAMPANHA COVID 19 MUNICIPIO DE TAUBATE | Instituto Butantan | Dimas Tadeu Covas, Antonio Jorge Martins, Claudia Renata dos Santos Barros, David Schlesinger, Debora Botequio Moretti, Elaine Cristina Marqueze, Elaine Vieira Santos, Evandra Strazza Rodrigues, Heidge Fukumasu, Jayme Augusto de Souza-Neto, José Salvatore Leister Patané, Luiz Alcantara, Luiz Lehmann Coutinho, Maria Carolina Elias, Maurício Lacerda Nogueira, Rafael dos Santos Bezerra, Raul Machado Neto, Rejane Maria Tommasini Grotto, Ricardo Haddad, Sandra Coccuzzo Sampaio Vessoni, Simone Kashima, Svetoslav Nanev Slavov, Vincent Louis Viala |
| EPI_ISL_2466257 | Laboratorio Central de Saude Publica do Estado de Alagoas (LACEN/AL) | Laboratory of Respiratory Viruses and Measles, Oswaldo Cruz Institute, FIOCRUZ | Paola Resende, Luciana Appolinario, Fernando Motta, Anna Carolina Paixao, Ana Carolina Mendonca, Alice Sampaio Rocha, Taina Venas, Elisa Cavalcante Pereira, Renata Serrano Lopes, Anderson Brandao Leite, Marilda Siqueira on behalf of the Fiocruz COVID-19 Genomic Surveillance Network |
| EPI_ISL_2466264 | Laboratorio Central de Saude Publica do Estado da Bahia (LACEN/BA) | Laboratory of Respiratory Viruses and Measles, Oswaldo Cruz Institute, FIOCRUZ | Paola Resende, Luciana Appolinario, Fernando Motta, Anna Carolina Paixao, Ana Carolina Mendonca, Alice Sampaio Rocha, Taina Venas, Elisa Cavalcante Pereira, Renata Serrano Lopes,Felicidade Pereira, Marilda Siqueira on behalf of the Fiocruz COVID-19 Genomic Surveillance Network |
| EPI_ISL_2466447, EPI_ISL_2466459 | HLAGYN - Laboratorio de Imunologia de Transplantes de Goias | HLAGYN - Laboratorio de Imunologia de Transplantes de Goias | Fernando Antonio Vinhal dos Santos, Erika Lopes Rocha Batista, Alessandro Leonardo Alvares Magalhaes, Frederico Rodrigues Vinhal, Sabrina Sara Moreira Duarte, Lucas Carlos Gomes Pereira, Daniel Ferreira de Sousa |
| EPI_ISL_2473714 | UBS PASTOR MARCELINO DEUNGARO RIO GRANDE | Instituto Butantan | Dimas Tadeu Covas, Antonio Jorge Martins, Claudia Renata dos Santos Barros, David Schlesinger, Debora Botequio Moretti, Elaine Cristina Marqueze, Elaine Vieira Santos, Evandra Strazza Rodrigues, Heidge Fukumasu, Jayme Augusto de Souza-Neto, José Salvatore Leister Patané, Luiz Alcantara, Luiz Lehmann Coutinho, Maria Carolina Elias, Maurício Lacerda Nogueira, Rafael dos Santos Bezerra, Raul Machado Neto, Rejane Maria Tommasini Grotto, Ricardo Haddad, Sandra Coccuzzo Sampaio Vessoni, Simone Kashima, Svetoslav Nanev Slavov, Vincent Louis Viala |
| EPI_ISL_2473727, EPI_ISL_2473734 | HOSP GERAL DE SAO PAULO | Instituto Butantan | Dimas Tadeu Covas, Antonio Jorge Martins, Claudia Renata dos Santos Barros, David Schlesinger, Debora Botequio Moretti, Elaine Cristina Marqueze, Elaine Vieira Santos, Evandra Strazza Rodrigues, Heidge Fukumasu, Jayme Augusto de Souza-Neto, José Salvatore Leister Patané, Luiz Alcantara, Luiz Lehmann Coutinho, Maria Carolina Elias, Maurício Lacerda Nogueira, Rafael dos Santos Bezerra, Raul Machado Neto, Rejane Maria Tommasini Grotto, Ricardo Haddad, Sandra Coccuzzo Sampaio Vessoni, Simone Kashima, Svetoslav Nanev Slavov, Vincent Louis Viala |
| EPI_ISL_2473740 | NUCLEO DE SAUDE VILA FALCAO DE BAURU | Instituto Butantan | Dimas Tadeu Covas, Antonio Jorge Martins, Claudia Renata dos Santos Barros, David Schlesinger, Debora Botequio Moretti, Elaine Cristina Marqueze, Elaine Vieira Santos, Evandra Strazza Rodrigues, Heidge Fukumasu, Jayme Augusto de Souza-Neto, José Salvatore Leister Patané, Luiz Alcantara, Luiz Lehmann Coutinho, Maria Carolina Elias, Maurício Lacerda Nogueira, Rafael dos Santos Bezerra, Raul Machado Neto, Rejane Maria Tommasini Grotto, Ricardo Haddad, Sandra Coccuzzo Sampaio Vessoni, Simone Kashima, Svetoslav Nanev Slavov, Vincent Louis Viala |
| EPI_ISL_2473786 | CADIP VILA REGINA FERNANDOPOLIS | Instituto Butantan | Dimas Tadeu Covas, Antonio Jorge Martins, Claudia Renata dos Santos Barros, David Schlesinger, Debora Botequio Moretti, Elaine Cristina Marqueze, Elaine Vieira Santos, Evandra Strazza Rodrigues, Heidge Fukumasu, Jayme Augusto de Souza-Neto, José Salvatore Leister Patané, Luiz Alcantara, Luiz Lehmann Coutinho, Maria Carolina Elias, Maurício Lacerda Nogueira, Rafael dos Santos Bezerra, Raul Machado Neto, Rejane Maria Tommasini Grotto, Ricardo Haddad, Sandra Coccuzzo Sampaio Vessoni, Simone Kashima, Svetoslav Nanev Slavov, Vincent Louis Viala |
| EPI_ISL_2493371 | SECRETARIA DE SAUDE DE SAO PEDRO | Instituto Butantan | Dimas Tadeu Covas, Antonio Jorge Martins, Claudia Renata dos Santos Barros, David Schlesinger, Debora Botequio Moretti, Elaine Cristina Marqueze, Elaine Vieira Santos, Evandra Strazza Rodrigues, Heidge Fukumasu, Jayme Augusto de Souza-Neto, José Salvatore Leister Patané, Luiz Alcantara, Luiz Lehmann Coutinho, Maria Carolina Elias, Maurício Lacerda Nogueira, Rafael dos Santos Bezerra, Raul Machado Neto, Rejane Maria Tommasini Grotto, Ricardo Haddad, Sandra Coccuzzo Sampaio Vessoni, Simone Kashima, Svetoslav Nanev Slavov, Vincent Louis Viala |
| EPI_ISL_2493383 | UNIDADE BASICA DE SAUDE DR MATHEUS GABRIEL BONASSA | Instituto Butantan | Dimas Tadeu Covas, Antonio Jorge Martins, Claudia Renata dos Santos Barros, David Schlesinger, Debora Botequio Moretti, Elaine Cristina Marqueze, Elaine Vieira Santos, Evandra Strazza Rodrigues, Heidge Fukumasu, Jayme Augusto de Souza-Neto, José Salvatore Leister Patané, Luiz Alcantara, Luiz Lehmann Coutinho, Maria Carolina Elias, Maurício Lacerda Nogueira, Rafael dos Santos Bezerra, Raul Machado Neto, Rejane Maria Tommasini Grotto, Ricardo Haddad, Sandra Coccuzzo Sampaio Vessoni, Simone Kashima, Svetoslav Nanev Slavov, Vincent Louis Viala |
| EPI_ISL_2493388, EPI_ISL_2493444 | SECRETARIA DE SAUDE DE SAO PEDRO | Instituto Butantan | Dimas Tadeu Covas, Antonio Jorge Martins, Claudia Renata dos Santos Barros, David Schlesinger, Debora Botequio Moretti, Elaine Cristina Marqueze, Elaine Vieira Santos, Evandra Strazza Rodrigues, Heidge Fukumasu, Jayme Augusto de Souza-Neto, José Salvatore Leister Patané, Luiz Alcantara, Luiz Lehmann Coutinho, Maria Carolina Elias, Maurício Lacerda Nogueira, Rafael dos Santos Bezerra, Raul Machado Neto, Rejane Maria Tommasini Grotto, Ricardo Haddad, Sandra Coccuzzo Sampaio Vessoni, Simone Kashima, Svetoslav Nanev Slavov, Vincent Louis Viala |
| EPI_ISL_2493697 | SECRETARIA MUNICIPAL DE SAUDE SOROCABA | Instituto Butantan | Dimas Tadeu Covas, Antonio Jorge Martins, Claudia Renata dos Santos Barros, David Schlesinger, Debora Botequio Moretti, Elaine Cristina Marqueze, Elaine Vieira Santos, Evandra Strazza Rodrigues, Heidge Fukumasu, Jayme Augusto de Souza-Neto, José Salvatore Leister Patané, Luiz Alcantara, Luiz Lehmann Coutinho, Maria Carolina Elias, Maurício Lacerda Nogueira, Rafael dos Santos Bezerra, Raul Machado Neto, Rejane Maria Tommasini Grotto, Ricardo Haddad, Sandra Coccuzzo Sampaio Vessoni, Simone Kashima, Svetoslav Nanev Slavov, Vincent Louis Viala |
| EPI_ISL_2493750 | AFIP | Instituto Butantan | Dimas Tadeu Covas, Antonio Jorge Martins, Claudia Renata dos Santos Barros, David Schlesinger, Debora Botequio Moretti, Elaine Cristina Marqueze, Elaine Vieira Santos, Evandra Strazza Rodrigues, Heidge Fukumasu, Jayme Augusto de Souza-Neto, José Salvatore Leister Patané, Luiz Alcantara, Luiz Lehmann Coutinho, Maria Carolina Elias, Maurício Lacerda Nogueira, Rafael dos Santos Bezerra, Raul Machado Neto, Rejane Maria Tommasini Grotto, Ricardo Haddad, Sandra Coccuzzo Sampaio Vessoni, Simone Kashima, Svetoslav Nanev Slavov, Vincent Louis Viala |
| EPI_ISL_2493820 | UBS JARDIM CAIUBY | Instituto Butantan | Dimas Tadeu Covas, Antonio Jorge Martins, Claudia Renata dos Santos Barros, David Schlesinger, Debora Botequio Moretti, Elaine Cristina Marqueze, Elaine Vieira Santos, Evandra Strazza Rodrigues, Heidge Fukumasu, Jayme Augusto de Souza-Neto, José Salvatore Leister Patané, Luiz Alcantara, Luiz Lehmann Coutinho, Maria Carolina Elias, Maurício Lacerda Nogueira, Rafael dos Santos Bezerra, Raul Machado Neto, Rejane Maria Tommasini Grotto, Ricardo Haddad, Sandra Coccuzzo Sampaio Vessoni, Simone Kashima, Svetoslav Nanev Slavov, Vincent Louis Viala |
| EPI_ISL_2493831 | USAFA SAO JORGE | Instituto Butantan | Dimas Tadeu Covas, Antonio Jorge Martins, Claudia Renata dos Santos Barros, David Schlesinger, Debora Botequio Moretti, Elaine Cristina Marqueze, Elaine Vieira Santos, Evandra Strazza Rodrigues, Heidge Fukumasu, Jayme Augusto de Souza-Neto, José Salvatore Leister Patané, Luiz Alcantara, Luiz Lehmann Coutinho, Maria Carolina Elias, Maurício Lacerda Nogueira, Rafael dos Santos Bezerra, Raul Machado Neto, Rejane Maria Tommasini Grotto, Ricardo Haddad, Sandra Coccuzzo Sampaio Vessoni, Simone Kashima, Svetoslav Nanev Slavov, Vincent Louis Viala |
| EPI_ISL_2493891 | PRONTO ATENDIMENTO DE VARGEM GRANDE PAULISTA | Instituto Butantan | Dimas Tadeu Covas, Antonio Jorge Martins, Claudia Renata dos Santos Barros, David Schlesinger, Debora Botequio Moretti, Elaine Cristina Marqueze, Elaine Vieira Santos, Evandra Strazza Rodrigues, Heidge Fukumasu, Jayme Augusto de Souza-Neto, José Salvatore Leister Patané, Luiz Alcantara, Luiz Lehmann Coutinho, Maria Carolina Elias, Maurício Lacerda Nogueira, Rafael dos Santos Bezerra, Raul Machado Neto, Rejane Maria Tommasini Grotto, |

| | | | |
|---|---|---|---|
| | | | Ricardo Haddad, Sandra Coccuzzo Sampaio Vessoni, Simone Kashima, Svetoslav Nanev Slavov, Vincent Louis Viala |
| EPI_ISL_2494003 | INSIDE CRSN PIRITUBA | Instituto Butantan | Dimas Tadeu Covas, Antonio Jorge Martins, Claudia Renata dos Santos Barros, David Schlesinger, Debora Botequio Moretti, Elaine Cristina Marqueze, Elaine Vieira Santos, Evandra Strazza Rodrigues, Heidge Fukumasu, Jayme Augusto de Souza-Neto, José Salvatore Leister Patané, Luiz Alcantara, Luiz Lehmann Coutinho, Maria Carolina Elias, Maurício Lacerda Nogueira, Rafael dos Santos Bezerra, Raul Machado Neto, Rejane Maria Tommasini Grotto, Ricardo Haddad, Sandra Coccuzzo Sampaio Vessoni, Simone Kashima, Svetoslav Nanev Slavov, Vincent Louis Viala |
| EPI_ISL_2494025 | INSIDE CRSSU IPIRANGA | Instituto Butantan | Dimas Tadeu Covas, Antonio Jorge Martins, Claudia Renata dos Santos Barros, David Schlesinger, Debora Botequio Moretti, Elaine Cristina Marqueze, Elaine Vieira Santos, Evandra Strazza Rodrigues, Heidge Fukumasu, Jayme Augusto de Souza-Neto, José Salvatore Leister Patané, Luiz Alcantara, Luiz Lehmann Coutinho, Maria Carolina Elias, Maurício Lacerda Nogueira, Rafael dos Santos Bezerra, Raul Machado Neto, Rejane Maria Tommasini Grotto, Ricardo Haddad, Sandra Coccuzzo Sampaio Vessoni, Simone Kashima, Svetoslav Nanev Slavov, Vincent Louis Viala |
| EPI_ISL_2494091 | AFIP SUL | Instituto Butantan | Dimas Tadeu Covas, Antonio Jorge Martins, Claudia Renata dos Santos Barros, David Schlesinger, Debora Botequio Moretti, Elaine Cristina Marqueze, Elaine Vieira Santos, Evandra Strazza Rodrigues, Heidge Fukumasu, Jayme Augusto de Souza-Neto, José Salvatore Leister Patané, Luiz Alcantara, Luiz Lehmann Coutinho, Maria Carolina Elias, Maurício Lacerda Nogueira, Rafael dos Santos Bezerra, Raul Machado Neto, Rejane Maria Tommasini Grotto, Ricardo Haddad, Sandra Coccuzzo Sampaio Vessoni, Simone Kashima, Svetoslav Nanev Slavov, Vincent Louis Viala |
| EPI_ISL_2494197 | VIGILANCIA EPIDEMIOLOGICA E CONTROLE DE VETORES PIRASSUNUN | Instituto Butantan | Dimas Tadeu Covas, Antonio Jorge Martins, Claudia Renata dos Santos Barros, David Schlesinger, Debora Botequio Moretti, Elaine Cristina Marqueze, Elaine Vieira Santos, Evandra Strazza Rodrigues, Heidge Fukumasu, Jayme Augusto de Souza-Neto, José Salvatore Leister Patané, Luiz Alcantara, Luiz Lehmann Coutinho, Maria Carolina Elias, Maurício Lacerda Nogueira, Rafael dos Santos Bezerra, Raul Machado Neto, Rejane Maria Tommasini Grotto, Ricardo Haddad, Sandra Coccuzzo Sampaio Vessoni, Simone Kashima, Svetoslav Nanev Slavov, Vincent Louis Viala |
| EPI_ISL_2494225 | BIOFAST LESTE | Instituto Butantan | Dimas Tadeu Covas, Antonio Jorge Martins, Claudia Renata dos Santos Barros, David Schlesinger, Debora Botequio Moretti, Elaine Cristina Marqueze, Elaine Vieira Santos, Evandra Strazza Rodrigues, Heidge Fukumasu, Jayme Augusto de Souza-Neto, José Salvatore Leister Patané, Luiz Alcantara, Luiz Lehmann Coutinho, Maria Carolina Elias, Maurício Lacerda Nogueira, Rafael dos Santos Bezerra, Raul Machado Neto, Rejane Maria Tommasini Grotto, Ricardo Haddad, Sandra Coccuzzo Sampaio Vessoni, Simone Kashima, Svetoslav Nanev Slavov, Vincent Louis Viala |
| EPI_ISL_2494291 | CENTRO MEDICO DR NELSON SALOME DE CONCHAL | Instituto Butantan | Dimas Tadeu Covas, Antonio Jorge Martins, Claudia Renata dos Santos Barros, David Schlesinger, Debora Botequio Moretti, Elaine Cristina Marqueze, Elaine Vieira Santos, Evandra Strazza Rodrigues, Heidge Fukumasu, Jayme Augusto de Souza-Neto, José Salvatore Leister Patané, Luiz Alcantara, Luiz Lehmann Coutinho, Maria Carolina Elias, Maurício Lacerda Nogueira, Rafael dos Santos Bezerra, Raul Machado Neto, Rejane Maria Tommasini Grotto, Ricardo Haddad, Sandra Coccuzzo Sampaio Vessoni, Simone Kashima, Svetoslav Nanev Slavov, Vincent Louis Viala |
| EPI_ISL_2497447, EPI_ISL_2497463 | HLAGYN - Laboratorio de Imunologia de Transplantes de Goias | HLAGYN - Laboratorio de Imunologia de Transplantes de Goias | Fernando Antonio Vinhal dos Santos, Erika Lopes Rocha Batista, Alessandro Leonardo Alvares Magalhaes, Frederico Rodrigues Vinhal, Sabrina Sara Moreira Duarte, Lucas Carlos Gomes Pereira, Daniel Ferreira de Sousa |
| EPI_ISL_2534864, EPI_ISL_2534882, EPI_ISL_2534926, EPI_ISL_2534940, EPI_ISL_2534958, EPI_ISL_2534969, EPI_ISL_2534970, EPI_ISL_2534972, EPI_ISL_2535013, EPI_ISL_2535019, EPI_ISL_2535026, EPI_ISL_2535057 | | | |
| see above | Laboratorio Central Noel Nutels | Bioinformatics Laboratory / LNCC | Luiz G P de Almeida, Alessandra P Lamarca, Ronaldo da Silva F Jr, Liliane Cavalcante, Alexandra L Gerber, Ana Paula de C Guimaraes, Douglas Terra Machado, Cassia Alves, Diana Mariani, Cintia Policarpo, Gleidson da Silva de Oliveira, Mario Sergio Ribeiro, Silvia Carvalho, Flavio Dias da Silva, Marcio Henrique de Oliveira Garcia, Leandro Magalhaes de Souza, Cristiane Gomes da Silva, Caio Luiz Pereira Ribeiro, Andrea Cony Cavalcanti, Claudia Maria Braga de Mello, Amilcar Tanuri, Ana Tereza R Vasconcelos |
| EPI_ISL_2535149, EPI_ISL_2535219, EPI_ISL_2535231 | Unidade de apoio ao diagnostico da COVID - UNADIG | Bioinformatics Laboratory / LNCC | Luiz G P de Almeida, Alessandra P Lamarca, Ronaldo da Silva F Jr, Liliane Cavalcante, Alexandra L Gerber, Ana Paula de C Guimaraes, Douglas Terra Machado, Cassia Alves, Diana Mariani, Cintia Policarpo, Gleidson da Silva de Oliveira, Mario Sergio Ribeiro, Silvia Carvalho, Flavio Dias da Silva, Marcio Henrique de Oliveira Garcia, Leandro Magalhaes de Souza, Cristiane Gomes da Silva, Caio Luiz Pereira Ribeiro, Andrea Cony Cavalcanti, Claudia Maria Braga de Mello, Amilcar Tanuri, Ana Tereza R Vasconcelos |
| EPI_ISL_2536228 | Labortorio Central de Saude Publica do Estado de Santa Catarina (LACEN/SC) | Laboratory of Respiratory Viruses and Measles, Oswaldo Cruz Institute, FIOCRUZ | Paola Resende, Luciana Appolinario, Fernando Motta, Anna Carolina Paixao, Ana Carolina Mendonca, Alice Sampaio Rocha, Taina Venas, Elisa Cavalcante Pereira, Renata Serrano Lopes, Darcita Buerger Rovaris, Sandra Bianchini Fernandes, Marilda Siqueira on behalf of the Fiocruz COVID-19 Genomic Surveillance Network |
| EPI_ISL_2603449, EPI_ISL_2603539, EPI_ISL_2603549, EPI_ISL_2603558, EPI_ISL_2603584, EPI_ISL_2603585 | Laboratorio Central de Saude Publica do Estado do Rio Grande do Sul (LACEN-RS) | Laboratory of Respiratory Viruses and Measles, Oswaldo Cruz Institute, FIOCRUZ | Paola Resende, Luciana Appolinario, Fernando Motta, Anna Carolina Paixao, Ana Carolina Mendonca, Alice Sampaio Rocha, Taina Venas, Elisa Cavalcante Pereira, Renata Serrano Lopes, Anderson Brandao Leite, Marilda Siqueira on behalf of the Fiocruz COVID-19 Genomic Surveillance Network |
| EPI_ISL_2614319 | Laboratory of Respiratory Viruses and Measles, Oswaldo Cruz Institute, FIOCRUZ | Laboratory of Respiratory Viruses and Measles, Oswaldo Cruz Institute, FIOCRUZ | Paola Resende, Luciana Appolinario, Fernando Motta, Anna Carolina Paixao, Ana Carolina Mendonca, Alice Sampaio Rocha, Taina Venas, Elisa Cavalcante Pereira, Renata Serrano Lopes, Marilda Siqueira on behalf of the Fiocruz COVID-19 Genomic Surveillance Network |
| EPI_ISL_2617903, EPI_ISL_2617906, EPI_ISL_2617919 | LABCOVID_HCPA | LABRESIS_HCPA | Wink PL, Martins AF, Volpato F, Monteiro F, Zavascki AP, Barth AL |
| EPI_ISL_2629762 | Laboratório de Virologia Molecular - Universidade Federal do Rio de Janeiro | Laboratório de Virologia Molecular - Universidade Federal do Rio de Janeiro | Filipe Romero Rebello Moreira, Mirela D'arc, Diana Mariani, Alice Laschuk Herlinger, Francine Bittencourt Schiffler, Átila Duque Rossi, Isabela de Carvalho Leitão, Thamiris dos Santos Miranda, Matheus Augusto Calvano Cosentino, Marcelo Calado de Paula Tôrres, Raíssa Mirella dos Santos Cunha da Costa, Cássia Cristina Alves Gonçalves, Débora Souza Faffe, Rafael Mello Galliez, Orlando da Costa Ferreira Junior, Renato Santana de Aguiar,, André Felipe Andrade dos Santos, Carolina Moreira Voloch, Terezinha Marta Pereira Pinto Castineiras, Amilcar Tanuri |
| EPI_ISL_2645622 | Laboratorio Central de Saude Publica do Estado do Espirito Santo (LACEN/ES) | Laboratory of Respiratory Viruses and Measles, Oswaldo Cruz Institute, FIOCRUZ | Paola Resende, Luciana Appolinario, Fernando Motta, Anna Carolina Paixao, Ana Carolina Mendonca, Alice Sampaio Rocha, Taina Venas, Eilisa Cavalcante Pereira, Renata Serrano Lopes, Rodrigo Ribeiro Rodrigues, Marilda Siqueira on behalf of the Fiocruz COVID-19 Genomic Surveillance Network |
| EPI_ISL_2645910, EPI_ISL_2645913, EPI_ISL_2645914 | Laboratorio Central de Saude Publica do Estado do Para (LACEN/PA) | Laboratory of Respiratory Viruses and Measles, Oswaldo Cruz Institute, FIOCRUZ | Paola Resende, Luciana Appolinario, Fernando Motta, Anna Carolina Paixao, Ana Carolina Mendonca, Alice Sampaio Rocha, Taina Venas, Elisa Cavalcante Pereira, Renata Serrano Lopes, Valnete Andrade, Marilda Siqueira on behalf of the Fiocruz COVID-19 Genomic Surveillance Network |
| EPI_ISL_2660571, EPI_ISL_2660572, EPI_ISL_2660574, EPI_ISL_2660575, EPI_ISL_2660579, EPI_ISL_2660582, EPI_ISL_2660585, EPI_ISL_2660586, EPI_ISL_2660590, EPI_ISL_2660591, EPI_ISL_2660592, EPI_ISL_2660593, EPI_ISL_2660597, EPI_ISL_2660598, EPI_ISL_2660606, EPI_ISL_2660607, EPI_ISL_2660608, EPI_ISL_2660622, EPI_ISL_2660624, EPI_ISL_2660634, EPI_ISL_2660635, EPI_ISL_2660645, EPI_ISL_2660649, EPI_ISL_2660654, EPI_ISL_2660657, EPI_ISL_2660658, EPI_ISL_2660662 | | | |
| see above | Laboratorio Central de Saude Publica do Estado de Sergipe (LACEN/SE) | Laboratory of Respiratory Viruses and Measles, Oswaldo Cruz Institute, FIOCRUZ | Paola Resende, Luciana Appolinario, Fernando Motta, Anna Carolina Paixao, Ana Carolina Mendonca, Alice Sampaio Rocha, Tainá Moreira Martins Venas, Elisa Cavalcante Pereira, Renata Serrano Lopes, Cliomar Alves dos Santos, Marilda Siqueira on behalf of the Fiocruz COVID-19 Genomic Surveillance Network |
| EPI_ISL_2661764, EPI_ISL_2661766, EPI_ISL_2661779, EPI_ISL_2661824, EPI_ISL_2661837, EPI_ISL_2661844, EPI_ISL_2661845, EPI_ISL_2661847, EPI_ISL_2661854, EPI_ISL_2661855, EPI_ISL_2661856, EPI_ISL_2661857, EPI_ISL_2661859 | | | |
| see above | Laboratorio Central de Saude Publica do Estado do Rio Grande do Sul (LACEN-RS) | Laboratory of Respiratory Viruses and Measles, Oswaldo Cruz Institute, FIOCRUZ | Paola Resende, Luciana Appolinario, Fernando Motta, Anna Carolina Paixao, Ana Carolina Mendonca, Alice Sampaio Rocha, Taina Venas, Elisa Cavalcante Pereira, Renata Serrano Lopes, Tatiana Schaffer Gregianini, Richard Salvato, Marilda Siqueira on behalf of the Fiocruz COVID-19 Genomic Surveillance Network |
| EPI_ISL_2663277 | Plataforma de Vigilancia Molecular (PVM) - FIOCRUZ/BA | Plataforma de Vigilancia Molecular (PVM) - FIOCRUZ/BA | Ricardo Khouri, Marina Cucco, Tiago Graf, Clarissa Araújo Gurgel, Leonardo Paiva Farias, Bruno Bezerril Andrade, Camila I. de Oliveira on behalf of the Fiocruz COVID-19 Genomic Surveillance Network. |
| EPI_ISL_2691113, EPI_ISL_2691119 | Instituto Adolfo Lutz - Regional de Bauru | Instituto Adolfo Lutz, Interdiciplinary Procedures Center, Strategic Laboratory | Claudio Tavares Sacchi, Claudia Regina Gonçalves, Erica Valessa Ramos Gomes, Karoline Rodrigues Campos, Caio Vinicius Dias Lopes, Leonardo Jose Tadeu de Araujo |
| EPI_ISL_2691342, EPI_ISL_2691346, EPI_ISL_2691365, EPI_ISL_2691405, EPI_ISL_2691406, EPI_ISL_2691407, EPI_ISL_2691408, EPI_ISL_2691413, EPI_ISL_2691423, EPI_ISL_2691429, EPI_ISL_2691452, EPI_ISL_2691542 | | | |
| see above | Laboratorio Central Noel Nutels | Bioinformatics Laboratory / LNCC | Luiz G P de Almeida, Alessandra P Lamarca, Ronaldo da Silva F Jr, Liliane Cavalcante, Alexandra L Gerber, Ana Paula de C Guimaraes, Douglas Terra Machado, Cassia Alves, Diana Mariani, Cintia Policarpo, Gleidson da Silva de Oliveira, Mario Sergio Ribeiro, Silvia Carvalho, Flavio Dias da Silva, Marcio Henrique de Oliveira Garcia, Leandro Magalhaes de Souza, Cristiane Gomes da Silva, Caio Luiz Pereira Ribeiro, Andrea Cony Cavalcanti, Claudia Maria Braga de Mello, Amilcar Tanuri, Ana Tereza R Vasconcelos |
| EPI_ISL_2691588, EPI_ISL_2691589 | Unidade de apoio ao diagnostico da COVID - UNADIG | Bioinformatics Laboratory / LNCC | Luiz G P de Almeida, Alessandra P Lamarca, Ronaldo da Silva F Jr, Liliane Cavalcante, Alexandra L Gerber, Ana Paula de C Guimaraes, Douglas Terra |

| | | | |
|---|---|---|---|
| | | | Machado, Cassia Alves, Diana Mariani, Cintia Policarpo, Gleidson da Silva de Oliveira, Mario Sergio Ribeiro, Silvia Carvalho, Flavio Dias da Silva, Marcio Henrique de Oliveira Garcia, Leandro Magalhaes de Souza, Cristiane Gomes da Silva, Caio Luiz Pereira Ribeiro, Andrea Cony Cavalcanti, Claudia Maria Braga de Mello, Amilcar Tanuri, Ana Tereza R Vasconcelos |
| EPI_ISL_2731541 | Laboratório de Biotecnologia Aplicada (LBA) - Laboratório de Biologia Molecular - Hospital das Clínicas, Faculdade de Medicina de Botucatu, Departamento de Bioprocessos e Biotecnologia - Faculdade de Ciências Agronômicas, UNESP - Botucatu/SP | Laboratory of Respiratory Viruses and Measles, Oswaldo Cruz Institute, FIOCRUZ | Paola Resende, Jayme Augusto de Souza Neto, Rejane Maria Tommasini, Leonardo Nazario de Moraes, Felipe Allan da Silva Costa, Patricia Akemi Assato, Luciana Appolinario, Fernando Motta, Anna Carolina Paixao, Ana Carolina Mendonca, Alice Sampaio Rocha, Taina Venas, Elisa Cavalcante Pereira, Renata Serrano Lopes, Marilda Siqueira on behalf of the Fiocruz COVID-19 Genomic Surveillance Network |
| EPI_ISL_983865 | Central Laboratory of Public Health of Rio Grande do Sul (Lacen-RS) | State Center for Health Surveillance of the Health Department of the State of Rio Grande do Sul (CEVS/SES-RS) | Aline Campos, Cynthia Molina, Lara Crescente, Leticia Garay, Ludmila Fiorenzano Baethgen, Richard Salvato, Tatiana Gregianini |
